# The Q-Warg Pipeline: A Robust and Versatile Workflow for Quantitative Analysis of Protoplast Culture Conditions

**DOI:** 10.1101/2025.03.05.641611

**Authors:** Léa Bogdziewiez, Rik Froeling, Patricia Schoppl, Jeanne Juquel, Ioanna Antoniadi, Vladimìr Skalický, Ambroise Mathey, Jacques Fattaccioli, Joris Sprakel, Stéphane Verger

## Abstract

Single cells offer a simplified model for investigating complex mechanisms such as cell-cell adhesion. Protoplasts, plant cells without cell walls (CWs), have been instrumental in plant research, industrial applications, and breeding. However, due to the absence of a CW, protoplasts are not considered “true” plant cells and making them less relevant for biophysical studies. Current protocols for CW recovery in protoplasts vary widely among laboratories and starting materials, requiring lab-specific optimizations that often depend on expert knowledge and qualitative assessments. To address this, we have developed a user-friendly streamlined workflow, the Q-Warg pipeline, which enables quantitative comparison of various conditions for CW recovery post-protoplasting. This pipeline employs fluorescence imaging and tailored processing to measure parameters such as morphometry, cell viability, and CW staining intensity. Using this approach, we optimized culture conditions to obtain single plant cells (SPCs) with recovered CWs. Additionally, we demonstrated the robustness and versatility of the workflow by quantifying different fluorescent signals in protoplast suspensions. Overall, the Q-Warg pipeline provides a widely available and user-friendly solution for robust and unbiased characterization of protoplasts culture. The quantitative data generated by the pipeline may be useful in the future to decipher the mechanisms regulating protoplast viability and regeneration.

**Significance statement:** Several fields of plant biology, ranging from biotechnology to biomechanics, have recently regained a strong interest in using and studying protoplasts and single cells. Here, we developed a widely accessible quantitative workflow to characterize cell culture recovery after protoplasting along with the demonstration of its usefulness and versatility in various cases. We hope this tool can help other research groups to streamline the procedure needed to establish single plant cell approaches in their lab.

## Introduction

Multicellular organisms are made of millions or trillions of cells coming together in complex tissues with heterogeneities at several scales. While these heterogeneities characterize the form and functions of those tissues, they often complicate our understanding of biological mechanisms occurring within them. For instance, studying the phenomenon of cell-cell adhesion in the tissue context is complex because tissue topology and cell geometry as well as cell-cell interface heterogeneities can have major confounding effects on the apparent strength of cell adhesion (Bidhendi and Geitmann, 2016). On the other hand, while often less physiologically relevant at the organism scale, working with single cells provide a strongly simplified framework to study cell-cell interaction (Atakhani, Bogdziewiez and Verger, 2022). Beyond the example of adhesion, such consideration applies to a large spectrum of biological questions which explains why cell cultures are such a widely used model system in animal sciences (Verma, Verma and Singh, 2020). However, when it comes to studying plants, “single plant cell cultures” do not really exist. It is possible to cultivate cells *in vitro* under the form of microcalli, which are small clusters of cells arising from division events and thus not real single cells (Pesquet, Wagner and Grabber, 2019; Krasteva, Georgiev and Pavlov, 2021). It is also possible to isolate and study protoplasts (see further details below; (Mukundan, Satyamoorthy and Babu, 2025)), which are effectively single cells but lack the cell wall, a crucial component of plant cells that controls their shape, mechanics, adhesion, growth and plays major roles in signalling, defence and cell-to-cell communication (Yokoyama *et al*., 2014; Anderson and Kieber, 2020; Cosgrove, 2024; Anderson and Pelloux, 2025). It would thus be particularly useful for the plant science community to establish an approach to generate single plant cell populations in order to use them as a new model system. One possibility is to start by extracting protoplasts and letting them recover their cell wall. The challenge lies in identifying culture conditions that promote strong and homogeneous cell wall recovery while ensuring minimal cell division and high cell survival rate over several days.

A protoplast is a plant cell without its cell wall (CW, term introduced by Hanstein in 1880). In 1892, the first isolation of protoplasts was successfully conducted by Klercker (*Eine Methode zur Isolierung lebender Protoplasten / von John Af Klercker*, 1892). In 1970, Nagata and Takabe, regenerated a full plant from tobacco leaves protoplasts (Nagata and Takebe, 1970). Protoplasts have proven their usefulness in plant research to study various cellular processes such as cell division, cell wall synthesis or cell differentiation (Jiang, Zhu and Liu, 2013; Yokoyama *et al*., 2016; Pasternak *et al*., 2020; Gilliard *et al*., 2021) but also in industry for metabolites production (Xu *et al*., 2022). They are also employed as a breeding technology since a whole plant can be regenerated from genetically transformed protoplasts (Reed and Bargmann, 2021; Reyna-Llorens, Ferro-Costa and Burgess, 2023; Mukundan, Satyamoorthy and Babu, 2025). Furthermore, recent development of non-transgenic genome editing with CRISPR/Cas9 and ribonucleoproteins (Yue *et al*., 2021; Laforest and Nadakuduti, 2022) has renewed the interest in protoplasts.

Despite more than a century of research, it is still challenging to work with protoplasts as exemplified by the large number of protocols for protoplast extraction and tissue/plant regeneration that exist (Table S2). The first step of extracting the protoplasts from a plant tissue is relatively straight forward and has been done from various plant species (Kumari, 2019). Most protocols use enzymatic digestion of the CW to release protoplasts. As the CW composition can vary from one tissue to another, enzymes must be carefully selected (Mukundan, Satyamoorthy and Babu, 2025). Once the protoplasts are isolated, the first quest is to find the right conditions to keep them alive. Without their CW, protoplasts are fragile and cannot withstand physical/mechanical stress. They require high osmolarity and low to no agitation to avoid bursting before the cell wall is recovered. However, the following steps leading to CW recovery and cell division, toward tissue regeneration appear to be the most challenging to establish and optimize (Xu *et al*., 2022). Plant cells are normally surrounded by their CW which is mainly composed of cellulose and a matrix of other polysaccharides deposited outside of the plasma membrane (Cosgrove, 2024). Likely the first challenge for the protoplasts after the CW removal is to retain newly synthesized cell wall polysaccharides at their surface. Matrix polysaccharides are secreted from the Golgi and may at first simply end up in the liquid medium (Oda *et al*., 2020; Hoffmann *et al*., 2021). Similarly, cellulose is synthesized at the surface of the plasma membrane and may primarily be extruded outward rather than sticking to the membrane (Paredez, Somerville and Ehrhardt, 2006; Purushotham, Ho and Zimmer, 2020). Yet, those cellulose microfibrils are tethered to the membrane via the cellulose synthase complex and may act as traps for secreted polysaccharides such as xyloglucans (Park and Cosgrove, 2012). It is also possible that Golgi-synthesized polysaccharides already interact with membrane bound protein before secretion and thus remain partially attached to the surface upon secretion (McKenna, Tolmie and Runions, 2014; Fruleux, Verger and Boudaoud, 2019). Furthermore, glycosylphosphatidylinositol (GPI) -anchored arabinogalactan proteins, with their extensive glycosylation and anchorage at the membrane, may be one of the players contributing to retain polysaccharides at the membrane surface (Tan *et al*., 2013; Leszczuk *et al*., 2023). Over time the cell wall polysaccharides may begin to accumulate in patches, while the cellulose could develop into a network which will then efficiently retain additional CW deposits on the cell surface (Tagawa *et al*., 2019). Ultimately a rather uniform CW may be recovered around the protoplast (Tagawa *et al*., 2019). However, this scenario remains very speculative as the mechanisms of CW recovery are still largely unknown. Although it seems that cell wall recovery would be almost inevitable over time, in reality, depending on the growth conditions, many protoplasts—often the majority—are unable to recover their CWs, and instead remain as seemingly wall-less protoplasts (Xu *et al*., 2022). Some protocols aiming to regenerate whole plants from protoplasts take the approach of embedding the protoplasts into a gel (alginate) to help the CW recovery and callus formation (Damm and Willmitzer, 1988; Masson and Paszkowski, 1992; Jeong *et al*., 2021; Sakamoto *et al*., 2022). This embedding likely allows the full retention of secreted polysaccharides at the protoplast surface and thus an efficient recovery of the CW, opening the way for the first cell division, microcalli formation and later on tissue regeneration. While CW recovery is a crucial step in the process, for applications aiming towards tissue regeneration, it is generally enough that small proportions of the protoplasts recover their CW (Jeong *et al*., 2021). These cells can then divide, proliferate, and ultimately constitute the majority of the cell culture over the protoplasts that didn’t recover their CW. It is likely for those reasons, along with protoplast embedding methods, that optimizing protoplast CW generation has not been a major research focus in the past. Yet, with the renewed interest in biomechanical studies in plants, our intention is to obtain large, homogeneous populations of single plant cells (SPCs) with “native” CWs (e.g. to test their adhesion properties) that can be manipulated as individual cells. Hence, we aim for the majority of the protoplasts to recover their CWs. Embedding is not an option as it either traps the cells, preventing manipulation, or may “contaminate” the CW surface with polymers from the gelling agent.

Several methods and protocols have nevertheless been developed specifically and claimed to yield high CW recovery (Table S2). However, preliminary results in our hand did not yield CW recovery at levels comparable to the literature values. Indeed, the existence of protocols with substantial variations in medium composition and growth conditions suggests that CW recovery efficiency may be highly lab-dependent. A protocol that works optimally in a given lab and its local conditions may not work efficiently in another lab. Parameters like temperature, local water parameters, air moisture level, local provider of chemicals and medium components, etc, could have direct or indirect effects on the medium and growth environment of the cells and may be challenging to track down. Instead, ideal medium and growth conditions may need to be tested and optimized in each lab. Furthermore, medium optimization is generally required when it comes to different species, varieties or even different tissues within the same species (Reed and Bargmann, 2021). Ideal conditions can vary for CW recovery of protoplasts extracted from leaf mesophyll to protoplasts extracted from seedlings or cell cultures. Surprisingly, despite decades of research, from a non-expert point of view, the approaches to optimize *in vitro* culture conditions almost appear as an art, often based on qualitative observations and relying on the skills and expertise of exceptional researchers and research groups with decades of experience. Most studies that document their optimization procedure report qualitative observations or largely manual measurements due to the lack of widely available quantitative and high-throughput methods. Fluorescence-Activated Cell Sorting (FACS) offers such a high throughput, quantitative and unbiased approach (Antoniadi *et al*., 2022) but unfortunately remains rarely accessible in many research labs and usually requires advanced skills and expertise.

Here we present an accessible workflow, developed to quantify cell viability and CW recovery after protoplasting. Using widely available fluorescence imaging and a custom image processing workflow, the quantitative cell wall regeneration (Q-Warg) pipeline reports several parameters of a cell or protoplast suspension, such as morphometry (size, circularity…), viability and CW staining intensities (see additional files or https://github.com/VergerLab/Q-Warg for user guide, protocol and scripts). This allows quick screening of numerous parameters to optimize and compare them with quantitative data. Thanks to this pipeline, we could optimize conditions to obtain SPCs derived from protoplasts. We also further investigated approaches to select an homogeneous population of SPCs. To test the robustness of our workflow, we also applied it in different settings, to optimize high cell viability and fast CW recovery in media designed to accelerate subsequent cell division and tissue regeneration. Finally, we demonstrate the versatility of the workflow by using it to quantify the proportion of protoplasts containing chloroplasts after protoplast extraction from seedlings. Overall, we provide a new approach and set of tools to improve efficiency, accuracy and reproducibility when working with protoplasts, their viability and their CW recovery, and demonstrate their usefulness in different scenarios.

## Results and Discussion

### Q-WARG: quantitative cell wall regeneration pipeline

The Q-Warg pipeline (<u>Q</u>uantitative cell <u>WA</u>ll <u>R</u>e<u>G</u>eneration) was originally designed as a quantitative screening approach to optimize *in vitro* culture conditions for cell viability and CW recovery after protoplasting. Note that throughout the manuscript we use the term recovery instead of regeneration to avoid confusion with the process of protoplast regeneration that implies the growth of a whole plant from one protoplast. The screening is based on brightfield and widefield fluorescence imaging, followed by quantitative image processing and data analysis (Figure 1). Such quantitative analysis helps screening optimal culture condition in a more objective and trackable way. Semi-automated imaging along with batch image processing and quantitative analysis make the process high throughput compared to classical manual quantification or qualitative observation. Compared to flow sorting methods, our approach uses tools widely available in research labs or core microscopy facilities (fluorescence light microscope), which makes it accessible to most research labs without specialized equipment. In this section we describe the main principles of this workflow, the key considerations we applied during its development as well as considerations for its usage. All the computational components of the workflow are available as supplemental material and on a GitHub repository (for the latest updates and potential bug fixes) along with a detailed protocol, user guide, and tutorial videos (https://github.com/VergerLab/Q-Warg, Method S1 to S4).

**Figure 1:**
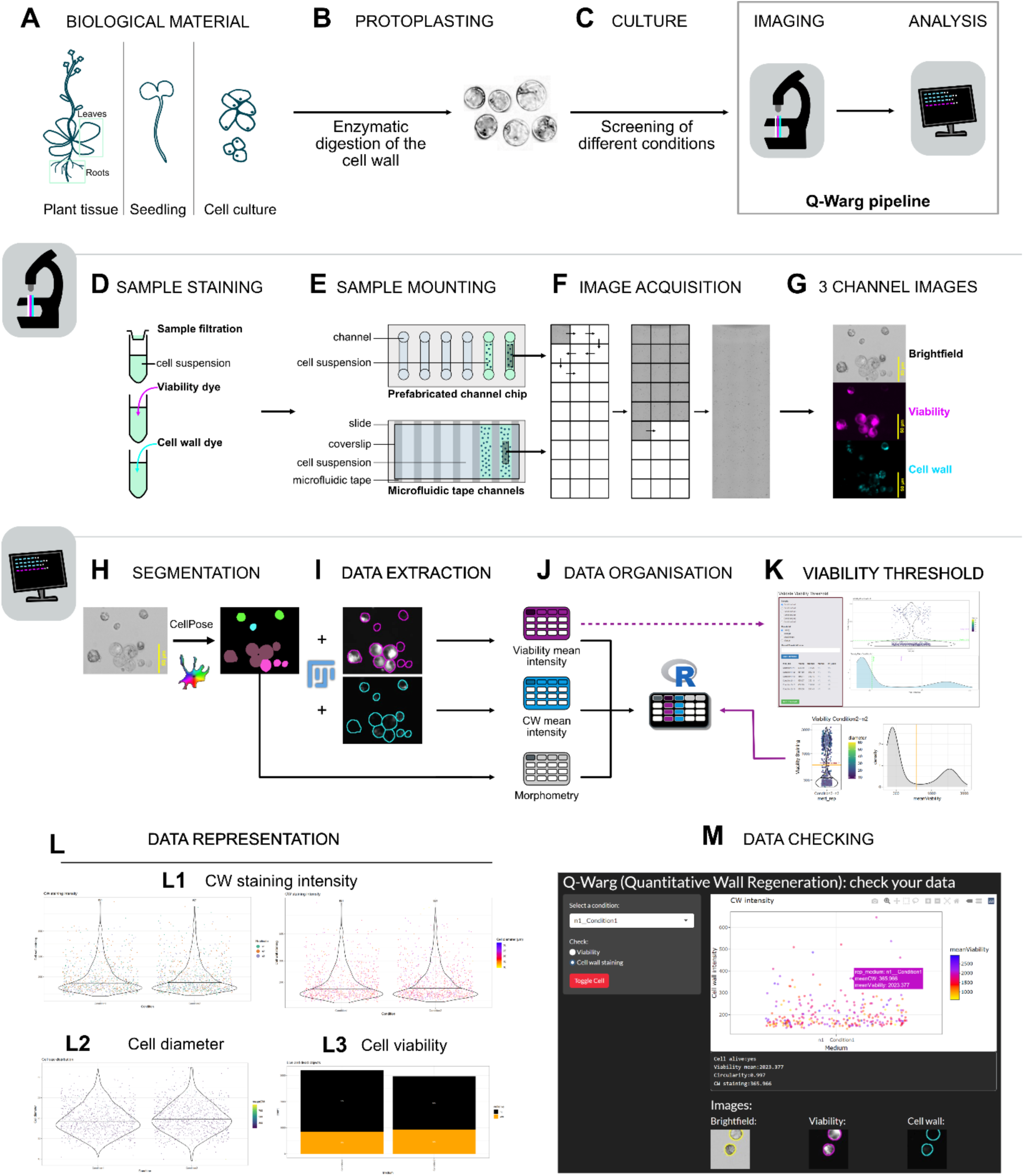
Overview of the Q-Warg screening pipeline. Q-Warg pipeline to screen and optimize protoplast culture conditions for viability and CW recovery after protoplasting. Panels describe each step from the experimental stage to the analysis. **(A,B,C)** Description of the experimental procedure in the wet lab. **(A)** Starting biological material can be any plant tissue. **(B)** Cell wall from tissue is digested by enzymes to release protoplasts. Protoplasting protocols depend on the plant tissue. **(C)** Protoplasts are cultivated in different conditions to be compared. Following a period of culture, the cell suspension is imaged and analyzed. **(D to G)** Description of the imaging process proposed in the Q-WARG workflow. **(D)** Sample staining for viability (in this paper FDA was used) and cell wall staining (Calcofluor). **(E)** Sample is mounted in prefabricated channel chips (loaded in the entry of the channel with a micropipette) or in microfluidic tape channels: microfluidic channels maintain a distance between the slide and the coverslip to avoid crushing the cells. Cell suspension is loaded by capillarity. **(F)** Image acquisition using the microscope tiling system. Large imaging by stitching multiple low magnification images to cover a large part of the sample. **(G)** 3 channels images are recorded: brightfield, viability and cell wall. **(H to M)** Description of the analysis process proposed in the Q-WARG workflow. **(H)** Segmentation based on the brightfield image using CellPose (cyto3 model). **(I)** ImageJ macro using MorpholibJ plugin to extract data from fluorescent images based on the label image and morphometry information. **(J)** Data tables are combined and organized thanks to an Rscript to be ready for analysis. **(K)** Viability threshold based on the viability staining is applied on the data to analyze only the living objects in the cell suspension. **(L)** Data plots: **(L1)** cell wall staining intensity for all conditions, the color gradient shows the cell size. (**L2)** cell diameter for all conditions, the color gradient shows the cell wall intensity. (**L3)** Cell viability proportion and living cells count. Orange shows the living cells, black shows the dead objects. **(M)** Data checking in Shiny app to make the link between quantitative data in the plot and images.

#### Sample preparation (Figure 1A-D)

After extraction and/or cultivation (Figure 1A-C), the protoplasts or cells can be stained with fluorescein-diacetate (FDA) to assess their viability and with Calcofluor to assess their CW recovery (Figure 1D). They can also be imaged with brightfield microscopy to estimate their size and sphericity. Thus, in this workflow we use co-staining with Calcofluor and FDA and image the cells with brightfield and widefield fluorescence microscopy to acquire 3-channels images: brightfield, FDA and Calcofluor (Figure 1G). Note that alternative cell viability or CW fluorescent reporters could also be used.

#### Quantification (Figure 1H and I)

Once images are acquired, they can be used to quantify cell morphology, viability and CW recovery. The first step of the analysis is to segment the cells from the brightfield channel image. Here we used CellPose (cyto3 pretrained model;(Stringer and Pachitariu, 2024)) to obtain the label images, containing individually labelled masks for each individually segmented cell or cell-like objects (Figure 1H). Note that any other segmentation method may be used at this step as long as the output is (or is made) compatible with the rest of the workflow (Method S2). We then developed a Fiji macro to extract data from the label image and the corresponding fluorescence channels (Figure 1I). In this macro we use the plugin MorpholibJ (Legland, Arganda-Carreras and Andrey, 2016) to quantify several morphometric parameters (including cell size and circularity) from the segmented labels, as well as to measure the mean signal intensity for the viability and CW fluorescence channels. The macro generates three data tables containing information about the morphometry, viability signal intensity and CW signal intensity.

#### Analysis (Figure 1J-K)

We conceived an R based pipeline (R notebook) for data processing and visualization (Figure 1J-L). From the three data tables generated by the Fiji macro, we first use the R script to combine the information in a single table (Figure 1J). Since many segmented cell-like objects may not be cells or be alive, we use the FDA staining value to screen the segmented dataset and keep only what can be considered as living cells based on FDA staining intensity. Due to the quantitative nature of the analysis the value of the signal quantified for viability and CW is not binary. There may also be heterogeneities of background signal and noise acquired from one image to the next. Thus, even dead cells may appear to have a background level of fluorescence as quantified by the workflow. Nevertheless, we generally observe a clear bi-modal distribution of fluorescence signal for FDA staining, with one low fluorescence intensity peak corresponding to dead cells and a higher intensity peak for live cells (Figure 1K). For unbiased analysis, the workflow allows the use of automated thresholding techniques, to differentiate between dead and living cell populations. On our samples, Huang and Triangle methods were the most accurate, but other thresholding algorithms can also be used such as MaxEntropy. It is noteworthy that the algorithms consistently identify a threshold, even in scenarios where the entire cell population falls into a single category (typically all dead). Consequently, there is a potential for misclassification, where a subset of cells might be wrongly categorized as viable in a predominantly non-viable population. To address this limitation, we developed an R-based Shiny application (Figure 1K) that facilitates user intervention in the thresholding process. This tool allows for the selection of the most appropriate automated method or the application of a manual threshold. However, it is crucial to emphasize that any manual threshold adjustment must be applied judiciously to maintain data integrity and avoid introducing bias into the analysis. To further avoid having false positive, like debris that are close to a living cell and are illuminated by the FDA staining of this cell, the workflow also applies a cell circularity filter. Indeed, the isolated living cells are generally very spherical as they recover their CW without environmental pressure (in suspension) while debris have various shapes. Based on preliminary observations, we chose a high circularity index (0,9) to discriminate single cells from debris and cell aggregates.

#### Results representation (Figure 1L)

The processed and filtered data is then plotted (Figure 1L). The first representation shows the CW intensity for each CW recovery condition tested, where the colour gradient of the dots represent the size of the cells (Figure L1). The number of cells alive in each condition is noted at the top of the graph. The second plot shows the size of the cells, where the colour gradient of the dots represents CW staining intensity (Figure L2). Both plots are also saved with the colour of the dots representing each experimental replicate. The last one shows the proportion of living cells compared to dead cell-like objects (Figure L3). While viability staining data usually shows a clear bimodal distribution from which we can define a threshold for dead versus alive, CW intensity rarely shows such clear distribution. Thus, contrary to what is usually described in publications reporting qualitative CW recovery from protoplasts, this pipeline doesn’t output a binary yes/no answer for CW recovery. We believe this is a more accurate output which better reflects the progressive nature of CW recovery as opposed to the more binary nature of cell viability.

#### Inspection (Figure 1M)

Because this workflow segments and quantifies in batch thousands of cells at a time, there is always a risk of inaccurate segmentation or signal quantification leading to errors and biases in the final analysis. For instance, an incorrectly segmented cell could encompass a brightly stained CW debris, thus incorrectly assigning a high CW recovery value to that cell. With regular plots, it is nearly impossible to trace back a specific point in a plot to the corresponding cell in the raw image from which data was extracted and check the validity of the quantification. We thus aimed to develop a tool that would allow us to interactively check the dataset from the plotted data. We developed a Shiny app (R) making use of the plotly library (https://plotly.com/r/ ; Figure 1M) where we can choose which condition to observe and look at the cells individually by browsing through the plot. This allows us to check if the highest points are indeed real living cells or debris highly stained with calcofluor. It is also possible to randomly check points to ensure the value reported fit what a user can see in the microscopy image as well as if the segmentation has been done properly.

#### Advantages and versatility

With the Q-Warg pipeline, many conditions to maintain a protoplast culture and recover CWs can be rapidly screened at once. It can be used after any protoplasting method and is not limited to the experimental conditions described here. Multiple parameters can be tested at once and compared quantitatively based on automatized measurements, eliminating the bias of phenotypical/qualitative observations. The workflow requires fluorescence images of the cell suspension as input. Those images can be acquired with any fluorescent microscope making this pipeline usable by most of the labs. The possible low sensitivity of widefield fluorescence microscopes can be one downfall compared to techniques that allow high detection of low signals such as FACS. However, using a confocal microscope can improve the detection of such low signals. All software (CellPose, Fiji and RStudio) are free, do not require high computational power nor specific operating system.

In this paper, special attention is paid to CW recovery after protoplasting, yet the viability alone is also a crucial parameter monitored after any type of protoplasting based experiments, which can be quantified in an accurate and traceable way thanks to this pipeline. As this pipeline is based on fluorescence images, other parameters could be quantified instead of the CW and viability (e.g. transformation efficiency, gene expression, etc.). While the pipeline currently reports simple metrics (shape, size and fluorescence intensity), the data acquired could be further refined using machine learning to classify and cluster cells or with deep-learning models to improve the accuracy of the analysis (i.e. by excluding signals coming from CW debris attached to a protoplast or predicting cell viability without the use of a binary threshold).

### Screening for improved CW recovery after protoplasting liquid-grown habituated Arabidopsis cell culture

We developed the Q-Warg workflow to solve the challenge we encountered when attempting to obtain “single plant cells” (SPC; Isolated cells with recovered CW). Our aim was to obtain large populations of SPCs with relatively homogeneous size, recovered CWs, and no cell divisions, to establish them as a model system for plant CW and biophysical studies. Their largely spherical shape is ideal for physical and mechanical consideration and potential computational modelling approach. Such cells could then be used in studies of cell adhesion strength or CW mechanics. Our intention was to start with protoplasting and letting the cells recover their CW before using them. However, while many methods, publications and protocols exist to promote CW recovery after protoplasting (Schirawski, Planchais and Haenni, 2000; Wu *et al*., 2009; Kuki *et al*., 2017; Pasternak, Paponov and Kondratenko, 2021; Jayachandran *et al*., 2023), none of the ones we tried yielded the expected results in our hands. Our preliminary observations led us to choose Arabidopsis habituated root cell cultures (Pesquet *et al*., 2010; Ménard *et al*., 2017, 2024) as starting material since cells extracted from it appeared more homogeneous. Similarly, our preliminary tests led us to use a modified protoplast extraction protocol (Yoo, Cho and Sheen, 2007) and a modified regeneration medium (M ; mannitol instead of trehalose (Kuki *et al*., 2017)) as the foundation for further optimization. Those preliminary protocol refinements, based on literature searches (Table S2), were assessed with qualitative observations of the protoplasts’ shape and debris in the medium (issued from cell death). To go further and assess our progress methodically we needed a quantitative way to compare the different conditions tested. Accordingly, we switched to developing and using our Q-Warg workflow.

From all parameters that were previously reported to have an impact on the protoplasting and the CW recovery (Tables S1-S2), we decided to focus on two main aspects: sugars (Sx media; Figure 2) and hormones (Hx media; Figure S1). The sugars have two different roles: keeping the osmolarity of the medium (mannitol, trehalose (Kuki *et al*., 2020)) and feeding the cells (glucose, sucrose). In the starting protocol NAA was used as an auxin source. We also tested the effect of 2,4-D and IAA on the CW recovery (Figure S1).

**Figure 2:**
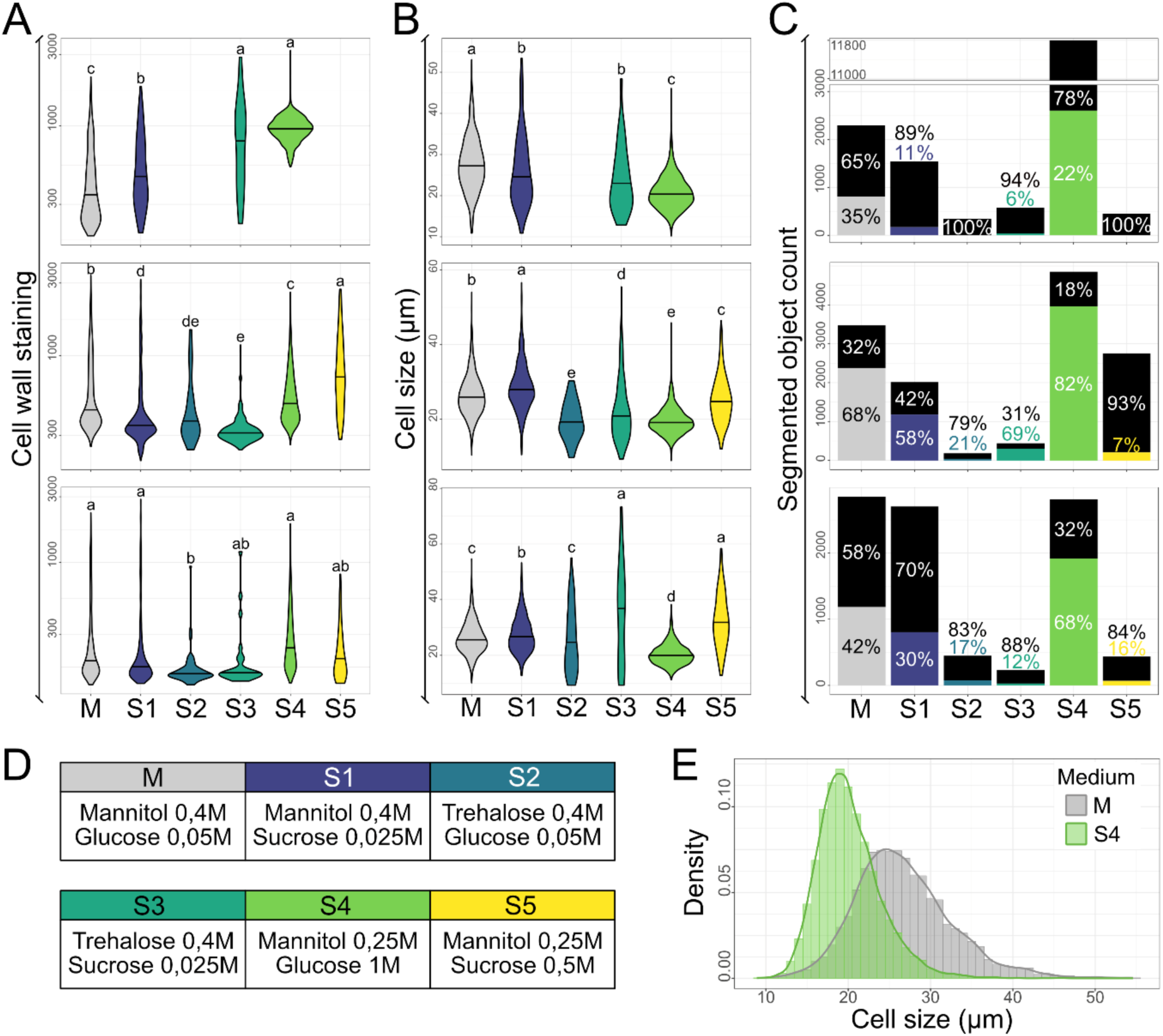
Screening for the effect sugar composition in the medium to improve the CW recovery. (**A-B-C**) Plots from data processed with the Q-Warg pipeline on the effect of sugar composition in the medium for cell wall recovery, cell size and viability, 4 days after protoplasting. Each plot corresponds to one replicate (n=3). (**A**) Cell wall mean intensity after staining with Calcofluor. (**B**) Cell size (diameter in µm). (**C**) Viability plots displaying the number of segmented objects (including debris, dead cells or living cells). On violin plots, letters describe the statistically significant differences between populations determined by one-way ANOVA followed by Tuckey’s HSD test (p < 0,05). In bar plots, each bar shows the total number of segmented objects, and the proportion of objects negative for viability staining (in black) and living cells (in color). Chi-square test of independence (significant if p < 0.05) was performed, followed by pairwise comparisons using the pairwise.prop.test function with Bonferroni correction for multiple testing. Results are in Data S1 (pages Fig2_viability-stats). (**D**) Table with the tested media composition. All media contain Gamborg B5, MES, NAA 1µM and the corresponding sugars. (**E**) Cell size distribution for M and S4 media from the 3 replicates shown in B.

We found that the impact of the type of sugars used on the viability of the cells is quite high (Figure 2D). For the 3 replicates, media S2, S3 and S5 showed a low number or no living cells. Both S2 and S3 contained trehalose instead of mannitol, which did not seem to help our cells to recover their CW. S1 and S5 media both contained sucrose and displayed either a low number of living cells and/or low CW staining intensity mean compared to the reference medium M (Figure 2B). Interestingly, S4 medium provided comparable results in CW staining as medium M. Moreover, the number of living cells after 4 days in culture was higher. Additionally, the cells were, on average, smaller and more homogeneous in size than those in medium M (Figure 2C,E). The smaller size may also explain why the cells in medium S4 recover their CW more efficiently, as they need less CW components to be secreted by the cells to recover the CW. In addition to the quantitative data showing that medium S4 was a good candidate, the cell suspension in medium S4 showed well isolated cells while in medium M more clumps/aggregates were present.

The results of the auxin screening were less clear. In our hands, the source of auxin did not seem to have any major impact on the CW recovery at the concentrations we tested when using our base medium (Figure S1B). Nevertheless, medium H1 seemed to promote a little bit more the CW recovery than medium M suggesting that using a higher concentration of NAA could help the protoplasts to recover their CW. Zhang et al., (Zhang *et al*., 2024) showed that higher of auxin enhanced the CW recovery. We then screened several concentrations of NAA (0 to 30µM, every 5µM), this time using medium S4 as base, to check if we could further increase CW recovery in our medium (Figure S1E). For both 5µM and 10µM a significant increase of CW recovery was observed. As the proportion of living cells was higher with 5µM of NAA, we choose to work with this concentration (Figure S1E).

Ultimately with this approach using the Q-Warg pipeline, we successfully improved our medium for CW recovery on the protoplasts extracted from Arabidopsis habituated root cell culture maintained at Umeå Plant Science Centre (UPSC). Starting with the M medium, we found that the S4 medium led to a 1.61- to 3.22-fold increase in the total number of living cells, up to a 2.12-fold increase in CW staining mean intensity, and more homogeneous cell sizes (from 26,7±5,89 µm for M to 20,1±3,82µm for S4). Building on the S4 medium, increasing the NAA concentration from 1 to 5µM resulted in a 1.12 fold increase in the total number of living cells and 1.08 fold increase in cell wall staining mean intensity, while cell size and homogeneity remained similar (from 19,9±4,71 µm to 19,1±3,92 µm). With this, we propose an improved medium for our use case, based on the S4 medium and containing 5µM NAA that we name CRRUM (CW Recovery Root cells UPSC Medium).

Note that we performed the original sugar and hormone screen with three independent biological replicates to assess the robustness of the pipeline as a screening approach. The 3 biological replicates, while showing some differences are largely in accordance with each other. We thus believe that our workflow can also be used as a robust and high throughput screening tool. First by testing various media and conditions and then using replicates for a subset of more promising media to confirm and refine choices in medium optimisation.

### Further selection of single plant cells with recovered CWs

Despite our efforts to improve CW recovery under our lab conditions, we have yet to identify a medium that reproducibly enables close to 100% of the cells to fully recover their CW. Furthermore, with the Q-Warg pipeline, we quantify fluorescent signal from CW staining which highlights the wide disparity in CW recovery. There is no clear bimodal distribution of cells with and without recovered CWs. Not all cells seem to recover their wall at the same pace. This raises the question at which point do we consider the CW as “recovered”? Furthermore, while our workflow effectively characterizes these properties, it does not allow selection of cells with recovered walls for further use.

One possibility to overcome this limitation, is to sort cells with FACS based on cell size and fluorescence staining intensity. Before sorting, the cell suspension was stained for viability and CW recovery assessment. The sorted cell suspension contained only isolated cells that were positive for both viability and CW staining (Figure S2). However, during sorting, the cells are exposed to high pressure and friction forces, which may be harmful and could reduce yield. This method is suitable if a large population of cells meets the sorting requirements, and the loss of some cells does not significantly affect the outcomes. Using FACS also implies the availability of suitable equipment, ideally sorting to be done in sterile conditions, and it requires specific skills that may not be widely accessible in many plant biology labs.

One key function of the CW is to balance turgor pressure coming from the cytoplasm. After CW removal, protoplasts are initially kept in an iso-osmotic medium to avoid bursting before the CW is recovered. In turn, it can be argued that a non-biased binary (yes/no) estimation of CW recovery would be whether or not a cell can sustain its own turgor pressure when it is placed back in pure water or in the normal cell culture growth medium (Figure 3C). A sudden change of medium osmolarity could provide both a binary estimate of CW recovery when tested through our quantitative workflow and serve as a mean to eliminate (burst) cells with non-recovered CW from a population. As a proof of concept, we first exposed the cell suspension (4 days after protoplasting in S4 medium) to pure water. To monitor the effect on individual cells, we used a microfluidic chip containing U-shaped traps (Sakai *et al*., 2019). This allowed us to keep the cells trapped in a fixed field of view for live imaging while allowing a dynamic change of the liquid medium. Switching from the S4 medium to pure water led most of the observed cells to inflate and burst within a few minutes (Figure 3A-B). Observations using both viability and CW staining (Figure 3B) further suggest that cells with a qualitatively well-recovered wall (clear fluorescent signal) do not burst upon change of medium while protoplasts that do not appear to have CW staining, burst. It is also likely that smaller walled cells showed a stronger resistance to the osmotic shock as a similar increase in turgor pressure would lead to higher stress in the CW of larger cells ((Sapala *et al*., 2018) Figure 3B). These preliminary observations indicate that our hypo-osmotic shock approach could be useful at a larger scale. We then turned to look at the effect on large scale cell populations and characterized the effect with the Q-Warg workflow. Pure water appeared to be too much of an osmotic shock which may even lead to bursting of walled cells (Figure 3AB). We used two media that contained less osmotic support than extraction or cultivation media for protoplasts to perform the osmotic shock: CRRUM without mannitol (CRRUM-noM) and the initial cell culture medium MS3%. The cells were transferred after 4 days in culture in CRRUM into their new medium. Surprisingly, quickly, all the cells in MS3% died suggesting this medium transition had a negative effect on cell survival regardless of their CW recovery status (Figure S3C). In CRRUM-noM, we observed an effect of the decreased osmotic pressure on the size of the cells as they inflate (average cell size increases) quickly after changing medium (Figure S3A). There was also a strong reduction in the number of cell-like objects counted, and the percentage of objects considered as living cells. Surprisingly, while we could expect the remaining living cells to be primarily strongly stained for CW, our quantification revealed the opposite trend, with cells having on average lower CW staining than the cell population kept in medium with osmotic support (Figure S3B). Nevertheless, after the osmotic shock, the CW fluorescence intensity average signal increased over cultivation time (3 days) while this trend is not seen in the medium with maintained osmotic support (Figure S3C). Thus, while we did not obtain the expected effect of binary selection of cells with recovered walls, the osmotic shock approach could still be useful to promote increased CW recovery overtime as suggested in other methods (Sakai *et al*., 2019; Pasternak, Paponov and Kondratenko, 2021).

**Figure 3:**
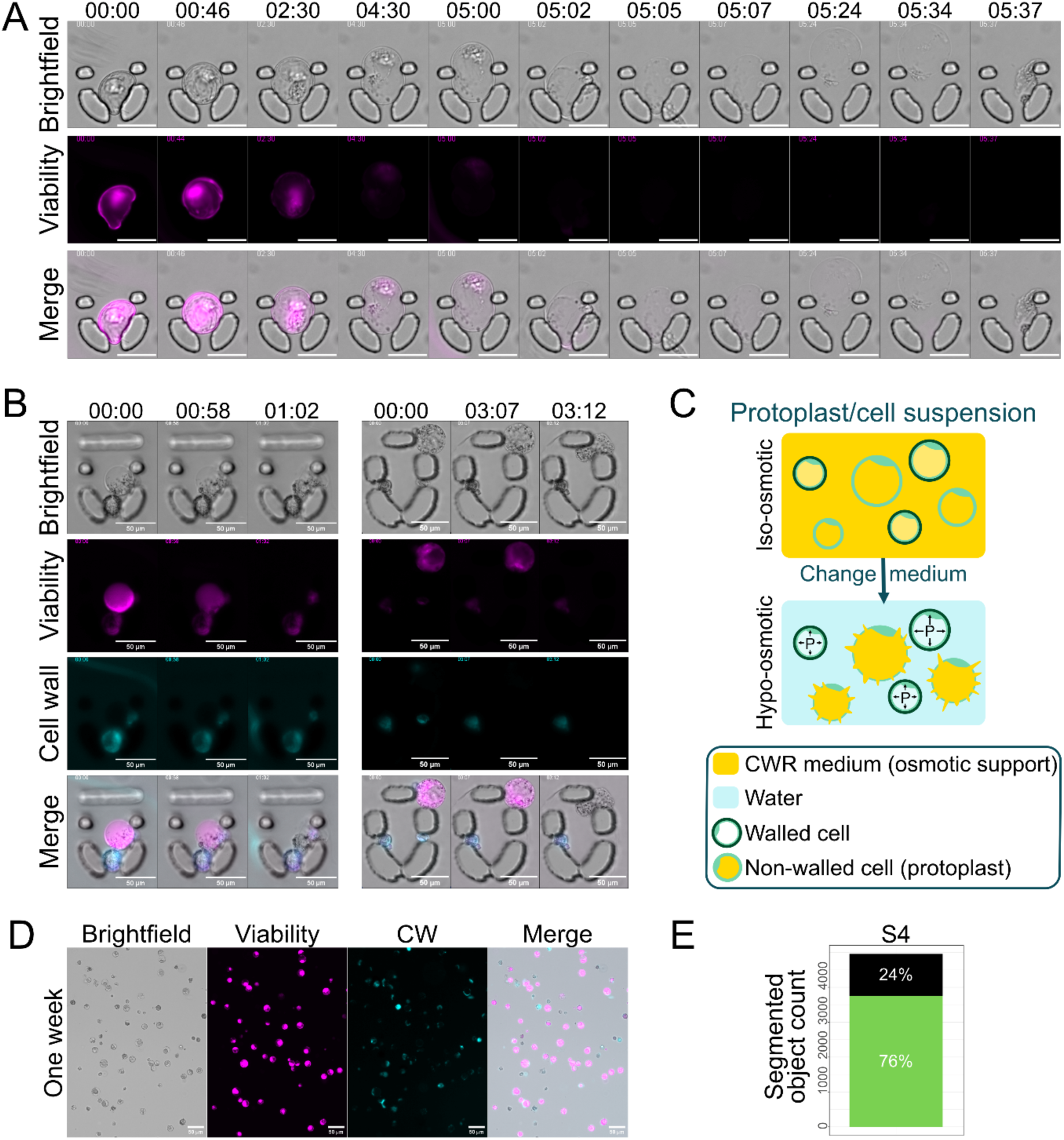
Cell selection based on osmolarity differences in culture medium and single plant cells. (**A-B**) Osmotic shock performed on SPC 4 days after protoplasting. MilliQ water is flowing inside the chip from time 0. The change in osmolarity makes the cells inflate and burst. The viability signal shows that the cell was alive at the start of the experiment. Note that the decrease in signal does not entirely correspond to the death of the cell as FDA stays fluorescent after conversion by cellular enzymes. In this case the decrease of the signal is partially due to the bleaching of fluorescence. (**A**) Opened trap. (**B**) Closed traps. Cell walls were stained with Calcofluor. (**C**) Scheme representing the osmotic shock strategy: the protoplast culture contains both walled and non-walled cells. the non-walled cells (protoplasts) are more susceptible to changes in medium osmolarity. In a hypo-osmotic medium, only walled cells will survive. (**D**) Single plant cells cultured into S4 medium for 1 week. Cells are stained with FDA for viability and calcofluor for CW. Imaged in microfluidic taped channels. (**E**) Quantification of living cells after one week of culture in S4.

### Assess isolated cell viability

To assess the behaviour and viability of SPCs over time, we used a small 3D printed microscope – Openflexure (Collins *et al*., 2020)– for long term observation. Cells were plated on a glass-bottom Petri dish with a poly-lysine coating to maintain them in place. Using this DIY setup, the cells were maintained and imaged every 15min for about 5 days. We could observe movement inside the cytoplasm of the cells showing that they were still alive (Movie S1). In parallel, cells were kept for a week in the dark and stained with viability and CW staining to check their state after a week of culture in S4. Cells were still showing a positive FDA signal indicating that they could survive for a long time as single cells (Figure 3DE). Interestingly, cells did not seem to divide in both setups, which gives us a large time window to use SPC in further experiments. To pursue divisions and calli development, cells might require other nutrients and/or hormones. This step could also be optimized using the Q-Warg pipeline.

### Screening for CW recovery in media aimed to enhance cell division

First, to test the reusability of the Q-Warg pipeline, a trial run has been conducted in other hands in a different lab with different biological material: at the Laboratory of Biochemistry, Wageningen University, using PSB-D cell culture derived from stem explants of *Arabidopsis thaliana* Landsberg erecta (Ler) ecotype. Unlike the previous results, imaging was done with a confocal microscope, showing the adaptability of the pipeline. We first conducted a similar screening of NAA concentration (ranging from 0 to 4µM) within the initial M medium during the first days of culture after protoplasting to assess cell viability. We could observe that in the absence of NAA, less than 30% of the segmented objects were living cells after 2 days of culture. By increasing the NAA concentration, we could note a positive effect on cell viability. After 2 days of culture, the more NAA was present in the medium, the larger the viable proportion of cells was (Figure 4A and S4). We also checked whether the CW recovery was increased at higher NAA concentration, yet in those conditions, we could not observe a significant difference in CW recovery (Figure S4A).

**Figure 4:**
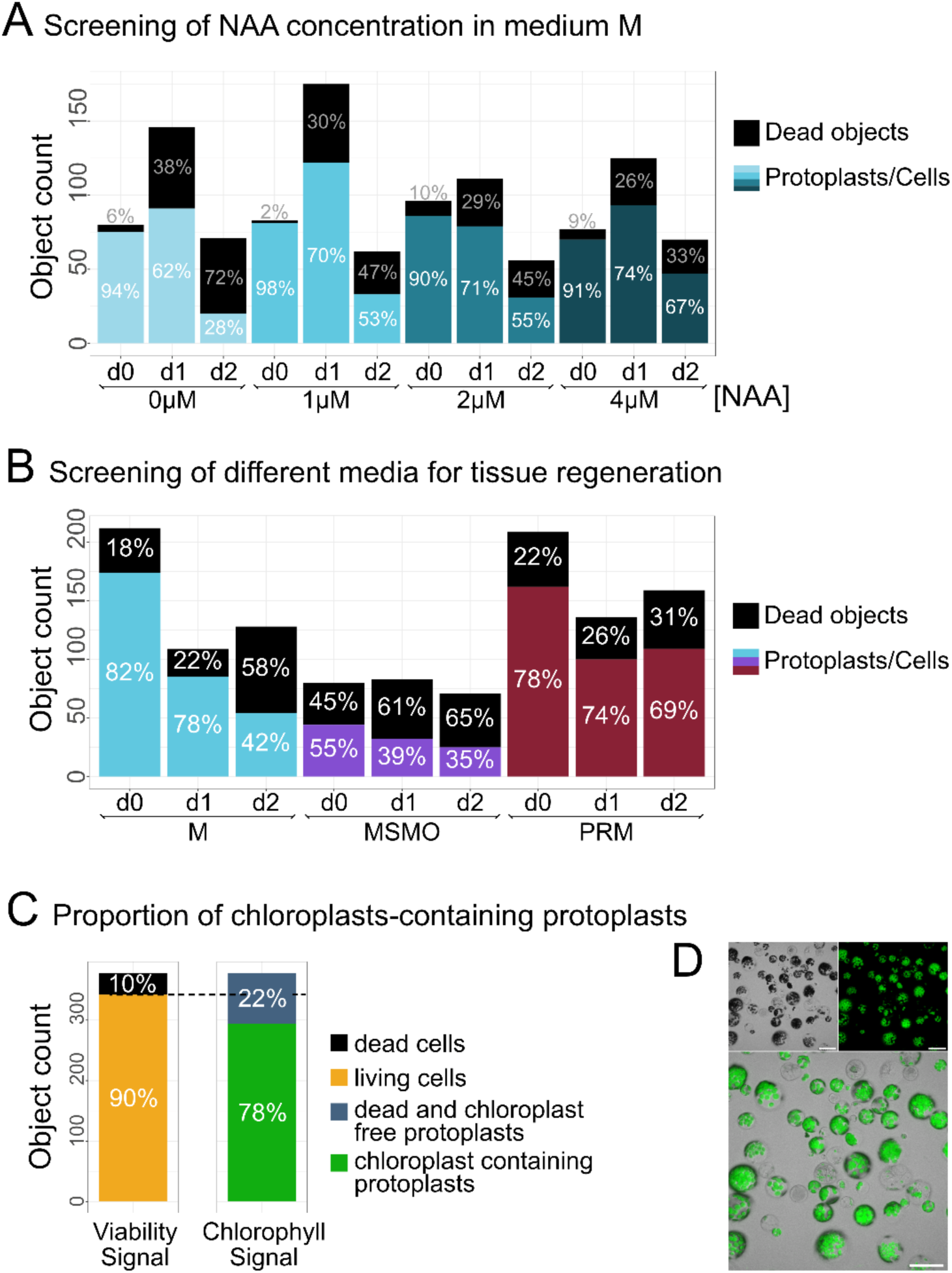
Challenging the Q-Warg pipeline. (**A**) Viability plot for NAA concentration screening in M medium. 4 different concentrations of NAA have been tested. d0 corresponds to the protoplasting day. (**B**) Viability plot for medium screening. Three media for tissue regeneration were tested. d0 corresponds to the protoplasting day. (**C-D**) Protoplast extracted from *Arabidopsis thaliana* seedlings and imaged the day of extraction. (**C**) Left bar plot represents the proportion of living cells in yellow (90%). The right bar shows the proportion of chloroplasts-containing cells based on chlorophyll autofluorescence (78%). The dotted line reports the proportion of dead cells onto the chlorophyll signal histogram. (**D**) Protoplast suspension in brightfield (top left), autofluorescence of chlorophyll (top right) and merged image (bottom). Scale bar 50µm. In plots, object count corresponds to the number of segmented objects (including debris, dead cells or living cells). Each bar shows the total number of segmented objects, and the proportion of objects negative for viability staining (in black) and living cells (in color). Protoplasts are issued from PSB-D cell culture. Chi-square test of independence (significant if p < 0.05) was performed, followed by pairwise comparisons using the pairwise.prop.test function with Bonferroni correction for multiple testing. Results are in Data S1 (pages Fig4_viability-stats).

In light of these results, we initiated a medium screening to determine the requirements for promoting micro-calli formation after protoplasting. Since cell divisions can only occur once the CW has been recovered, the first goal was to establish which parameters maintain a high viability and secondly promote CW recovery but using base media that contain hormones and nutrients aimed at enhancing cell division and regeneration. The screening was performed with three media over three days starting on the day of the protoplast extraction. In two tested media, “Murashige and Skoog with Minimal Organics” (MSMO) and “Protoplast Regeneration Medium” (PRM ; (Chupeau *et al*., 2013)), kinetin, a form of cytokinin was used to promote cell division when associated with auxin (respectively NAA or 2,4-D ; (Barciszewski, Massino and Clark, 2007)). To compare with the previous results obtained with the root cell culture used in the beginning of this study, we also tested the M medium. Over this period of observation, we did not notice a significantly better medium for CW recovery. Cells in M and MSMO showed a similar increase in CW signal, but viability decreased to about 40% of the segmented objects (Figure 4B). PRM medium did not show an improved CW recovery compared with M and MSMO (Figure S4B). However, the proportion of living cells after 2 days was maintained at 70% of the segmented objects, which made this medium promising. Future work will determine the best condition to achieve quick cell divisions to reach the micro-calli stage.

### Pipeline versatility: Quantification of chloroplast-containing cells

The Q-Warg pipeline is based on brightfield and fluorescence imaging and quantification and is not limited to viability and CW staining. The intensity quantification can in principle be done on any fluorescence marker or autofluorescence signal. To show the versatility of the pipeline, we quantified the proportion of chloroplast-containing protoplasts extracted from whole seedlings. Using the viability proportion plot, we first quantified the proportion of living cells in the suspension with the viability staining (FDA). Then inside this population of living cells, we used the workflow to quantify the number of cells containing chloroplasts (Figure 4D-E). To do this, instead of taking the viability staining measurement, we used the autofluorescence of chloroplasts. The viability plot given by the pipeline is then used to report the cells without chloroplast signal as “dead” and the living cells are positive for the fluorescence and therefore are the ones that should contain the chloroplasts. In this experiment, 78% of the protoplasts extracted from seedlings, contained chloroplasts.

## Conclusion and perspectives

In this work, we introduced the Q-Warg pipeline, an imaging and image processing-based workflow aimed at making studies of protoplast culture, viability, and CW recovery more accessible, quantitative and standardized. We further applied it in several use cases to show its usefulness in different configurations, demonstrating its robustness and versatility.

Our initial aim when developing this pipeline was to create a tool that we could confidently use to assess our progresses when optimizing CW recovery media for the obtention of “single plants cells”. Thus, in this work, we report the steps taken in this direction by testing the quantitative effect of changing a subset of the CW recovery medium composition, namely the sugars and auxins used, and finding an improved medium. However, given the number of parameters that may influence protoplast culture and CW recovery (Table S1), and the range of possible values to be tested for each parameter, there is a virtually infinite number of media combinations to be tested. This is obviously impossible for practical reasons, but it is in principle possible to use Design of Experiment approaches to explore parameter spaces and define optimal settings without having to test every possible combination (Cano, Moguerza and Redchuk, 2012; Peng *et al*., 2022). We also believe our workflow could further help in the future to streamline the media optimisation process. The standardized nature of the pipeline also aimed to increase the reusability of the data. We provide a detailed user guide for the whole procedure (Method S2) and aimed to make the computational workflow user-friendly. We also strongly encourage future users to deposit their data in publicly accessible repository for future reuse. This is because, during the medium optimisation process, in practice it is more reasonable to test a restricted set of parameters, until reaching an acceptable result and moving on. Researchers have been doing this for several decades, but unfortunately those optimisations steps are rarely reported, nor quantitatively measured, and there is no standardized format available. Instead, *in vitro* culture medium optimisation is almost referred to as an art, only mastered by a few experts with decades of experience. We believe the instinctive expertise acquired by such expert could be understood scientifically and be made available more broadly if we can generate enough standardized data on protoplast viability and regeneration, and study them through advanced multivariate analyses or novel deep learning approaches. We could for instance imagine training a model with a large dataset gathering matching information on the parameters used in the media, types of tissues, varieties, species, growth conditions etc., and matching quantification of viability and CW recovery. In turn we may be able to interrogate the model to predict a set of media to test for further optimisation, given a species, variety, tissues, and growth conditions of choice. This would effectively allow a much shorter time for medium refinement, by using complex knowledge from prior optimisation experiments as a starting point for a subset of media to test. In parallel, the Q-Warg pipeline could be used for quantitative screening of large sets of mutants with improved or impaired protoplast viability or CW recovery, in order to decipher the biological mechanisms regulating these processes.

The pipeline we developed here is also potentially highly versatile. Because of our original aim, in this paper we focused mainly on the cell viability and the CW recovery after protoplasting. However, the pipeline can also be used to explore other parameters. Here we also used it to determine the percentage of chloroplasts-containing cells inside a population of protoplasts extracted from seedlings. This workflow could also be used to check the transformation efficiency with a fluorescent reporter or other parameters that can be observed quantitatively with fluorescent dyes or reporters at the cell level.

Encouraging open data practices and leveraging deep learning approaches thanks to datasets acquired with such tools could help to make significant strides in unravelling the biological complexity of protoplast, ultimately contributing to more robust and reproducible research outcomes and biotechnological applications.

## Methods

### Biological material

#### Root cell culture

The cell culture used in this study (unless stated otherwise) is derived from root cells of *Arabidopsis thaliana*, ecotype Colombia, and was previously reported (Pesquet *et al*., 2010). Prior to our set of experiments the cell culture was re-screened for habituation (Ménard *et al*., 2017, 2024). Growth medium MS 3% contains Murashige and Skoog medium including vitamins (M0222, Duchefa), 3% sucrose (VWR) and MES 20mM. The pH is adjusted to 5.7 with KOH. Cells are growing in the dark under constant shaking (22°C, 120rpm). The cell culture was maintained weekly by inoculating 72mL of fresh MS3% with 8mL from the 1-week-old culture.

#### PSB-D culture

*Arabidopsis thaliana*, ecotype Landsberg, cell suspension cultures (PSB-D) were donated by Geert de Jaeger (ABRC stock no. CCL84840) (Menges and Murray, 2002; Van Leene *et al*., 2011). MSMO contains Murashige and Skoog Basal Salts with minimal organics 0.44% (Sigma-Aldrich) and sucrose 88 mM. The pH is adjusted to 5.7 with KOH. After autoclaving, NAA 2.7µM and kinetin 0.23µM were added. PSB-D cultures were grown in 50 mL MSMO (dark, 25°C, 135rpm) and were maintained weekly by taking 2.5 mL from the 1-week-old culture with 47.5 mL of fresh MSMO.

#### Seedling growth

Arabidopsis thaliana, ecotype Columbia-0 (Col-0), seeds were sterilized by washing them with ethanol 70%, followed by a treatment with NaOCl 0.8% for 3min. Seeds were washed 5 times with ultrapure water and were kept, in water, in the dark at 4°C for two days. Seeds were then sown on MS agar plates. MS agar plates contain half-strength Murashige & Skoog medium (Duchefa), MES 5mM, and plant agar 0.8% (Duchefa). The pH was adjusted to 5.7 with KOH. The plates were vertically incubated at 22 °C under a long day growth period (16h light, 8h dark).

### Protoplast extraction

#### Root cell culture

The protocol was adapted from (Yoo, Cho and Sheen, 2007) and is detailed in the protocol (Method S1). In brief, cells are incubated in enzymatic solution for 4 hours in the dark at 120rpm, 24°C in non-treated 6-well plates. Enzymatic digestion is stopped with W5, and protoplast suspension is filtered in 70µm nylon basket before centrifugation and resuspension in culture medium. For all root cell culture protoplast extractions, the cell culture was 4 days-old (4 days after passing).

#### PSB-D protoplast extraction

PSB-D protoplasts are extracted similarly to root cell culture protoplasts, with the following adjustments. A 5-day-old PSB-D culture was used. The culture was incubated in the dark at 200rpm and 25°C. Protoplasts were filtered through a 40µm nylon basket. The final pellet was resuspended in 1mL of the desired medium. Protoplast density was adjusted to 0.8*10^6^ cells/mL and 200µL of suspension was transferred to each well of an Ibidi µ-Slide (8 Well high, Polymer Coverslip, Uncoated). Slides were placed in a dark growth chamber at 22°C for 3 days.

#### Seedling protoplast extraction

The protocol is detailed in Method S1. Protocol is adapted from (Zhai, Jung and Vatamaniuk, 2009) for protoplast extraction of 18-days old seedlings with the following modifications. The protoplast suspension was filtered through a 40µm nylon basket. W5+ was used instead of W5 to retrieve protoplasts within the sucrose gradient. W5+ contained glucose 5.6mM and additional KCl 6mM dissolved in W5. Protoplast density was adjusted to 0.8*10^6^ cells/mL and 200µL of suspension was transferred to each well of an Ibidi µ-Slide (8 Well high, Polymer Coverslip, Uncoated). Seedling protoplasts were imaged directly after extraction.

### CW recovery medium

Initial medium for CW recovery, medium M, contains Gamborg B5 (Gamborg’s B-5 Basal Medium with Minimal Organics, G5893, Sigma-Aldrich), Mannitol 0.4M, Glucose 0.05M, NAA 1µM, MES 20mM filled up to the wanted volume with MilliQ water and pH 5.7 adjusted with KOH. The different combinations of sugars and hormones that have been tested are reported in (Figure 2A). The protoplasts were transferred directly after extraction in medium for CW recovery and kept in the dark at 22°C without shaking. After screening the medium CRRUM was used for CW recovery with root cell culture: S4 with NAA 5µM. For all changes of medium, the cell suspension was placed in Falcon 50mL tubes and centrifuged at 200g in a swing-rotor centrifuge 3min at room temperature. The supernatant was carefully removed by pipetting and the new medium was added. Cells were resuspended by gently turning over the Falcon tube. The media used for CW recovery with the PSB-D cells were CRRUM, MSMO (previously described) and protoplast regeneration medium (PRM). PRM was modified from (Chupeau *et al*., 2013) and contains half-strength Murashige & Skoog medium (Duchefa), MES 20mM, mannitol 0.3M, glucose 0.2M, 2,4-D 4.5µM, kinetin 0.46µM filled up to the wanted volume with MilliQ water and pH 5.7 adjusted with KOH.

### Microscopy sample preparation and imaging

#### Protoplasts derived from root cell culture

Viability was assessed by fluorescein-diacetate (FDA; F1303 Thermofischer). Four minutes before imaging, FDA was added to the cell suspension to reach 8µg/mL. For CW characterization, the cells were stained just before imaging with calcofluor white (Calcofluor White Stain, Sigma-Aldrich) diluted 1:100 directly in the cell suspension. Double stained cells were then loaded in an Ibidi channel chamber (µ-Slide VI 0.4 Bioinert) or an in-house chamber. In-house chambers are composed of a microscopy slide on which strips of microfluidic tape (S5005DC, Adhesive Applications) are put to form channels and maintain a distance from the coverslip (see protocol file Method S1). Fluorescence imaging was done with Leica DMi8 inverted brightfield and epifluorescence microscope with a motorized stage and navigator function. The microscope is equipped with a 10X objective lens (NA 0.32, air objective) and Leica DFC9000GT camera mounted on a 1X C-mount adapter. Tile images made of 3 x 9 (27) individual images (12bits, 2048×2048 resolution for each image, pixel size of 1.048×1.048µm) were acquired with the Navigator function of the microscope using a 10 % overlap. Tile images were merged with the LAS X software (Leica Microsystems, Germany). Before acquisition, autofocus is done on the brightfield channel. 3-channels were acquired: brightfield, FDA signal excited with LED 460nm, and emission detected above 515nm (long pass filter) and calcofluor signal excited with LED 365nm and emission detected between 435 and 485nm.

#### Protoplasts derived from PSB-D and seedlings

Protoplasts were imaged in Ibidi µ-Slide (8 Well high, Polymer Coverslip, Uncoated) with an inverted Nikon ECLIPSE Ti2 microscope equipped with a Nikon C2 confocal laser scanning head and a 40X (NA 0.95) dry objective. Five minutes before imaging, protoplasts were stained with FDA 5µg/mL. PSB-D protoplasts were also stained with calcofluor white (Calcofluor White Stain, Sigma-Aldrich) diluted 1:100. Calcofluor white/chloroplasts and FDA were excited sequentially using a 405 and 488nm laser line respectively. Fluorescence was detected between 440 and 460nm (calcofluor white/chloroplasts) and between 500 and 550nm (FDA). With the 488-laser line, transmitted light was also collected. Z-stacks (512×512 pixels, pixel size 0.62 µm) were taken at 1 µm intervals of the bottom 12 µm of the cells. For Q-Warg analysis, maximum-intensity Z-projections were used for the fluorescence channels, and for the transmitted light channel, the topmost Z-slice was used.

### Computational workflow

The analysis workflow is first based on 1) cell segmentation on the bright field image using CellPose, 2) an ImageJ macro to extract quantitative information from the images, 3) a R script to analyse and represent the quantitative data. The procedure described here can be downloaded from GitHub (https://github.com/VergerLab/Q-Warg) and Method S3-S4. The overall concept and procedure are described in the main text (Figure 1) and a detailed step-by-step description of the procedure is available in the user guide available in the GitHub repository and as Method S2.

#### Cell segmentation

Cell segmentation is done on the brightfield image with the generalist model cyto3 (Stringer and Pachitariu, 2024). Used settings were default except the cell diameter (40 pixels) and the flow threshold (0.2). The label image generated from the segmentation were saved as PNG. Details on how to use CellPose can be found in Q-Warg user guide (Method S1).

#### Quantification

We developed a batch processing imageJ macro using Fiji (Schindelin *et al*., 2012) to extract quantitative information from segmented label images and the corresponding cell wall and viability fluorescence images (CWRegenerationQuantification.ijm; Method S3). The macro splits the multichannel image (brightfield, CW, Viability) and saves individual images as tif as well as brightfield images with cell segmentation (contours) overlap. The macro uses the ImageJ plugin MorpholibJ (Legland, Arganda-Carreras and Andrey, 2016) to extract morphometric information (*Plugins>MorphoLibJ>Analyze>Analyze Regions*) on individual cells from the label image, as well as fluorescence intensity (*Plugins>MorphoLibJ>Analyze>Intensity Measurements 2D/3D*) on the CW and Viability images for individual cells using the label image.

#### Data analysis

We developed an R script in a markdown notebook (QWARG.Rmd; Method S4) usable in RStudio (https://posit.co/download/rstudio-desktop/) to analyse the data issued from the quantification. Data tables generated by the CWRegenerationQuantification.ijm (additional file 4) macro were combined and organized to be treated all at once (tidyverse (Wickham et al., 2019)). Automatic thresholding was computed to discriminate dead and living cells using the autothreshold library (Nolan [aut *et al*., 2023) and applied in a shiny app using plotly to render interactive plots (shiny (Chang *et al*., 2024; *Create Interactive Web Graphics via ‘plotly.js’ [R package plotly version 4.10.4]*, 2024)). The plots were done with the ggplot2 library. Another Shiny app helps to check the results of the thresholding and cell segmentation by linking interactive plots (plotly) to the microscopy images (magick (Ooms [aut and cre, 2024)). All information about other libraries used can be found in the user guide (Method S2).

#### Statistical analysis

We performed statistical analysis within the R notebook using the agricolae library (Mendiburu, 2023). One-way Analysis of Variance (ANOVA) followed by Tukey’s Honestly Significant Difference (HSD) test were performed for multiple comparisons of means for CW staining intensity and cell size plots. Chi-square test of independence (significant if p < 0.05) was performed, followed by pairwise comparisons using the pairwise.prop.test function with Bonferroni correction for multiple testing.

### Osmotic shock in microfluidic chips

#### Chip fabrication

The microfluidic chips were fabricated according to the method detailed in (Sakai et al., 2019). Starting from a SU8-on-silicon master mold (IPGG, Paris), a 9:1 PDMS (Silgard^TM^ 184 Silicone Elastomer, Dow) mixture is poured on top of it, degassed for two hours in a vacuum chamber and cured at 70°C overnight. The chips are then individually cut with a razor blade, and inlets and outlets are created using a biopsy puncher (1mm diameter, pfm medical). PDMS chips are then rinsed with isopropanol and stuck to a glass bottom petridish (WPI Fluorodish P35-100) in a cleanroom (Nanolab, Umeå) using an oxygen plasma cleaner (Plasma cleaning system ATTO, 0.35mbar, 36sec).

#### Chip preparation

Before loading the cells, a flow of EtOH 96% was performed to sterilize the tubing (0.022” ID x 0.042” OD, PTFE, Masterflex) and the chip but also to reduce the apparition of bubbles in the setup. Then a filtered (0.2µm filter) of pluronic F-127 0.02% solution in water was added overnight inside the chip. The pluronic solution was replaced with medium before loading the cells.

#### Osmotic shock

Four days post-extraction, the cell suspension was stained as explained previously with calcofluor and FDA. The cells were loaded at 100µL/min with a DIY syringe pump (Baas and Saggiomo, 2021) until most of the traps contained a cell. MilliQ water was then flown into the chip 50µL/min. Short-term bright field timelapses of the chip are recorded with the Leica Dmi8 microscope.

#### Time lapse experiment

Protoplasts were cultured in FluoroDish (FD35, World Precision Instruments) coated with poly-lysine to maintain cells in place. FluoroDish glass bottom was covered by a solution of poly-L-lysine 0.1% (P8920, Sigma-Aldrich) for 5min at room temperature. The liquid was removed as much as possible with a pipette and quickly dried with an air gun. The plates were dried in an oven at 60°C for one hour. A 3D printed microscope Openflexure (high resolution version, (Collins *et al*., 2020)) was used to image the culture in brightfield, without staining. Images were taken every 15 minutes with white LED illumination for the duration of the experiments. Cells were cultured at room temperature (20-22°C).

### Fluorescence-Activated Cell Sorting

Before analysis and sorting, the cells were washed from the culture medium and put into W5 modified (W5m). The cell suspension was transferred into a Falcon tube 15mL and centrifuged at 200rcf (swing out rotor centrifuge) for 5min, RT. W5m contains only 2mM calcium (instead of 125mM) to avoid the precipitation with the sheath fluid (70% FACS Flow; BD Bioscience, San Jose, CA, USA) used during sorting and 0,1% BSA to prevent sticking of the cells to the plastic walls of the sorting plate.

The FACS BD Aria III equipped with four lasers: violet (405 nm), blue (488 nm), yellow green (561 nm) and red (633 nm) lasers (BD Biosciences, San Jose, CA, USA). BD FACSDiva software version 7.0 was used for handling of the cytometer and respective data analysis. For each condition, the non-stained sample was subjected for analysis as a negative control. Cell suspension was stained 30min with 1:1000 dilution CarboTrace 680 for CW staining and FDA (stock solution 5mg/mL, predilution: 1.6µL for 1mL medium, then add 1.6µL of the predilution for 1mL cell suspension).

Protoplast analysis and sorting: Filtered protoplast suspension through 70µm Flowmi Cell strainer (SP Bel-Art, Wayne, NJ, USA) was loaded in the cell sorter (room temperature, mild agitation of 100 rpm) and forced through the cuvette in a single file stream, where laser lights intercepted the stream at the sample interrogation point. After passing through the cuvette, the stream entered the integrated 100 µm nozzle tip, where the drop drive broke the stream into the droplets for sorting. The forward scatter (FSC) of light was initially filtered through a 1.5 neutral density filter and then perceived by a photodiode detector with a 488/10 bandpass filter. Light scatter information was collected for identification of protoplast population and design sorting strategy (Figure S2).

Different protoplast populations based on fluorescent properties given by used fluorophores were sorted for microscopy analysis into 500µL of CWR medium in a 24-well plate. To remove the debris, all events smaller than 16µm were disregarded. The cells were sorted based on their viability (FDA staining) and their CW staining (CarboTrace 680). We used CarboTrace 680 instead of Calcofluor for CW staining to limit the toxicity applied on the cells. To identify and subsequently exclude doublets (the droplets containing more than one protoplast), the ratio of fluorescence signal width to the respective area was measured (Suda *et al*., 2007).

## Data statement

All the data raw generated for this study along with the processed data and quantification has been deposited at Zenodo and is publicly available as of the date of publication (10.5281/zenodo.14971281).

The code for the Q-Warg workflow is publicly available on github (github.com/VergerLab/Q-Warg) and the specific version used for this study has been archived at Zenodo and is publicly available as of the date of publication at (10.5281/zenodo.14973382).

Requests for further information and resources should be directed to and will be fulfilled by the corresponding author, Stephane Verger

## Acknowledgements

We thank Åsa Strand and Xu Jin for providing the root cell culture material and initial technical help. We thank Adrien Heymans for helping with the statistical part of the R script. We thank Pauline Durand and Cyril Grandjean, Antoine Chevallier and Isaty Melogno, Mateusz Majda and Lukas Hoermayer for discussions about protoplast extraction and culture optimization. We thank the facilities and technical assistance of the Umeå Plant Science Center (UPSC), the microscopy facility and the growth facility. We thank the NanoLab (Physics department, Umeå University) for the technical resources for microfluidic chips fabrication. This work benefited from the technical contribution of the joint service unit CNRS UAR 3750 (Paris, France) at IPGG. This work was supported by grants from the Swedish research council (VR, 2020–03974), Novo Nordisk foundation (NNF21OC0067282), Åforsk foundation (20-502), Knut and Alice Wallenberg Foundation (KAW 2022.0029) to S.V. R.F. is financially supported by the TopSector TKI Horticulture & Starting Materials, the Netherlands, through the project TU202312. J.S. and P.S. are funded by the European Research Council project Catch, project number 101000981. I.A. and V.S. were supported by grants from the Swedish Research Council (VR 2021-04938) and Kempestiftelserna (SMK21-0041). This work was also supported by Umeå Plant Science Centre with grants from the Knut and Alice Wallenberg Foundation (KAW 2016.0352 and KAW 2020.0240), the Swedish Governmental Agency for Innovation Systems (VINNOVA 2016–00504) and Bio4Energy, a Strategic Research Environment supported through the Swedish Government’s Strategic Research Area initiative.

## Short legends for supporting information

### Supporting figures

Figure S1: Screening for best CW recovery medium with different hormones

Figure S2: Cell selection with Fluorescence Activated Cell Sorting

Figure S3: Additional information from the osmotic change

Figure S4: Additional information for figure 4

### Supporting movie

Movie S1: Longterm imaging of SPCs culture. Timelapse over five days of SPCs culture in glass bottom petri dish and imaged with a 3D-printed microscope (Openflexure) every 15 minutes. The petri dish is coated with polylysine to maintain cells at the same position. Black arrow presents an example of a dead cell. White arrow highlights an example of a living cell. The close-up view comes from the same experiment and shows a living cell.

### Supporting tables

Table S1: Optimization points for protoplast extraction and culture

Table S2: Selection of protoplasting and CW recovery protocols

### Supporting data

Data S1: Excel file with all data shown in principal and supplementary figures.

### Supporting methods

Method S1: Protocol for protoplast extraction and sample preparation for Q-Warg

Method S2: Q-Warg User-guide

Method S3: Macro Fiji for parameters measurements CWRegenerationQuantification.ijm

Method S4: R notebook for data analysis and checking Q-Warg.Rmd

## References

Anderson, C.T. and Kieber, J.J. (2020) ‘Dynamic Construction, Perception, and Remodeling of Plant Cell Walls’, Annual Review of Plant Biology, 71(Volume 71, 2020), pp. 39–69. Available at: 10.1146/annurev-arplant-081519-035846.

Anderson, C.T. and Pelloux, J. (2025) ‘The Dynamics, Degradation, and Afterlives of Pectins: Influences on Cell Wall Assembly and Structure, Plant Development and Physiology, Agronomy, and Biotechnology’, Annual Review of Plant Biology [Preprint]. Available at: 10.1146/annurev-arplant-083023-034055.

Antoniadi, I. et al. (2022) ‘Fluorescence activated cell sorting-A selective tool for plant cell isolation and analysis’, Cytometry. Part A: The Journal of the International Society for Analytical Cytology, 101(9), pp. 725–736. Available at: 10.1002/cyto.a.24461.

Atakhani, A., Bogdziewiez, L. and Verger, S. (2022) ‘Characterising the mechanics of cell–cell adhesion in plants’, Quantitative Plant Biology, 3, p. e2. Available at: 10.1017/qpb.2021.16.

Baas, S. and Saggiomo, V. (2021) ‘Ender3 3D printer kit transformed into open, programmable syringe pump set’, HardwareX, 10, p. e00219. Available at: 10.1016/j.ohx.2021.e00219.

Barciszewski, J., Massino, F. and Clark, B.F.C. (2007) ‘Kinetin—A multiactive molecule’, International Journal of Biological Macromolecules, 40(3), pp. 182–192. Available at: 10.1016/j.ijbiomac.2006.06.024.

Bidhendi, A.J. and Geitmann, A. (2016) ‘Relating the mechanics of the primary plant cell wall to morphogenesis’, Journal of Experimental Botany, 67(2), pp. 449–461. Available at: 10.1093/jxb/erv535.

Cano, E.L., Moguerza, J.M. and Redchuk, A. (2012) ‘Design of Experiments with R’, in E.L. Cano, J.M. Moguerza, and A. Redchuk (eds) Six Sigma with R: Statistical Engineering for Process Improvement. New York, NY: Springer, pp. 197–215. Available at: 10.1007/978-1-4614-3652-2_11.

Chang, W. et al. (2024) ‘shiny: Web Application Framework for R’. Available at: https://cran.r-project.org/web/packages/shiny/ (Accessed: 19 February 2025).

Chupeau, M.-C. et al. (2013) ‘Characterization of the Early Events Leading to Totipotency in an Arabidopsis Protoplast Liquid Culture by Temporal Transcript Profiling’, The Plant Cell, 25(7), pp. 2444–2463. Available at: 10.1105/tpc.113.109538.

Collins, J.T. et al. (2020) ‘Robotic microscopy for everyone: the OpenFlexure microscope’, Biomedical Optics Express, 11(5), pp. 2447–2460. Available at: 10.1364/BOE.385729.

Cosgrove, D.J. (2024) ‘Structure and growth of plant cell walls’, Nature Reviews Molecular Cell Biology, 25(5), pp. 340–358. Available at: 10.1038/s41580-023-00691-y.

Create Interactive Web Graphics via ‘plotly.js’ [R package plotly version 4.10.4] (2024). Comprehensive R Archive Network (CRAN). Available at: https://CRAN.R-project.org/package=plotly (Accessed: 19 February 2025).

Damm, B. and Willmitzer, L. (1988) ‘Regeneration of fertile plants from protoplasts of different Arabidopsis thaliana genotypes’, Molecular and General Genetics MGG, 213(1), pp. 15–20. Available at: 10.1007/BF00333392.

Eine Methode zur Isolierung lebender Protoplasten / von John Af Klercker (1892). Available at: http://sammlungen.ub.uni-frankfurt.de/botanik/4449859 (Accessed: 6 February 2025).

Fruleux, A., Verger, S. and Boudaoud, A. (2019) ‘Feeling Stressed or Strained? A Biophysical Model for Cell Wall Mechanosensing in Plants’, Frontiers in Plant Science, 10. Available at: 10.3389/fpls.2019.00757.

Gilliard, G. et al. (2021) ‘Protoplast: A Valuable Toolbox to Investigate Plant Stress Perception and Response’, Frontiers in Plant Science, 12. Available at: 10.3389/fpls.2021.749581.

Hoffmann, N. et al. (2021) ‘Subcellular coordination of plant cell wall synthesis’, Developmental Cell, 56(7), pp. 933–948. Available at: 10.1016/j.devcel.2021.03.004.

Jayachandran, D. et al. (2023) ‘Engineering and characterization of carbohydrate-binding modules for imaging cellulose fibrils biosynthesis in plant protoplasts’, Biotechnology and Bioengineering, 120(8), pp. 2253–2268. Available at: 10.1002/bit.28484.

Jeong, Y.Y. et al. (2021) ‘Optimization of protoplast regeneration in the model plant Arabidopsis thaliana’, Plant Methods, 17(1), p. 21. Available at: 10.1186/s13007-021-00720-x.

Jiang, F., Zhu, J. and Liu, H.-L. (2013) ‘Protoplasts: a useful research system for plant cell biology, especially dedifferentiation’, Protoplasma, 250(6), pp. 1231–1238. Available at: 10.1007/s00709-013-0513-z.

Krasteva, G., Georgiev, V. and Pavlov, A. (2021) ‘Recent applications of plant cell culture technology in cosmetics and foods’, Engineering in Life Sciences, 21(3–4), pp. 68–76. Available at: 10.1002/elsc.202000078.

Kuki, H. et al. (2017) ‘Quantitative confocal imaging method for analyzing cellulose dynamics during cell wall regeneration in Arabidopsis mesophyll protoplasts’, Plant Direct, 1(6), p. e00021. Available at: 10.1002/pld3.21.

Kuki, H. et al. (2020) ‘Xyloglucan Is Not Essential for the Formation and Integrity of the Cellulose Network in the Primary Cell Wall Regenerated from Arabidopsis Protoplasts’, Plants, 9(5), p. 629. Available at: 10.3390/plants9050629.

Kumari, I. (2019) ‘A Simple Procedure for Isolation, Culture of Protoplast and Plant Regeneration’, in M. Kumar et al. (eds) In vitro Plant Breeding towards Novel Agronomic Traits. Singapore: Springer Singapore, pp. 263–273. Available at: 10.1007/978-981-32-9824-8_14.

Laforest, L.C. and Nadakuduti, S.S. (2022) ‘Advances in Delivery Mechanisms of CRISPR Gene-Editing Reagents in Plants’, Frontiers in Genome Editing, 4, p. 830178. Available at: 10.3389/fgeed.2022.830178.

Legland, D., Arganda-Carreras, I. and Andrey, P. (2016) ‘MorphoLibJ: integrated library and plugins for mathematical morphology with ImageJ’, Bioinformatics, 32(22), pp. 3532–3534. Available at: 10.1093/bioinformatics/btw413.

Leszczuk, A. et al. (2023) ‘Review: structure and modifications of arabinogalactan proteins (AGPs)’, BMC Plant Biology, 23(1), p. 45. Available at: 10.1186/s12870-023-04066-5.

Masson, J. and Paszkowski, J. (1992) ‘The culture response of Arabidopsis thaliana protoplasts is determined by the growth conditions of donor plants’, The Plant Journal, 2(5), pp. 829–833. Available at: 10.1111/j.1365-313X.1992.tb00153.x.

McKenna, J.F., Tolmie, A.F. and Runions, J. (2014) ‘Across the great divide: the plant cell surface continuum’, Current Opinion in Plant Biology, 22, pp. 132–140. Available at: 10.1016/j.pbi.2014.11.004.

Ménard, D. et al. (2017) ‘Establishment and Utilization of Habituated Cell Suspension Cultures for Hormone-Inducible Xylogenesis’, in M. de Lucas and J.P. Etchhells (eds) Xylem: Methods and Protocols. New York, NY: Springer, pp. 37–57. Available at: 10.1007/978-1-4939-6722-3_4.

Ménard, D. et al. (2024) ‘Inducible Pluripotent Suspension Cell Cultures (iPSCs) to Study Plant Cell Differentiation’, in J. Agusti (ed.) Xylem: Methods and Protocols. New York, NY: Springer US, pp. 171–200. Available at: 10.1007/978-1-0716-3477-6_13.

Mendiburu, F. de (2023) ‘agricolae: Statistical Procedures for Agricultural Research’. Available at: https://cran.r-project.org/web/packages/agricolae/index.html (Accessed: 19 February 2025).

Menges, M. and Murray, J.A.H. (2002) ‘Synchronous Arabidopsis suspension cultures for analysis of cell-cycle gene activity’, The Plant Journal, 30(2), pp. 203–212. Available at: 10.1046/j.1365-313X.2002.01274.x.

Mukundan, N.S., Satyamoorthy, K. and Babu, V.S. (2025) ‘Advancing plant protoplasts: innovative techniques and future prospects’, Plant Biotechnology Reports [Preprint]. Available at: 10.1007/s11816-025-00957-1.

Nagata, T. and Takebe, I. (1970) ‘Cell Wall Regeneration and Cell Division in Isolated Tobacco Mesophyll Protoplasts’, Planta, 92(4), pp. 301–308.

Nolan [aut, R., et al. (2023) ‘autothresholdr: An R Port of the “ImageJ” Plugin “Auto Threshold”’. Available at: https://cran.r-project.org/web/packages/autothresholdr/index.html (Accessed: 19 February 2025).

Oda, Y. et al. (2020) ‘Sucrose starvation induces the degradation of proteins in trans-Golgi network and secretory vesicle cluster in tobacco BY-2 cells’, Bioscience, Biotechnology, and Biochemistry, 84(8), pp. 1652–1666. Available at: 10.1080/09168451.2020.1756736.

Ooms [aut, J. and cre (2024) ‘magick: Advanced Graphics and Image-Processing in R’. Available at: https://cran.r-project.org/web/packages/magick/index.html (Accessed: 19 February 2025).

Paredez, A.R., Somerville, C.R. and Ehrhardt, D.W. (2006) ‘Visualization of Cellulose Synthase Demonstrates Functional Association with Microtubules’, Science, 312(5779), pp. 1491–1495. Available at: 10.1126/science.1126551.

Park, Y.B. and Cosgrove, D.J. (2012) ‘A Revised Architecture of Primary Cell Walls Based on Biomechanical Changes Induced by Substrate-Specific Endoglucanases’, Plant Physiology, 158(4), pp. 1933–1943. Available at: 10.1104/pp.111.192880.

Pasternak, T. et al. (2020) ‘From Single Cell to Plants: Mesophyll Protoplasts as a Versatile System for Investigating Plant Cell Reprogramming’, International Journal of Molecular Sciences, 21(12), p. 4195. Available at: 10.3390/ijms21124195.

Pasternak, T., Paponov, I.A. and Kondratenko, S. (2021) ‘Optimizing Protocols for Arabidopsis Shoot and Root Protoplast Cultivation’, Plants, 10(2), p. 375. Available at: 10.3390/plants10020375.

Peng, L. et al. (2022) ‘Design of experiment techniques for the optimization of chromatographic analysis conditions: A review’, ELECTROPHORESIS, 43(18–19), pp. 1882–1898. Available at: 10.1002/elps.202200072.

Pesquet, E. et al. (2010) ‘The Microtubule-Associated Protein AtMAP70-5 Regulates Secondary Wall Patterning in *Arabidopsis* Wood Cells’, Current Biology, 20(8), pp. 744–749. Available at: 10.1016/j.cub.2010.02.057.

Pesquet, E., Wagner, A. and Grabber, J.H. (2019) ‘Cell culture systems: invaluable tools to investigate lignin formation and cell wall properties’, Current Opinion in Biotechnology, 56, pp. 215–222. Available at: 10.1016/j.copbio.2019.02.001.

Purushotham, P., Ho, R. and Zimmer, J. (2020) ‘Architecture of a catalytically active homotrimeric plant cellulose synthase complex’, Science, 369(6507), pp. 1089–1094. Available at: 10.1126/science.abb2978.

Reed, K.M. and Bargmann, B.O.R. (2021) ‘Protoplast Regeneration and Its Use in New Plant Breeding Technologies’, Frontiers in Genome Editing, 3. Available at: 10.3389/fgeed.2021.734951.

Reyna-Llorens, I., Ferro-Costa, M. and Burgess, S.J. (2023) ‘Plant protoplasts in the age of synthetic biology’, Journal of Experimental Botany, 74(13), pp. 3821–3832. Available at: 10.1093/jxb/erad172.

Sakai, K. et al. (2019) ‘Design of a comprehensive microfluidic and microscopic toolbox for the ultra-wide spatio-temporal study of plant protoplasts development and physiology’, Plant Methods, 15(1), p. 79. Available at: 10.1186/s13007-019-0459-z.

Sakamoto, Y. et al. (2022) ‘Transcriptional activation of auxin biosynthesis drives developmental reprogramming of differentiated cells’, The Plant Cell, 34(11), pp. 4348–4365. Available at: 10.1093/plcell/koac218.

Sapala, A. et al. (2018) ‘Why plants make puzzle cells, and how their shape emerges’, eLife, 7, p. e32794. Available at: 10.7554/eLife.32794.

Schindelin, J. et al. (2012) ‘Fiji: an open-source platform for biological-image analysis’, Nature Methods, 9(7), pp. 676–682. Available at: 10.1038/nmeth.2019.

Schirawski, J., Planchais, S. and Haenni, A.-L. (2000) ‘An improved protocol for the preparation of protoplasts from an established *Arabidopsis thaliana* cell suspension culture and infection with RNA of turnip yellow mosaic tymovirus: a simple and reliable method’, Journal of Virological Methods, 86(1), pp. 85–94. Available at: 10.1016/S0166-0934(99)00173-1.

Stringer, C. and Pachitariu, M. (2024) ‘Cellpose3: one-click image restoration for improved cellular segmentation’. bioRxiv, p. 2024.02.10.579780. Available at: 10.1101/2024.02.10.579780.

Suda, J. et al. (2007) ‘Flow Cytometry and Ploidy: Applications in Plant Systematics, Ecology and Evolutionary Biology’, in Flow Cytometry with Plant Cells. John Wiley & Sons, Ltd, pp. 103–130. Available at: 10.1002/9783527610921.ch5.

Tagawa, S. et al. (2019) ‘Dynamics of structural polysaccharides deposition on the plasma-membrane surface of plant protoplasts during cell wall regeneration’, Journal of Wood Science, 65(1), p. 47. Available at: 10.1186/s10086-019-1826-0.

Tan, L. et al. (2013) ‘An Arabidopsis Cell Wall Proteoglycan Consists of Pectin and Arabinoxylan Covalently Linked to an Arabinogalactan Protein[W]’, The Plant Cell, 25(1), pp. 270–287. Available at: 10.1105/tpc.112.107334.

Van Leene, J. et al. (2011) ‘Isolation of Transcription Factor Complexes from Arabidopsis Cell Suspension Cultures by Tandem Affinity Purification’, in L. Yuan and S.E. Perry (eds) Plant Transcription Factors: Methods and Protocols. Totowa, NJ: Humana Press, pp. 195–218. Available at: 10.1007/978-1-61779-154-3_11.

Verma, A., Verma, M. and Singh, A. (2020) ‘Chapter 14 - Animal tissue culture principles and applications’, in A.S. Verma and A. Singh (eds) Animal Biotechnology (Second Edition). Boston: Academic Press, pp. 269–293. Available at: 10.1016/B978-0-12-811710-1.00012-4.

Wickham, H. et al. (2019) ‘Welcome to the Tidyverse’, Journal of Open Source Software, 4(43), p. 1686. Available at: 10.21105/joss.01686.

Wu, F.-H. et al. (2009) ‘Tape-Arabidopsis Sandwich - a simpler Arabidopsis protoplast isolation method’, Plant Methods, 5(1), p. 16. Available at: 10.1186/1746-4811-5-16.

Xu, Y. et al. (2022) ‘Protoplasts: small cells with big roles in plant biology’, Trends in Plant Science, 27(8), pp. 828–829. Available at: 10.1016/j.tplants.2022.03.010.

Yokoyama, R. et al. (2014) ‘The Biosynthesis and Function of Polysaccharide Components of the Plant Cell Wall’, in Plant Cell Wall Patterning and Cell Shape. John Wiley & Sons, Ltd, pp. 1–34. Available at: 10.1002/9781118647363.ch1.

Yokoyama, R. et al. (2016) ‘Arabidopsis Regenerating Protoplast: A Powerful Model System for Combining the Proteomics of Cell Wall Proteins and the Visualization of Cell Wall Dynamics’, Proteomes, 4(4), p. 34. Available at: 10.3390/proteomes4040034.

Yoo, S.-D., Cho, Y.-H. and Sheen, J. (2007) ‘Arabidopsis mesophyll protoplasts: a versatile cell system for transient gene expression analysis’, Nature Protocols, 2(7), pp. 1565–1572. Available at: 10.1038/nprot.2007.199.

Yue, J.-J. et al. (2021) ‘Protoplasts: From Isolation to CRISPR/Cas Genome Editing Application’, Frontiers in Genome Editing, 3, p. 717017. Available at: 10.3389/fgeed.2021.717017.

Zhai, Z., Jung, H. and Vatamaniuk, O.K. (2009) ‘Isolation of Protoplasts from Tissues of 14-day-old Seedlings of Arabidopsis thaliana’, Journal of Visualized Experiments : JoVE, (30), p. 1149. Available at: 10.3791/1149.

Zhang, S. et al. (2024) ‘Phenotyping of single plant cells on a microfluidic cytometry platform with fluorescent, mechanical, and electrical modules’, The Analyst, 149(17), pp. 4436–4442. Available at: 10.1039/d4an00682h.

