## Supplementary figures S1-S4 for "The Q-Warg Pipeline: A Robust and Versatile Workflow for Quantitative Analysis of Protoplast Culture Conditions"

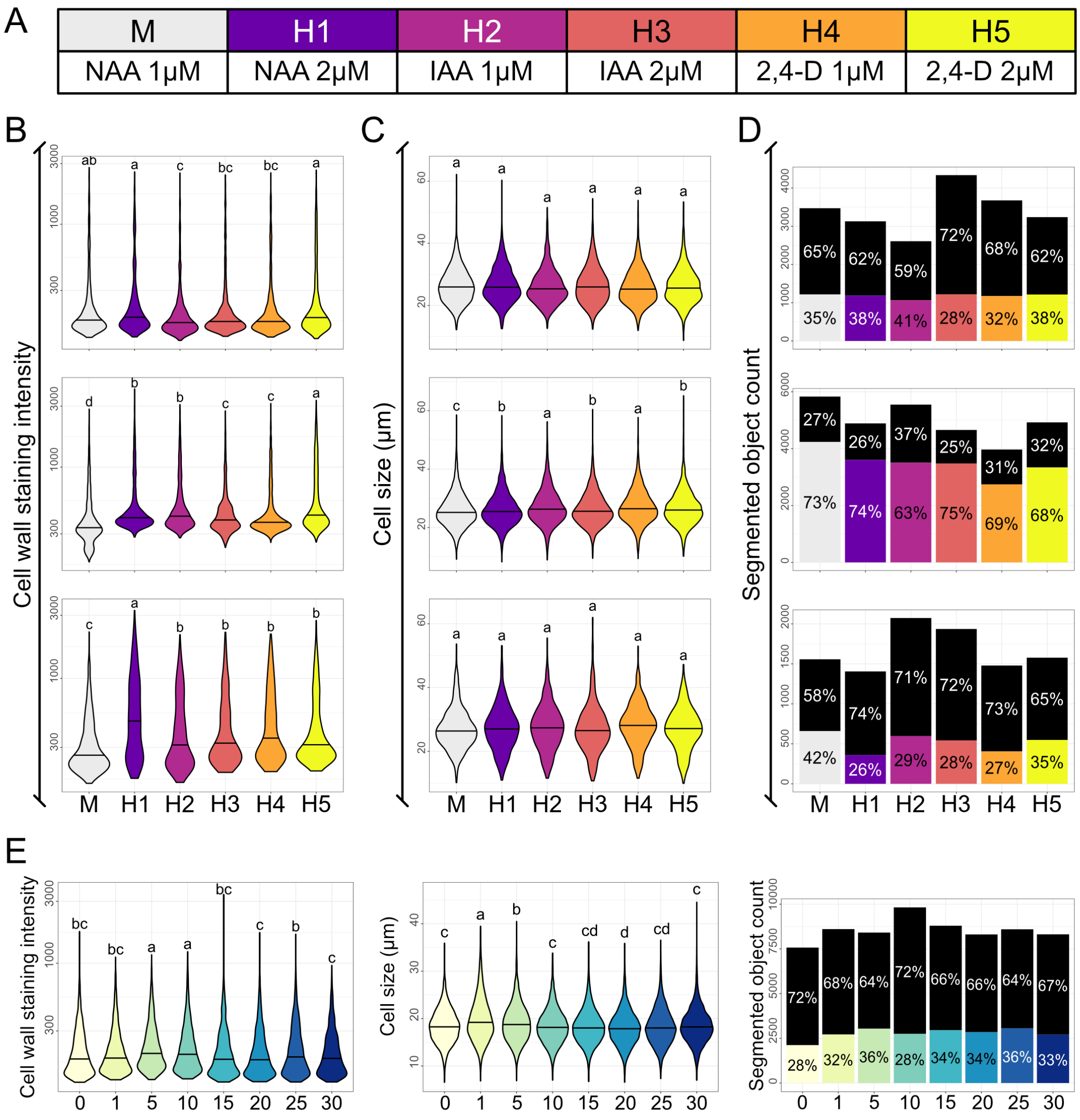


**Figure S1: Screening for best CW recovery medium with different hormones**
(**A**) Table with the tested media composition. All media contain Gamborg B5, MES, Mannitol 0,4M, Glucose 0,05M and the corresponding NAA concentration (**B-C-D**) Plots from data processed with the Q-Warg pipeline on the effect of auxin concentration in the medium for cell wall recovery, cell size and viability, 4 days after protoplasting. Each plot corresponds to one replicate (n=3). (**B**) CW mean intensity after staining with Calcofluor. (**C**) Cell diameter (µm). (**D**) Viability plots displaying the number of segmented objects (including debris, dead cells or living cells). (**E**) Screening of the effect of NAA concentration from 0 to 30µM in S4 medium on cell wall recovery (left), cell size (middle) and viability (right), 4 days after protoplasting. On violin plots, letters describe the statistically significant differences between population determined by one-way ANOVA followed by Tuckey's HSD test ($p < 0,05$). In bar plots, each bar shows the total number of segmented objects, and the proportion of object negative for viability staining (in black) and living cells (in color) and statistical analysis results are in tables Chi-square test of independence (significant if $p < 0.05$) was performed, followed by pairwise comparisons using the pairwise.prop.test function with Bonferroni correction for multiple testing. Results are in additional file 7 (pages FigS1_viability-stats).

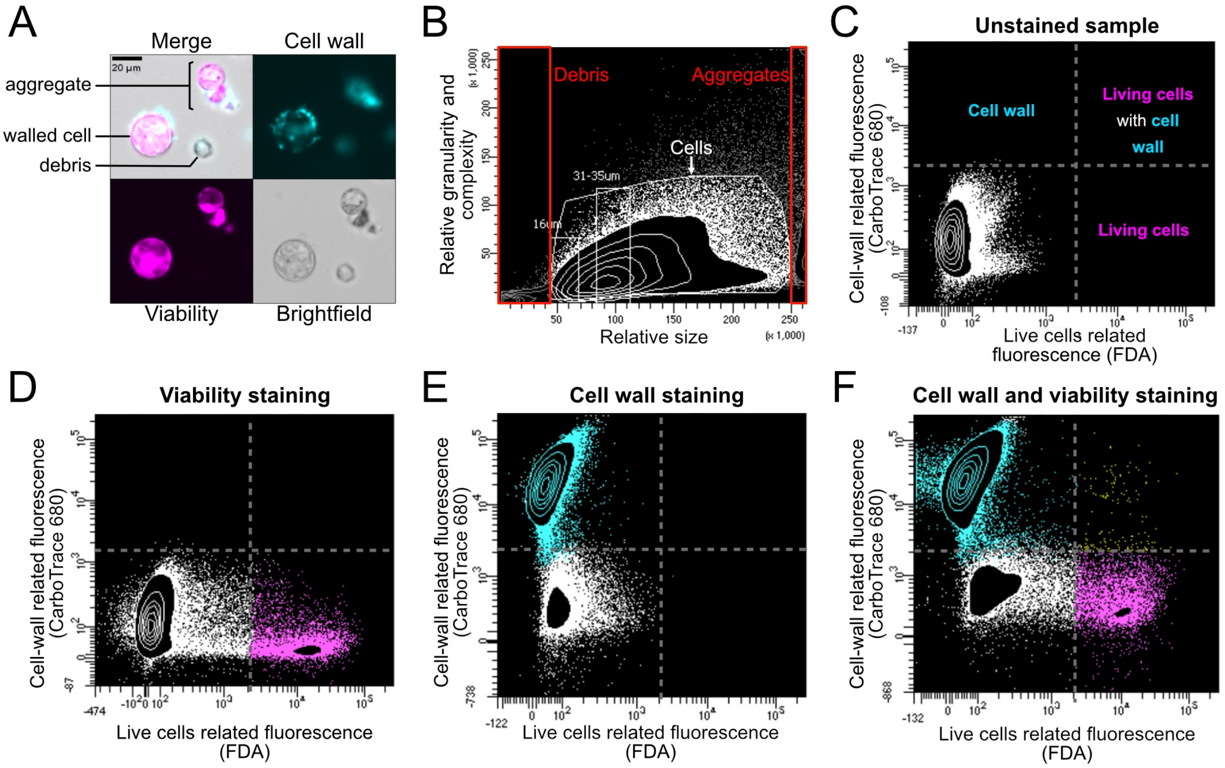


G

| Sub-population | Proportion amongst the total population | Proportion amongst the parent population |
|---|---|---|
| Objects larger than 16µm | 81,5% | / |
| Living cells (Viability +) | 4,1% | 5% of Objects > 16µm |
| Living cells with CW signal | 0,2% | 3,7% of living cells (Viability +) |
| Objects with CW signal | 69,1% | / |

**Figure S2: Cell selection with Fluorescence Activated Cell Sorting**
(**A**) Microscopy images of cell suspension after 3 days of culture in CW recovery medium (here CRRUM: S4 + 5µM NAA). Images highlight the presence of isolated cells, cell debris and cell aggregates as labeled in the panel. CW stained by calcofluor; viability marker is FDA. (**B**) Dot plot generated after FACS sorting. Each event is represented by a point. All events displayed based on their size and granularity. 3 populations are defined: debris, cells and aggregates. (**C-F**) Population of defined cells (based on event size) displayed based on their fluorescence markers Calcofluor white to stain CW and FDA as viability staining. (**C**) Unstained sample. Description of the different gates. (**D**) viability staining only, positive events are magenta (**E**) CW staining only, positive events are cyan. (**F**) Double staining, positive events for both staining are displayed in yellow. (**G**) Proportions of sub-populations in the total events count (total population) and in their parent populations (parent population refers to the larger group from which the sample is drawn; E.g “Living cells” is the parent group of “Living cells with CW signal”). Count done on 100.000 events.

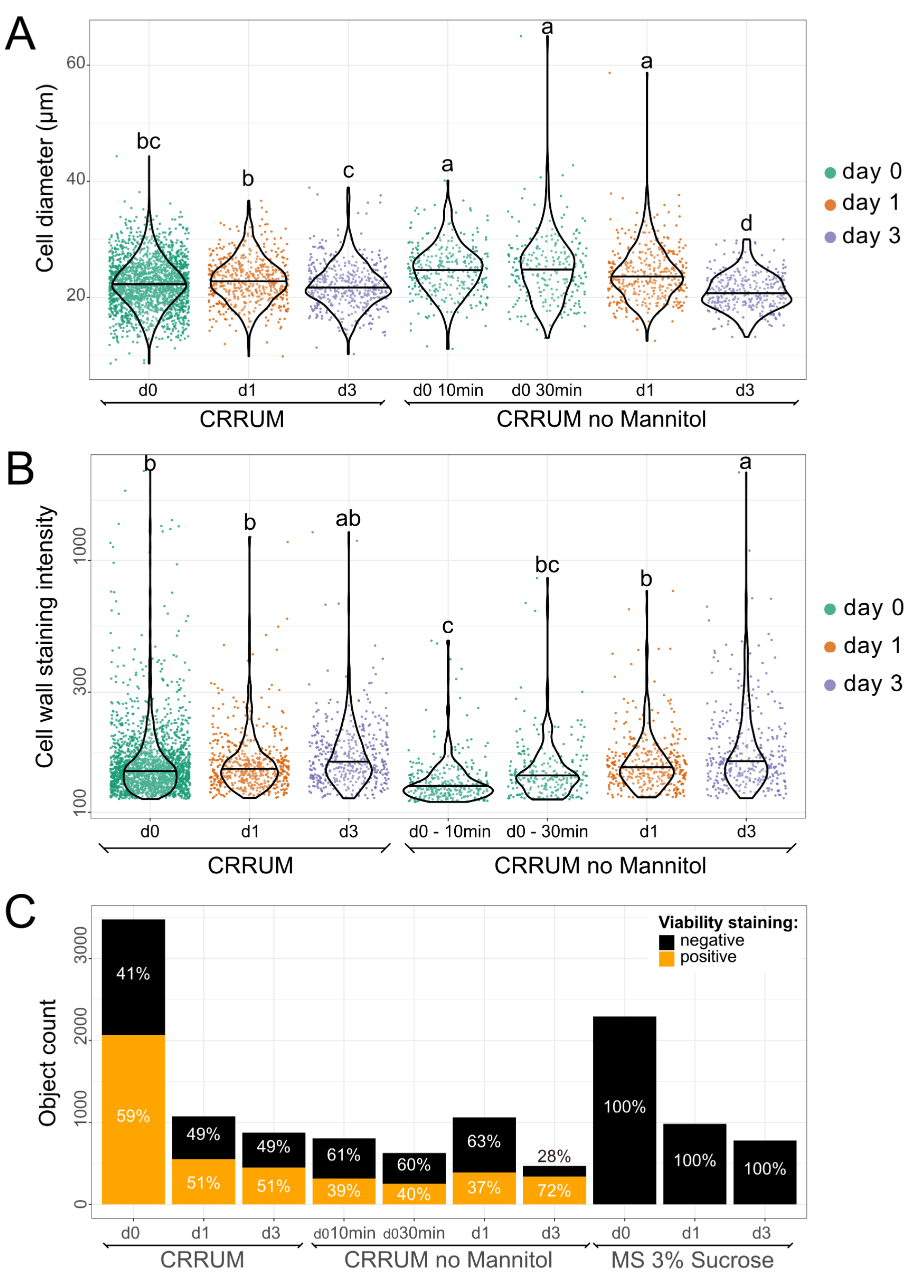


**Figure S3: Additional information from the osmotic change**

(**A**) Violin plot representing the cell size in CRRUM and in CRRUM without mannitol at different time points. Cell diameter increase right after change in medium osmolarity (remove the mannitol). The size is recovered after 3days. (**B**) Violin plot of the CW staining intensity quantification in CRRUM and in CRRUM without mannitol at different time points. (**A-B**) Note that MS 3% sucrose was also used but no cell survived (see C). Letters describe the statistically significant differences between population determined by one-way ANOVA followed by Tuckey's HSD test ($p < 0{,}05$). (**C**) Bar plot of the proportion of living cells after medium change. All cells exposed to MS 3% sucrose are dead. Chi-square test of independence (significant if $p < 0.05$) was performed, followed by pairwise comparisons using the pairwise.prop.test function with Bonferroni correction for multiple testing. Results are in additional file 7 (pages FigS3_viability-stats).

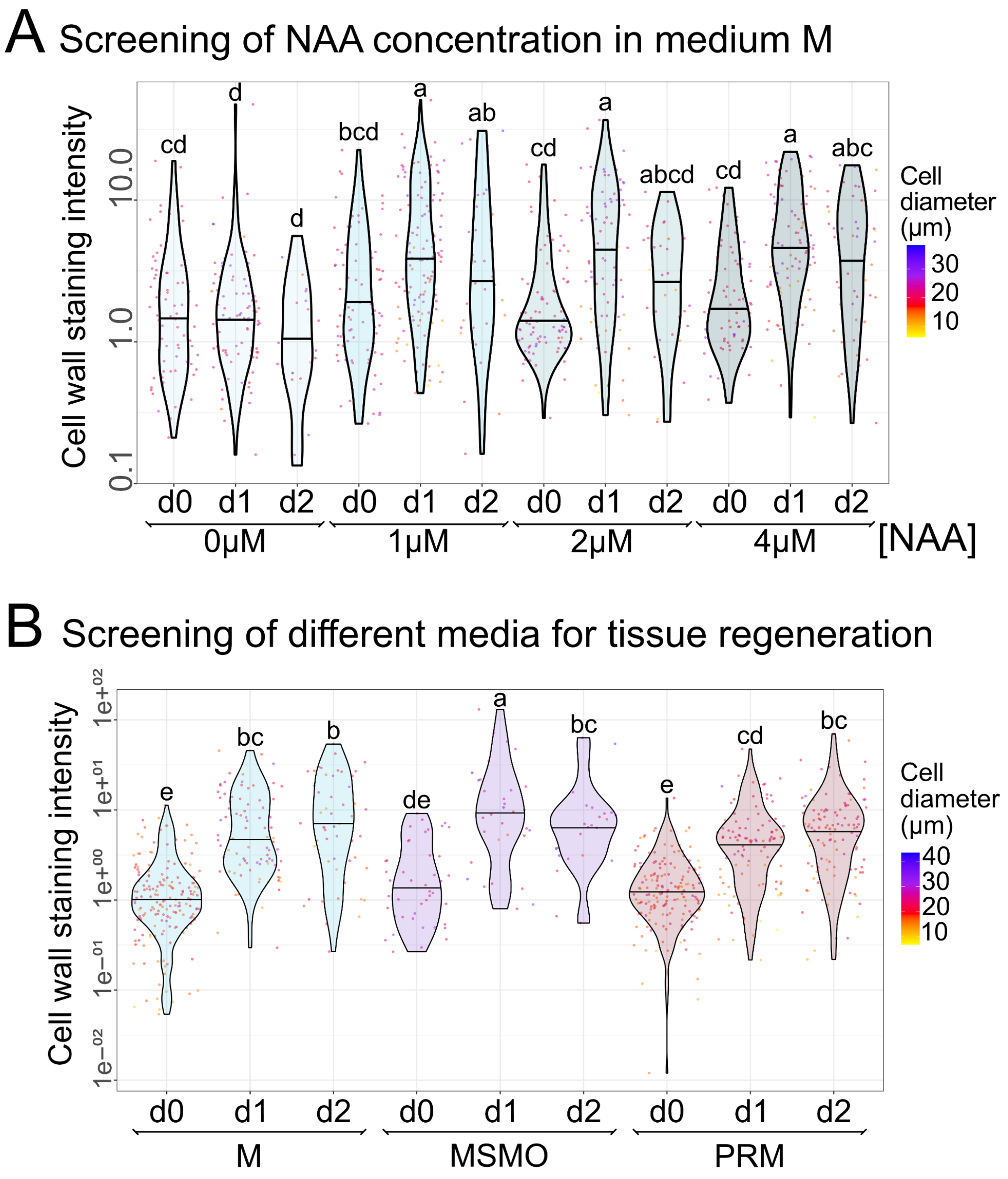


**Figure S4: Additional information for figure 4**

Violin plots of the CW mean intensity after staining with Calcofluor. Letters describe the statistically significant differences between population determined by one-way ANOVA followed by Tuckey's HSD test ($p < 0,05$). (**A**) NAA concentration screening. 4 different concentrations of NAA have been tested in M medium. d0 corresponds to the protoplasting day. (**B**) Medium screening. d0 corresponds to the protoplasting day.
