## Supporting Tables S1-S2 for "The Q-Warg Pipeline: A Robust and Versatile Workflow for Quantitative Analysis of Protoplast Culture Conditions"

**Supplementary table 1: Optimization points for protoplast extraction and culture**
Examples of what can be modified in a protoplast extraction and culture protocol. The right column shows the final parameters selected as optimized following the screening and previous attempts to get single plant cells.

| **Parameters** | | **Examples** | **Have been optimized to** |
|---|---|---|---|
| **Biological material** | Specie | Arabidopsis<br>Tobacco<br>Populus | Arabidopsis roots |
| | Variety, ecotype, accession | Columbia<br>Landsberg<br>Wassilewskija | |
| | Tissue | Leaf<br>Seedling<br>Roots<br>Cell culture | Cell culture |
| | Age | | 4days after passing (refreshing medium) |
| **Protoplasting** | Enzymes | Which ones<br>Concentration | Cellulase R-10 1,5%<br>Macerozyme R-10 0,4% |
| | Digestion medium | Osmolarity<br>Composition | |
| | Conditions during extraction | Shaking<br>Light/dark<br>Temperature<br>Container (6-well plate, flask...) | Shaking<br>Dark<br>24<br>6-well plate non treated |
| | Extraction time | | 4 hours |
| | Tissue preparation | Cuts<br>Full organ<br>Quantity | |
| | Washing steps | How many?<br>Centrifugation speed | 1 with low centrifugation |
| **Culture** | Basis medium | MS<br>Gamborg B5<br>Vitamins | Gamborg B5 |
| | Sugars | For osmolarity<br>For feeding | Mannitol<br>Glucose |
| | Hormones | Auxin<br>Brassinosteroides | NAA |
| | Conditions | Shaking<br>Light/dark<br>Temperature<br>Container (6-well plate, flask...) | No shaking<br>Dark<br>22<br>6-well plate non treated |

**Supplementary table 2: Selection of protoplasting and CW recovery protocols**
Conditions tested prior and for this paper have been chosen by cross-referencing protoplasting protocols and CW recovery media from this list of papers. Note that the list is not an exhaustive list of all protoplasting paper available but the most relevant for our study. The list is classified by alphabetical order of the last author's name.

| Reference | Title | doi |
|---|---|---|
| Aoyagi (2011) | Application of plant protoplasts for the production of useful metabolites | https://doi.org/10.1016/j.bej.2010.05.004 |
| Asai et al. (2000) | Fumonisin B1–Induced Cell Death in Arabidopsis Protoplasts Requires Jasmonate-, Ethylene-, and Salicylate-Dependent Signaling Pathways | https://doi.org/10.1105/tpc.12.10.1823 |
| Brirnbaum et al. (2003) | A Gene Expression Map of the Arabidopsis Root | https://doi.org/10.1126/science.1090022 |
| Jayachandran et al. (2023) | Engineering and characterization of carbohydrate-binding modules for imaging cellulose fibrils biosynthesis in plant protoplasts | https://doi.org/10.1002/bit.28484 |
| Chupeau MC (2013) | Characterization of the Early Events Leading to Totipotency in an Arabidopsis Protoplast Liquid Culture by Temporal Transcript Profiling | https://doi.org/10.1105/tpc.113.109538 |
| Gilliard et al. (2021) | Protoplast: A Valuable Toolbox to Investigate Plant Stress Perception and Response | https://doi.org/10.3389/fpls.2021.749581 |
| Van de Meene (2021) | Interactions between Cellulose and (1,3;1,4)-β-glucans and Arabinoxylans in the Regenerating Wall of Suspension Culture Cells of the Ryegrass Lolium multiflorum | https://doi.org/10.3390/cells10010127 |
| Ehlert et al. (2006) | Two-hybrid protein–protein interaction analysis in Arabidopsis protoplasts: establishment of a heterodimerization map of group C and group S bZIP transcription factors | https://doi.org/10.1111/j.1365-313X.2006.02731.x |
| Chen et al. (2020) | An impedance-coupled microfluidic device for single-cell analysis of primary cell wall regeneration | https://doi.org/10.1016/j.bios.2020.112374 |
| Glanc et al. (2021) | AGC kinases and MAB4/MEL proteins maintain PIN polarity by limiting lateral diffusion in plant cells | https://doi.org/10.1016/j.cub.2021.02.028 |
| Ford (1990) | Plant regeneration from Arabidopsis thaliana protoplasts | https://doi.org/10.1007/BF00820203 |
| Schirawski et al. (2000) | An improved protocol for the preparation of protoplasts from an established Arabidopsis thaliana cell suspension culture and infection with RNA of turnip yellow mosaic tymovirus: a simple and reliable method | https://doi.org/10.1016/S0166-0934(99)00173-1 |
| Hahne et al. (1983) | Wall formation and cell division in fluorescence-labelled plant protoplasts | https://doi.org/10.1007/BF01279812 |
| Planchais et al. (2022) | Protocols for Studying Protein Stability in an Arabidopsis Protoplast Transient Expression System | |
| Van Amstel et al. (1996) | Callose deposition in the primary wall of suspension cells and regenerating protoplasts, and its relationship to patterned cellulose synthesis | https://doi.org/10.1139/b96-128 |
| Tagawa et al. (2019) | Dynamics of structural polysaccharides deposition on the plasma-membrane surface of plant protoplasts during cell wall regeneration | https://doi.org/10.1186/s10086-019-1826-0 |
| Pasternak et al. (2021) | Optimizing Protocols for Arabidopsis Shoot and Root Protoplast Cultivation | https://doi.org/10.3390/plants10020375 |

| | | |
|---|---|---|
| Wu et al. (2009) | Tape-Arabidopsis Sandwich - a simpler Arabidopsis protoplast isolation method | https://doi.org/10.1186/1746-4811-5-16 |
| Wenk et al. (1995) | Large-scale protoplast isolation and regeneration of Arabidopsis thaliana. | https://europepmc.org/article/med/7598898 |
| Sakamoto et al. (2018) | Complete substitution of a secondary cell wall with a primary cell wall in Arabidopsis | https://doi.org/10.1038/s41477-018-0260-4 |
| Gonzàlez-Garcìa et al. (2020) | Fluorescence-Activated Cell Sorting Using the D-Root Device and Optimization for Scarce and/or Non-Accessible Root Cell Populations | https://doi.org/10.3390/plants9040499 |
| Yokoyama et al. (2000) | Functional Diversity of Xyloglucan-Related Proteins and its Implications in the Cell Wall Dynamics in Plants | https://doi.org/10.1055/s-2000-16643 |
| Hye-Kyoung Kwon et al. (2005) | A Proteomic Approach to Apoplastic Proteins Involved in Cell Wall Regeneration in Protoplasts of Arabidopsis Suspension-cultured Cells | https://doi.org/10.1093/pcp/pci089 |
| Yokoyama et al. (2016) | Arabidopsis Regenerating Protoplast: A Powerful Model System for Combining the Proteomics of Cell Wall Proteins and the Visualization of Cell Wall Dynamics | https://doi.org/10.3390/proteomes4040034 |
| Hiroaki Kuki et al. (2017) | Quantitative confocal imaging method for analyzing cellulose dynamics during cell wall regeneration in Arabidopsis mesophyll protoplasts | https://doi.org/10.1002/pld3.21 |
| Hiroaki Kuki et al. (2020) | Xyloglucan Is Not Essential for the Formation and Integrity of the Cellulose Network in the Primary Cell Wall Regenerated from Arabidopsis Protoplasts | https://doi.org/10.3390/plants9050629 |
| Masson et al. (1992) | The culture response of Arabidopsis thaliana protoplasts is determined by the growth conditions of donor plants | https://doi.org/10.1111/j.1365-313X.1992.tb00153.x |
| Karesch et al. (1991) | Direct gene transfer to protoplasts of Arabidopsis thaliana | https://doi.org/10.1007/BF00232335 |
| Barnes et al. (2019) | An Arabidopsis protoplast isolation method reduces cytosolic acidification and activation of the chloroplast stress sensor SENSITIVE TO FREEZING 2 | https://doi.org/10.1080/15592324.2019.1629270 |
| Jeong et al. (2021) | Optimization of protoplast regeneration in the model plant Arabidopsis thaliana | https://doi.org/10.1186/s13007-021-00720-x |
| Kovtun et al. (2000) | Functional analysis of oxidative stress-activated mitogen-activated protein kinase cascade in plants | https://doi.org/10.1073/pnas.97.6.2940 |
| Sheen (2001) | Signal Transduction in Maize and Arabidopsis Mesophyll Protoplasts | https://doi.org/10.1104/pp.010820 |
| Yoo et al. (2007) | Arabidopsis mesophyll protoplasts: a versatile cell system for transient gene expression analysis | https://doi.org/10.1038/nprot.2007.199 |
| Shafi et al. (2019) | Ectopic expression of SOD and APX genes in Arabidopsis alters metabolic pools and genes related to secondary cell wall cellulose biosynthesis and improve salt tolerance | https://doi.org/10.1007/s11033-019-04648-3 |
| Sakamoto et al. (2022) | Transcriptional activation of auxin biosynthesis drives developmental reprogramming of differentiated cells | https://doi.org/10.1093/plcell/koac218 |
| Mathur et al. (1995) | A simple method for isolation, liquid culture, transformation and regeneration of Arabidopsis thaliana protoplasts | https://doi.org/10.1007/BF00233637 |
| Nagata et al. (1970) | Cell wall regeneration and cell division in isolated tobacco mesophyll protoplasts | https://doi.org/10.1007/BF00385097 |

| | | |
|---|---|---|
| Steffen Abel and Athanasios Theologis (1994) | Transient transformation of Arabidopsis leaf protoplasts: a versatile experimental system to study gene expression | https://doi.org/10.1111/j.1365-313X.1994.00421.x |
| Vasil & Vasil (1972) | Totipotency and embryogenesis in plant cell and tissue cultures | https://doi.org/10.1007/BF02619487 |
| Raghupathy et al. (2006) | Transfection of Arabidopsis protoplasts with a Plum pox virus (PPV) infectious clone for studying early molecular events associated with PPV infection | https://doi.org/10.1016/j.jviromet.2006.05.009 |
| Cole et al. (2013) | Live imaging of DORNRÖSCHEN and DORNRÖSCHEN-LIKE promoter activity reveals dynamic changes in cell identity at the microcallus surface of Arabidopsis embryonic suspensions | https://doi.org/10.1007/s00299-012-1339-4 |
| Damm et al. (1988) | Regeneration of fertile plants from protoplasts of different Arabidopsis thaliana genotypes | https://doi.org/10.1007/BF00333392 |
| Damm et al. (1989) | Efficient transformation of *Arabidopsis thaliana* using direct gene transfer to protoplasts | https://doi.org/10.1007/BF00330935 |
