## Supporting method S1 for "The Q-Warg Pipeline: A Robust and Versatile Workflow for Quantitative Analysis of Protoplast Culture Conditions"

### Protoplast extraction and Cell Wall regeneration optimization

*Screening CW* regeneration – *protocol*

### Introduction

#### Aim of the protocol and Q-Warg pipeline

This protocol describes how to optimize the cell wall (CW) regeneration of single plant cells after protoplasting. If getting protoplasts is relatively straightforward, keeping the cells alive in suspension as well as the cell wall regeneration step is challenging. It appears that the medium used for CW regeneration is also dependent on the lab and various parameters can be modified.

Here we propose a screening approach of media to quickly optimize the CW regeneration step. The protocol contains the steps occurring after any protoplast extraction procedure: how to handle and culture protoplasts, and imaging to be used in the Q-Warg pipeline (Method S2 (user guide)).

#### Procedure steps


1- Protoplast extraction
2- Culture of protoplasts
    a. Preparing the media to test
    b. Culture for 4 days
3- Imaging
    a. Staining and mounting
    b. Imaging
4- Q-Warg analysis


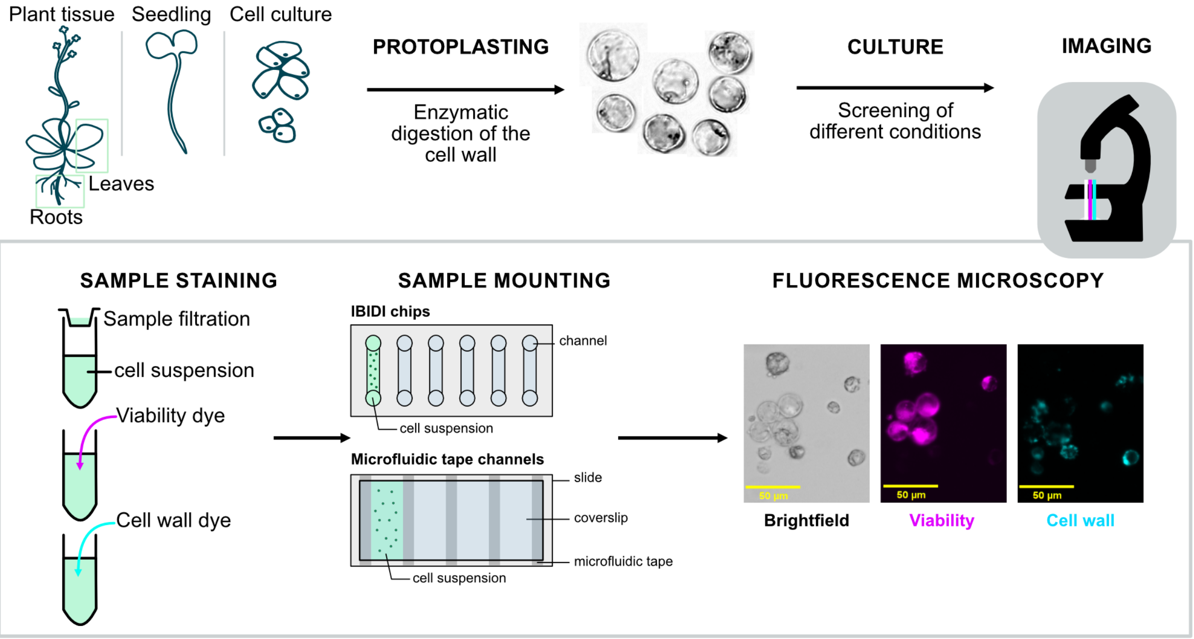

### I - Protoplast extraction

Numerous protocols exist to extract protoplasts from different plant species and/or tissues. In the Table S2 is a list of most used protoplasting protocols.

Here are the detailed protocols for protoplast extraction used in the Q-Warg paper.

##### From root cell culture (UPSC)

The protocol is an adaptation of the protocol from (Yoo, Cho, and Sheen 2007). All solutions containing mannitol were prepared fresh.

- Prepare protoplasting buffer (PB): MES 20mM, mannitol 0,4M, KCl 20mM, $CaCl_2$ 10mM, 0,1% BSA and filled up to the wanted volume with MilliQ water. Adjust pH with KOH to 5,7.
- For 30mL of cell culture, prepare 20mL of enzymatic solution:
  - Warm up PB to 55°C on stirring plate
  - Add Cellulase R-10 1,5% (C8001, Duchefa) and Macerozyme R-10 0,4% (M8002, Duchefa) in PB.
  - Let the solution 5 min at 55°C with agitation
  - Cool it to room temperature and sterilize with syringe filter 0,2µm before use.
- Centrifugate 30mL of 4 days-old cell culture 200g for 3min at room temperature (swing-out rotor centrifuge) in falcon 50mL and remove the supernatant before adding the enzymatic solution.
- Gently mix the cells with the enzymatic solution by flipping the falcon 50mL.
- Transfer the suspension to non-treated 6-well plates sealed with Parafilm. Incubation in the dark for 4 hours under rotation (120rpm) at 24°C.
- During the extraction time, prepare the washing solution (W5): MES 2mM, NaCl 154mM, CaCl2 125mM, KCl 5mM. Adjust pH with KOH to 5,7. W5 can be sterilized by autoclaving or filtration 0,2µm.
- To stop the enzymatic digestion, add the same volume of W5 in each well and place the suspension under rotation 30rpm for 5min, at room temperature in the dark.

*From this step handle very gently the protoplast suspension. Using sterile plastic Pasteur pipettes is less detrimental for the protoplasts as the pressure on them is less important. If you prefer the use of P1000, cut the tip extremity.*

- Filter the protoplast solution with a 70µm cell strainer into a 50mL falcon tube. Keep the falcon with an angle so the solution does not drop but follows the wall of the falcon tube.

*An angled falcon holder can be 3D printed. STL file can be found here:*
*https://www.printables.com/model/785079-falcon-holder-tilt-40degree-angle*

- Centrifuge at 200g for 3min at RT. Remove the supernatant.
- Re-suspend the protoplasts into 5mL of the cell wall regeneration medium.
- Count the cells using a haemocyter (e.g.: Bürker chamber) and adjust the cell concentration to $10^5$ cells/mL.

*If several media are tested at the same time. The protoplasts suspension should be split before centrifugation to avoid multiple centrifugation steps.*

##### From seedlings

Protocol is adapted from (Zhai, Jung, and Vatamaniuk 2009). All solutions were filter sterilized using a 0,2µm syringe filter.

- Prepare TVL solution: sorbitol 0,3M and CaCl2 50mM.
- Transfer about 2g of 18-day-old whole seedlings to a petri dish containing 15 mL TVL solution and were cut into small pieces.
- Prepare *Seedling* enzymatic solution: MES 10mM, sucrose 0.5M, CaCl2 20mM, KCl 40 mM, cellulase R-10 1,0% and macerozyme R-10 1,0% (Duchefa).
- Transfer the macerated tissue to a 250 mL Erlenmeyer and add 20 mL of *seedling* enzymatic solution. Close the Erlenmeyer with parafilm and covered with aluminum foil and incubate in the dark while shaking (35 rpm, 22°C, 16h).
- Prepare W5+ solution: glucose 5,6mM and additional KCl 6mM dissolved in W5 (see root cell culture extraction protocol).
- Filter the protoplast suspension through a 40µm nylon basket, pre-wet in W5+ solution, and divide it equally over 2 50mL tubes.
- Overlay 10mL W5+ onto the protoplast suspension and centrifugate at 100g for 7 min.
- Collect 2 aliquots of 6mL supernatant and transfer to a 12mL round-bottom tube, twice per 50 mL tube (4 aliquots total).
- To each tube, add 6mL W5+ and centrifugate at 60g for 5 min.
- Resuspend the pellet in 13mL W5+, and again centrifugate at 60g for 5 min.
- Final pellet is resuspended in 1mL W5+.
- Protoplast density was adjusted to $0{,}8*10^6$ cells/mL and 200µL of suspension was transferred to each well of an Ibidi µ-Slide (8 Well high, Polymer Coverslip, Uncoated).

##### Notes and tips about protoplasting:

6-well plates should be non-treated (specified when bought). The protoplasts are very attracted to the plastic especially if the plates are treated. With non-treated plates, we can recover most of the protoplasts extracted. Most of the commercially available plates are treated as it is primarily used for animal cells.

To handle the protoplasts, we need to be very gentle. To avoid destroying them by putting too much pressure on them:

- Use plastic Pasteur pipettes
- Cut the P1000 tips
- Do as few centrifugation steps as possible

At lower concentration than $10^5$cells/mL, we observed that CW regeneration is more difficult and most of the cells die. Yet, at higher concentrations, the cells are happier and undergo CW regeneration. Conclusion: cells would rather be packed than dispersed.

### II - Culture

- Culture of protoplasts can be done in 6-well non treated plates.
- Keep the protoplasts in CW regeneration medium in the dark at 22°C without shaking.

If a change of medium is required:

- Centrifuge at 200g for 3min at RT. Remove the supernatant.
- Re-suspend the protoplasts into 5mL of the cell wall regeneration medium.
- Count the cells using a haemocyter (e.g.: Bürker chamber) and adjust the cell concentration to $10^5$ cells/mL.

*During the first week, the shaking of the suspension should be avoided to increase cell viability and CW regeneration. Yet, it has **not** been investigated if it is necessary to restart shaking the cell suspension later.*

### III - Imaging

The imaging was done with an epifluorescence microscope.

- After 4days in culture, the cells are filtered with 70µm nylon filter to remove the big clamps of cells and separate them.
- Staining: (A)
  - put 1mL of cell suspension into a 1.5mL Eppendorf tube
  - add 1,6µL of FDA 5mg/mL staining for viability staining 4min before imaging (no washing)
  - add 10µL of calcofluor (stock solution) as cell wall staining just before imaging.

*FDA staining protocol: IBIDI application note 33*
*https://ibidi.com/img/cms/support/AN/AN33_Live_Dead_staining_with_FDA_and_PI.pdf*

*Other staining can be used to image viability and cell wall regeneration (see Notes and tips).*

For imaging, we need to gently immobilize cells without compressing them. Microfluidic channels provide an optimal solution that also ensures uniform cell dispersion, which significantly simplifies subsequent cell segmentation.

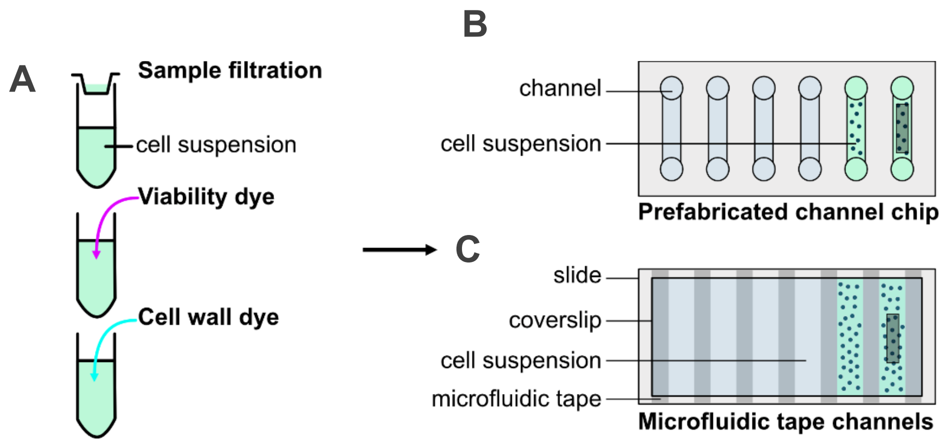


- Sample mounting:
  - Prefabricated channel chips (e.g.: IBIDI channel slides, B)

- Microfluidic tape channels: (C)
    - Microfluidic double-sided tape is used to bind slide and coverslip:

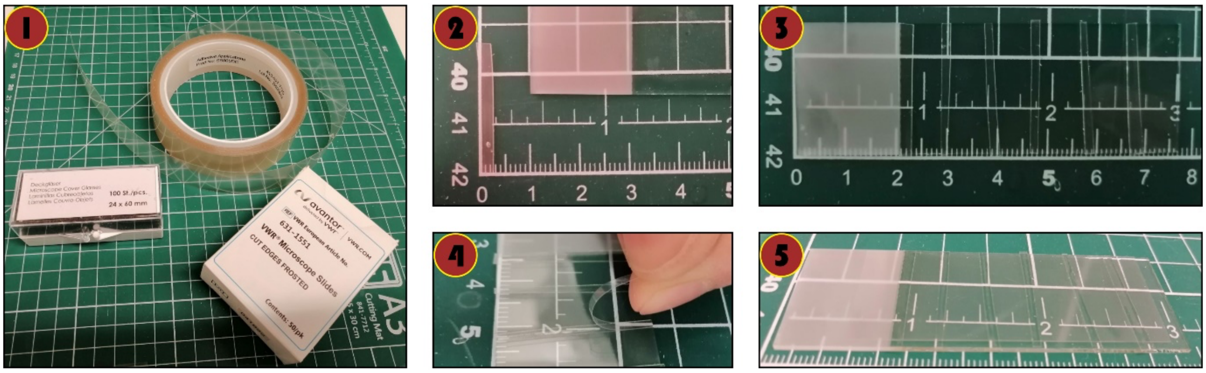


1- Material: double-sided microfluidic tape (e.g.: [Adhesive Applications S5005DC]), microscopy slide and coverslip.
2- Cut the microfluidic tape to have a length as long as the width of the microscopy slide and large of 2-3mm.
3- Remove one protective side and tape the slide to have about 5mm large channels (5 to 6 channels per slide with similar materials).
4- Remove the 2nd protective layer.
5- Put the coverslip on top of the tape and apply gentle pressure to make the tape glue to the coverslip.

- Cell suspension will be drawn into the channels through capillarity.

*Large images are preferable – lots of cells to observe per condition to have quantitative data.*

##### Notes and tips about imaging:

CW and viability staining can be done with other dyes. FDA and calcofluor have the advantage of being fast to stain. The viability could be also shown by the use of a fluorescent reporter line where the fluorescence is lost whenever the cells are dead. Cell wall can also be stained with Carbotrace (Ebba Biotech) or CarboTag (Besten et al. 2024).

### VII – Analysis

To find the best conditions for CW regeneration, analysis of the data is done using the Q-Warg pipeline. A complete guide is available as Method S2. In brief, the first step consists of segmenting the images using CellPose or another segmentation tool. The files are then organized to be used in an imageJ macro. This macro extract data from the images about fluorescence intensities of cell wall and viability staining and record morphometry measurements. The quantitative data is compiled and plotted. Associated with this, an app helps to check the viability threshold, segmentation and the validity of the results.

### Parameters to optimize

**Age of biological material:** For cell culture, it is crucial that cells are in the growing phase. It was observed that cells kept in the same medium for more than 5 days were detrimental for protoplast extraction. The resulting protoplasts and cultured protoplasts were in poor condition, with low survival rates. For other tissues, see Table S2.

**Enzymatic concentration and extraction time**: Enzymatic concentration can affect the health of protoplasts in culture. No obvious difference was observed when testing various extraction times. Note that the extraction time might lead to different degrees of CW digestion and therefore cell separation. It might be beneficial to limit the digestion time to have cells recover faster their CW as they would still have some CW component attached to their membrane. But that would mean that the cell suspension is not homogenous, and the CW would be different from one cell to another.

**Temperature**: The temperature during extraction is critical. While initial work was conducted at 22°C to maintain basal cell conditions, we observed that enzymes were more efficient at 24°C, resulting in higher protoplast concentrations. It was noted that temperature during culture might also require adjustment.

**Protoplast concentration**: Most literature protocols recommend working with 10^5 cells/mL. Observations showed that when protoplasts were dispersed in the medium, they tended to die faster and appeared unhealthy (exhibiting unusual shapes and low cell wall regeneration). No clear effect of higher protoplast concentration on cell wall regeneration was observed. However, it was noted that excessively high concentrations led to cell clumping, which could be problematic for applications requiring single cells.

**Composition of the CW regeneration medium**: The literature reveals nearly as many CW regeneration medium recipes as there are papers on protoplasting and plant regeneration from a single cell. This medium appears to be highly lab dependent. It was noted that most media used to enhance cell wall regeneration and promote callus formation are in gel form rather than liquid, which significantly reduces cell accessibility for applications requiring free-floating individual cells. Various parameters can be modified (Table S1).

### Bibliography:

Besten, Maarten, Milan Hendriksz, Lucile Michels, Bénédicte Charrier, Elwira Smakowska-Luzan, Dolf Weijers, Jan Willem Borst, and Joris Sprakel. 2024. "CarboTag: A Modular Approach for Live and Functional Imaging of Plant Cell Walls." bioRxiv. https://doi.org/10.1101/2024.07.05.597952.

Yoo, Sang-Dong, Young-Hee Cho, and Jen Sheen. 2007. "Arabidopsis Mesophyll Protoplasts: A Versatile Cell System for Transient Gene Expression Analysis." *Nature Protocols* 2 (7): 1565–72. https://doi.org/10.1038/nprot.2007.199.

Zhai, Zhiyang, Ha-il Jung, and Olena K. Vatamaniuk. 2009. "Isolation of Protoplasts from Tissues of 14-Day-Old Seedlings of Arabidopsis Thaliana." *Journal of Visualized Experiments : JoVE*, no. 30 (August), 1149. https://doi.org/10.3791/1149.
