## Supporting method S2 for "The Q-Warg Pipeline: A Robust and Versatile Workflow for Quantitative Analysis of Protoplast Culture Conditions"

User-guide for the analysis of cell wall regeneration post-protoplasting using the Q-Warg workflow

### Q-Warg

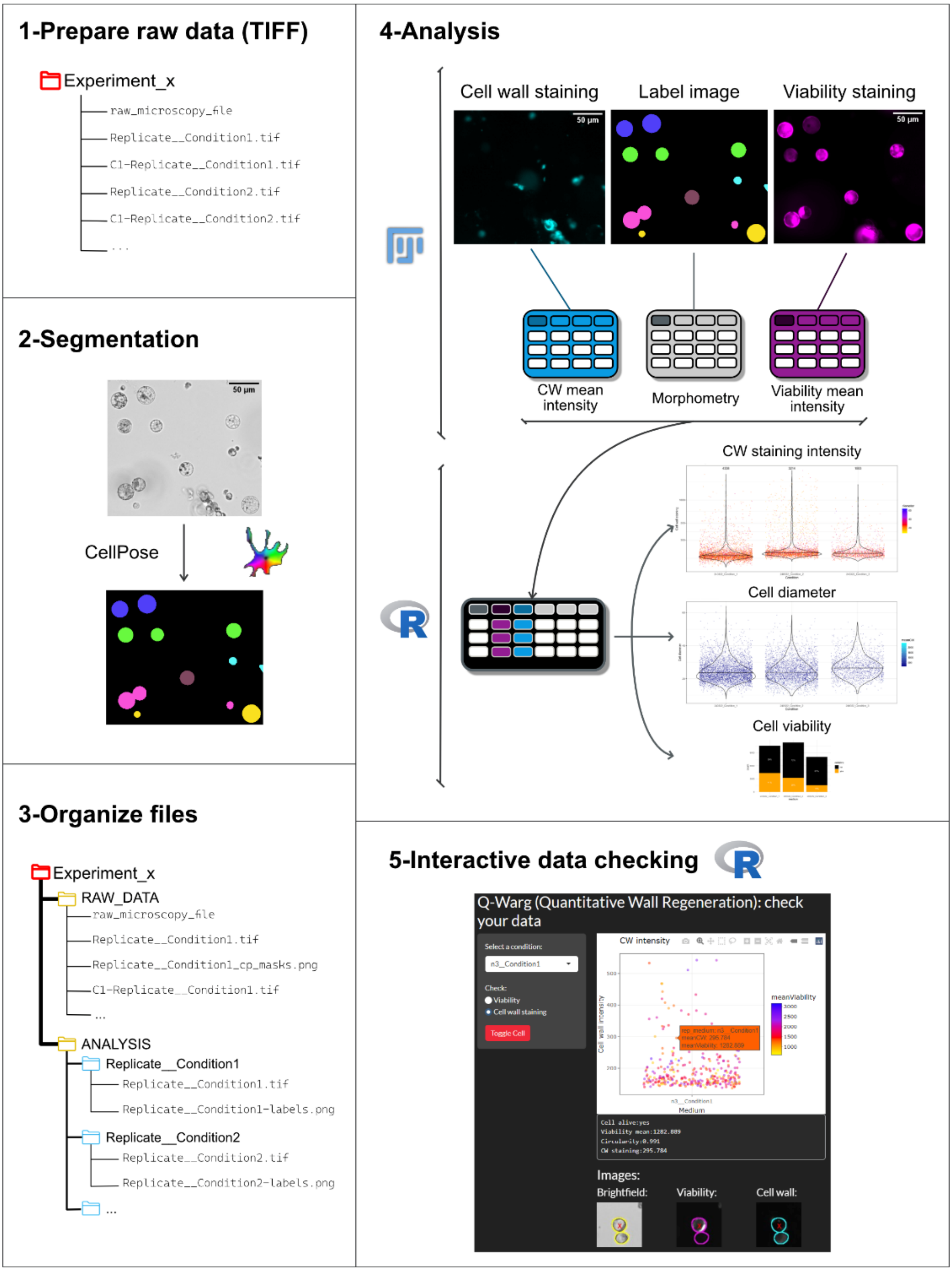

### Procedure overview and quick guide

Here we describe step-by-step how to operate the image analysis workflow to quantify viability and cell wall (CW) regeneration after protoplast extraction (see additional file 2) to help the user to screen quickly and efficiently CW regeneration media. This workflow uses different components under the form of ImageJ (Fiji) macro and R scripts (Method S3, S4). Tutorial videos can be found as on https://github.com/VergerLab/Q-Warg.git.

1- **Raw data preparation**: save_as_tiff.ijm
   a. Save raw images as TIFF with the 3 channels: brightfield, viability and cell wall staining
   b. Save a TIFF image with only the brightfield channel to be used for the segmentation
2- **Segmentation with CellPose**:
   a. Load the brightfield image into CellPose
   b. Run the segmentation
   c. Save the label image as PNG
3- **Organizing files for Q-Warg**: R_orga_folder.R
   a. Prepare a raw data image that will stay untouched by the pipeline
   b. Prepare the folders to be used with the pipeline
      i. Each folder contains the 3-channels image and the corresponding label image
      ii. Rename files to fit Q-Warg requirements
4- **Analysis**:
   a. Run the ImageJ macro CWRegenerationQuantification.ijm to extract quantification data: viability and cell wall staining intensity, morphometry information (such as size and circularity)
   b. Run the R script QWARG-notebook.Rmd:
      i. Tidy data
      ii. Set viability threshold
      iii. Plot data
      iv. Save data table and plots of cell wall mean intensity and cell size
5- **Interactive data checking**:
      v. Reorganize the folders
      vi. Check data: observe the cells corresponding to the quantitative data

See below for the detailed guide.

### Pre-requisites:

#### Software & plugins required:

- Anaconda
- CellPose
- Fiji (https://fiji.sc/)
  - MorphoLibJ (“IJBP-Plugins” update site)
- RStudio (https://posit.co/download/rstudio-desktop/)
  - Libraries:
    - data.table (Barrett *et al.*, 2024)
    - rlist (Ren, 2021)
    - hrbrthemes (Rudis [aut *et al.*, 2024)
    - viridis (Garnier *et al.*, 2024)
    - tidyverse (Wickham *et al.*, 2019)
    - tcltk (Grosjean, 2014)
    - shiny (Chang *et al.*, 2024)
    - plotly (Create Interactive Web Graphics via ‘plotly.js’ [R package plotly version 4.10.4], 2024)
    - magick (Ooms [aut and cre, 2024)
    - autothreshold (Nolan [aut *et al.*, 2023)
    - patchwork (Pedersen, 2024)
    - agricolae (Mendiburu, 2023)
    - bslib (Sievert *et al.*, 2025)
    - sortable (Vries *et al.*, 2023)

#### Workflow scripts:

- save_as_tiff.ijm
- R_orga_folder.R
- CWRegenerationQuantification.ijm (Method S3)
- QWARG-notebook.Rmd (Method S4)

#### Data:

- The original image in TIFF with the 3 channels:
  - Brightfield (should always be channel 1)
  - Viability staining
  - Cell wall staining

#### Installation

##### 1- GitHub

Both CellPose and Q-Warg are available on GitHub.

- Go on the repository page with the provided link.
- Click on Code
- Choose Local

- Download ZIP: the repository will be downloaded into the download folder of your computer.
- Extract all files from the compressed folder to your folder of choice

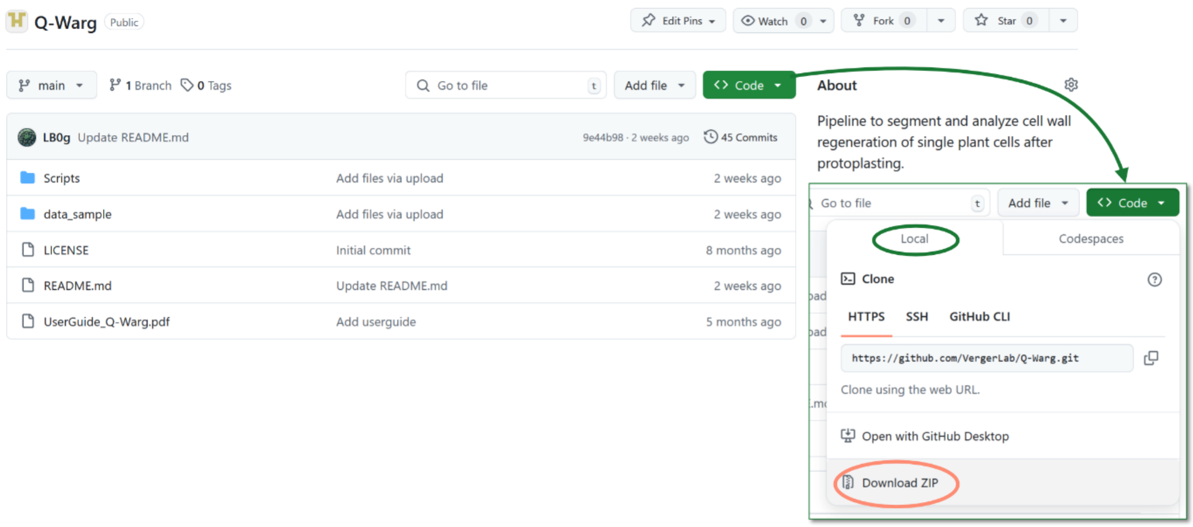

#### 2- CellPose

We used CellPose for segmentation, but alternative segmentation tools could be used as well, as long as the output is (or is made) compatible with the rest of the workflow.

##### Anaconda

CellPose requires a python environment that can be provided by Anaconda.

Download and install anaconda here: https://www.anaconda.com/download/success.

##### CellPose

Detailed guidelines to install CellPose can be found on the readme file of the CellPose repository in GitHub: https://github.com/mouseland/cellpose

#### 3- Download Q-WARG from GitHub

All scripts can be found in the Scripts folder of the GitHub Q-Warg repository found here: https://github.com/VergerLab/Q-Warg.git

#### 4- To use ijm files

##### Fiji and MorpholibJ

To install Fiji/ImageJ, see https://fiji.sc/

To install the plugin MorphoLibJ, turn on the IJBP-Plugins update site:

1. Go in the “Help” menu of Fiji and click on “Update…”. This will open the “ImageJ updater”.
2. In “ImageJ updater”, click on “Manage update sites”.
3. In the list, find e.g. “IJPB-Plugins” and click in the square next to it to add it.

4. Then close the window and apply changes on "ImageJ updater".
5. Once this is done, Fiji needs to be restarted.

*If more explanation needed, see* https://imagej.net/update-sites/following

##### Macro

Both imageJ macro CWRegenerationQuantification.ijm and save_as_tiff.ijm are in the downloaded folder **Scripts**.

- Drag & drop the file on Fiji user interface to open a macro
- Start the macro by clicking Run in the macro window

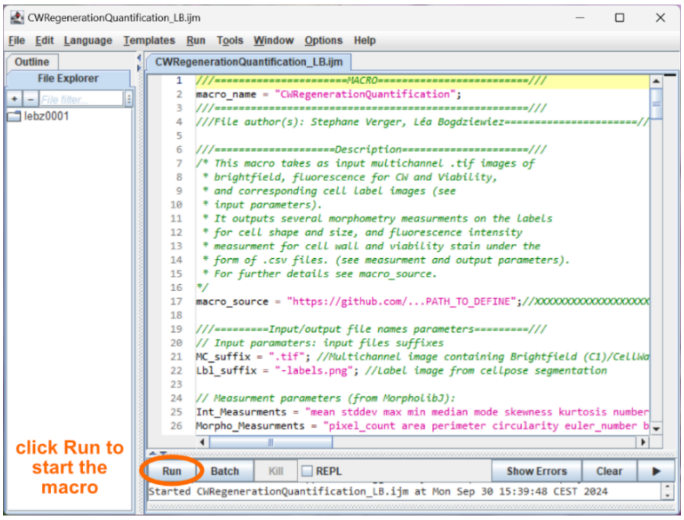


##### 5- RStudio

- Install RStudio see: https://posit.co/download/rstudio-desktop/
- Install the packages needed to run the different scripts. Copy/paste the following code in the console of RStudio:

```
# Define the packages you want to install
packages <- c("data.table", "rlist", "hrbrthemes", "viridis", "tidyverse", "shiny",
"tcltk",  "plotly", "magick", "autothreshold", "bslib", "sortable", "agricolae",
"patchwork")
# Find out which packages are already installed
installed_packages <- rownames(installed.packages())
# Determine which packages are not installed
packages_to_install <- setdiff(packages, installed_packages)
# Install the packages that are not yet installed
if (length(packages_to_install) > 0) {
  install.packages(packages_to_install)
} else {
  cat("All packages are already installed.\n")
}
```

The code is also available in the downloaded Q-Warg-Scripts folder as Install_R_packages.R.

*The code checks if the required packages are installed and adds the ones that are not. For more information on how to install packages, see* https://cran.r-project.org/doc/manuals/r-release/R-admin.html#Default-packages *.*

Once RStudio and the scripts are downloaded, double-click on the script QWARG-notebook.Rmd to open it with RStudio. It is then ready to use.

### Q-Warg: Using the pipeline

The Q-Warg pipeline can be fully tested using a sample dataset available in the GitHub repository. This dataset, located in the data_sample folder, includes the original TIFF images for 3 replicates under 2 experimental conditions and their corresponding CellPose labels (in data_sample_labels.zip).

Using these files and the provided scripts, you can run the complete Q-Warg workflow to verify its functionality before applying it to your own data. The user guide includes screenshots that illustrate the expected output at each step of the process.

#### 1- Prepare raw data

##### Files naming:

- **All files' names should have the following format:** Replicate__conditionX.tif
- **Replicate can be the date or nx. It is needed for the plots later.**
- **Use "__" to separate Replicate from condition. It should only be used to separate replicate from the used condition.**
- **Do not use "--" (simple – is fine) to not interfere with the scripts.**
- **Do not use spaces or dots (except for file extension) in the path of files names**

*If you plan on making only one replicate for screening, put the date as first part of the file name (e.g.: 240213__medium2).*

*Do not change the names of the files after starting using the pipeline. It is a crucial parameter onto which several operations are based.*

##### Save 3-channels images as TIFF:

The aim of this step is to convert proprietary format files issued from microscopes (like .lif for Leica or .czi for Zeiss) into tif files. TIFF files will then be used in to run the pipeline.

**Option 1:** (automated)

- Open images in Fiji by dragging and dropping the file onto Fiji user interface
- Open the macro save_as_tiff.ijm by dragging and dropping the file onto Fiji user interface

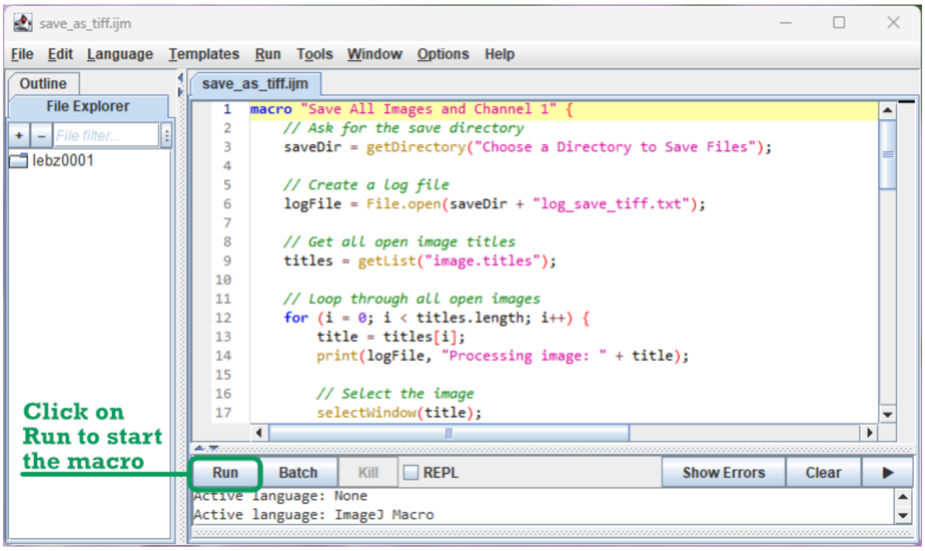


- Run the macro by clicking Run:
    - Choose a folder where all images will be saved as tiff: 3-channels images and the C1 brightfield channel alone to be used in CellPose for segmentation.
- A log file is saved to check if every image has been saved as expected.

**Option 2:** (manual)

- Open images:
    - Open images in Fiji by dragging and dropping the file onto Fiji user interface
    - To save: File > Save As > Tiff…
- Save the brightfield (BF) image independently to be used for the segmentation step:
    - Image > Color > Split Channels
    - Save as TIFF the Brightfield channel
- Close images

*We advise keeping a raw data folder containing all raw images and the saved TIFF.*

The folder should have an organization like:

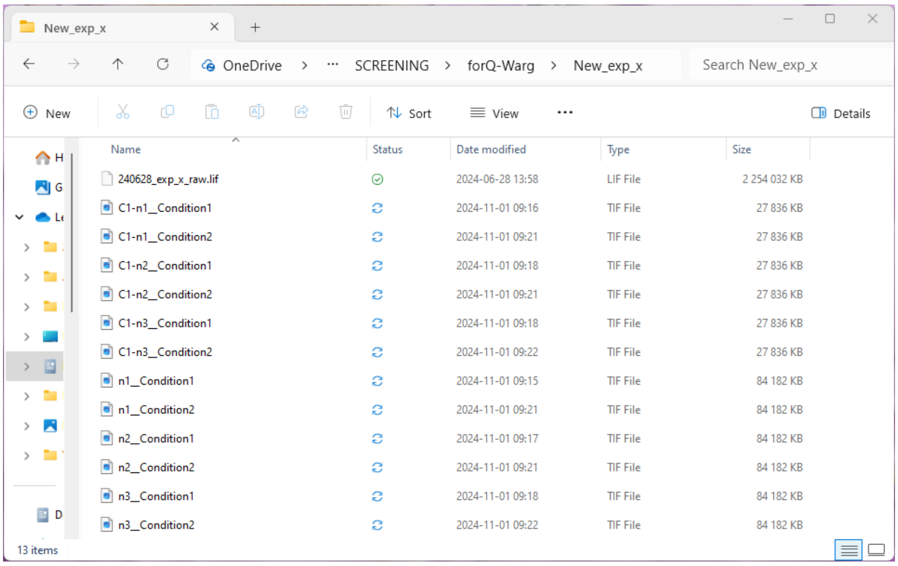
New_exp_x
OneDrive
SCREENING
forQ-Warg
New_exp_x
Search New_exp_x
New
Sort
View
Details
Name
Status
Date modified
Type
Size
240628_exp_x_raw.lif 2024-06-28 13:58 LIF File 2 254 032 KB
C1-n1__Condition1 2024-11-01 09:16 TIF File 27 836 KB
C1-n1__Condition2 2024-11-01 09:21 TIF File 27 836 KB
C1-n2__Condition1 2024-11-01 09:18 TIF File 27 836 KB
C1-n2__Condition2 2024-11-01 09:21 TIF File 27 836 KB
C1-n3__Condition1 2024-11-01 09:18 TIF File 27 836 KB
C1-n3__Condition2 2024-11-01 09:22 TIF File 27 836 KB
n1__Condition1 2024-11-01 09:15 TIF File 84 182 KB
n1__Condition2 2024-11-01 09:21 TIF File 84 182 KB
n2__Condition1 2024-11-01 09:17 TIF File 84 182 KB
n2__Condition2 2024-11-01 09:21 TIF File 84 182 KB
n3__Condition1 2024-11-01 09:18 TIF File 84 182 KB
n3__Condition2 2024-11-01 09:22 TIF File 84 182 KB
13 items

#### 2- Cell segmentation: CellPose

Cell segmentation is required for extracting data from the images. CellPose, more specifically cyto3 generalist model, is working very well for our samples. This model should work for most of the isolated cells issued from protoplasting. However, any segmentation program can be used as long as it gives a PNG label image to be used in the next steps of the pipeline.

*Detailed information about the software and how to use it can be found here: [http://www.cellpose.org/](http://www.cellpose.org/) ; [https://cellpose.readthedocs.io/en/latest/gui.html](https://cellpose.readthedocs.io/en/latest/gui.html)*

- Drag & drop a brightfield file in the user interface to open the image:

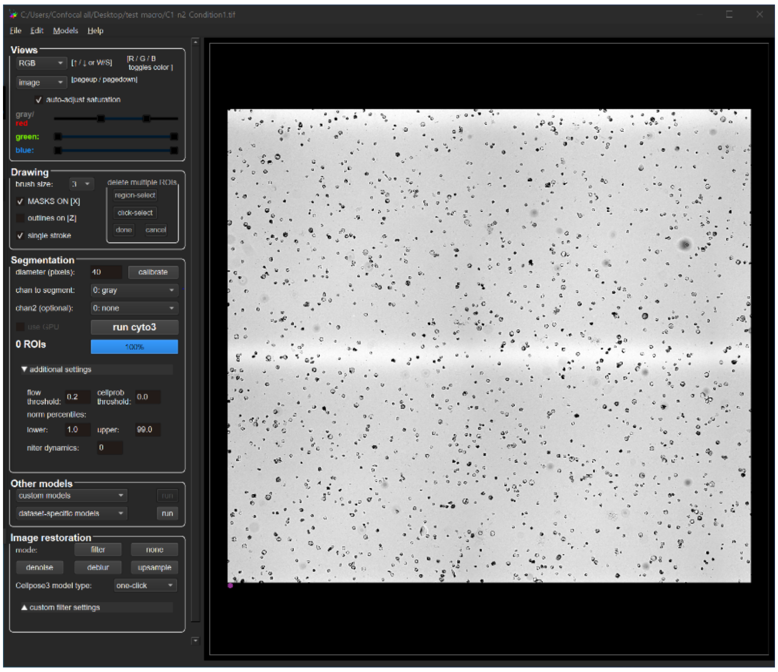


- Adjust parameters to fit your cells

*Used in the publication: Parameters used for CellPose with cyto3 defaults settings:*

  - *Cell diameter (pixels): 40*
  - *flow_threshold: 0.2. "The `flow threshold` parameter is the maximum allowed error of the flows for each mask. The default is `flow_threshold=0.4`. Increase this threshold if CellPose is not returning as many ROIs as you would expect. Similarly, decrease this threshold if CellPose is returning too many ill-shaped ROIs."*

- Run segmentation
- Save labels as PNG in File > Save masks as PNG/tif (the PNG image is automatically saved in the same folder as the original image).

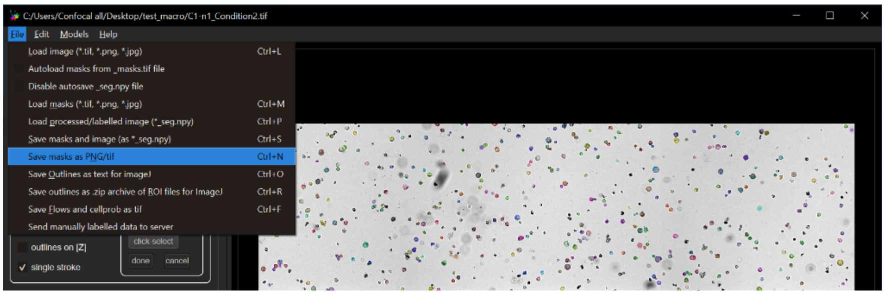


*We recommend saving a copy of the label image in the raw data folder.*

#### 3- Data organization

The data with which to use Q-Warg should be organized as follows:

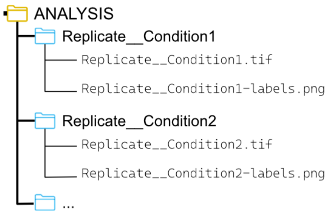


Each condition has its own folder containing the 3-channels image as TIFF and the label image as PNG. After segmentation with CellPose, the folder resembles:

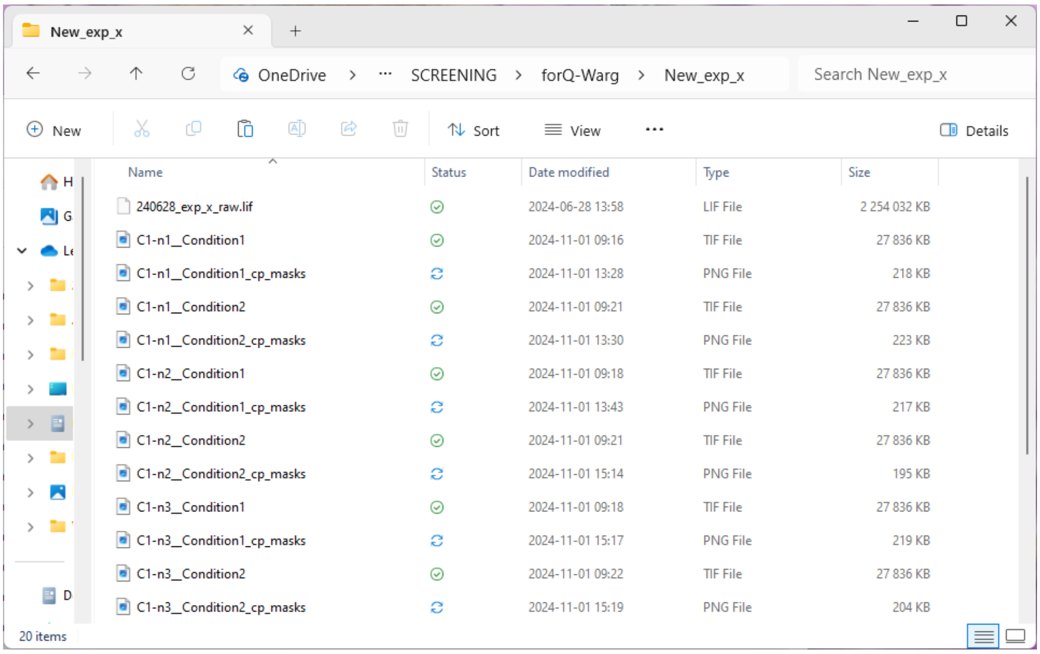


##### Organize folders:

**Option 1:** (automated)

- Open the R script R_orga_folder.R by double clicking on it (RStudio will automatically start).
- Run the code:

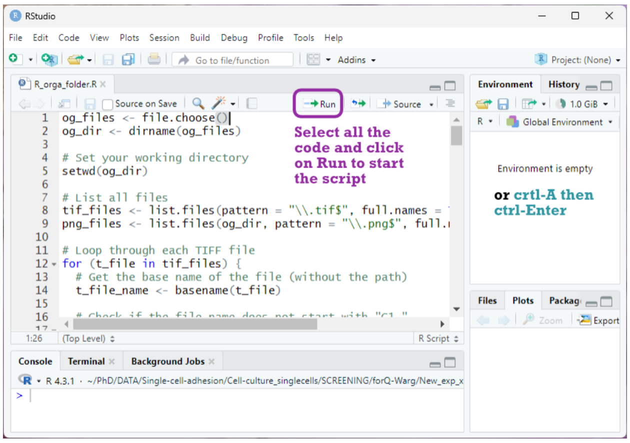

- A pop-up window will open to choose which folder to organize. Select any file in this folder to get it tidy.

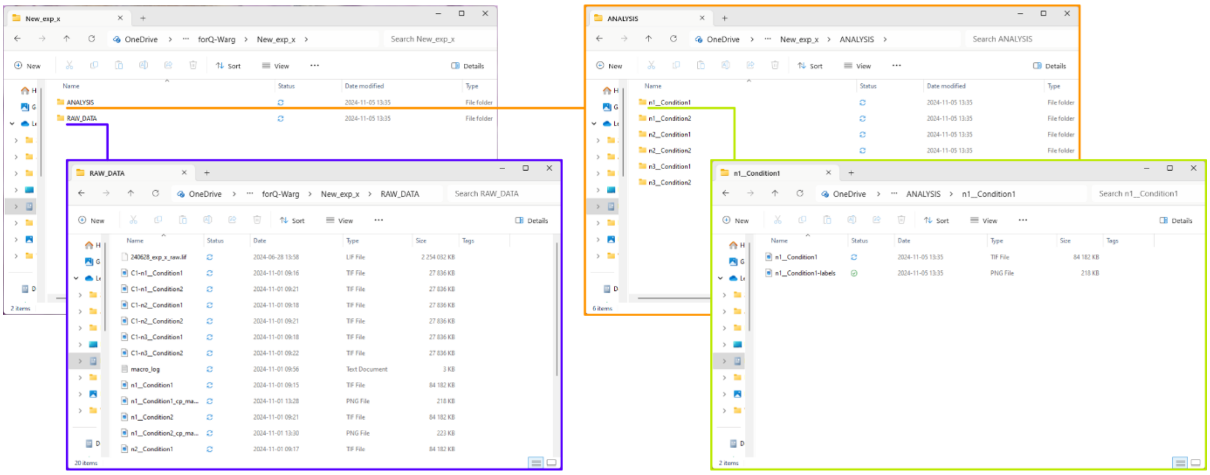


*The script can also be used if the segmentation method is not CellPose if it is a PNG file and it contains the name of the original file. The script removes any extension to the file name and adds -labels.*

**Option 2:** (manual)

- Please organize the folders as described above, creating one folder for each condition. Inside each folder, include only the 3-channel TIFF image and the corresponding label image.
- Naming: (see section 1 for other guidelines)
  - Both images (3-channels TIFF and the PNG label image) must have the same base name e.g.: “Condition_01” (make sure to give meaningful names).
  - The label image must contain “-labels” at the end of the name
  - Do not use spaces

The working folder for the next steps is here called “ANALYSIS”

*By using the automated script, the RAW_DATA folder contains already the original images and the label images. Tip: the RAW_DATA folder can be compressed to free up space.*

#### 4- CWRegenerationQuantification.ijm (Fiji macro)

The data are now organized and ready to be used for analysis. The next step is to start the quantification for fluorescence intensity and cell morphometry.

- Start the macro (as detailed in the installation part of this guide).
- A pop-up window opens and asks to choose in which order are the channels: for each Cn-, select the corresponding channel (BF: Brightfield; CW: Cell Wall; Via: Viability).

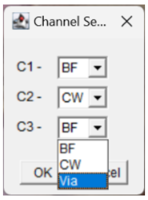

- Choose the working folder in the pop-up window. *In the example here "New_exp_x\ANALYSIS.*

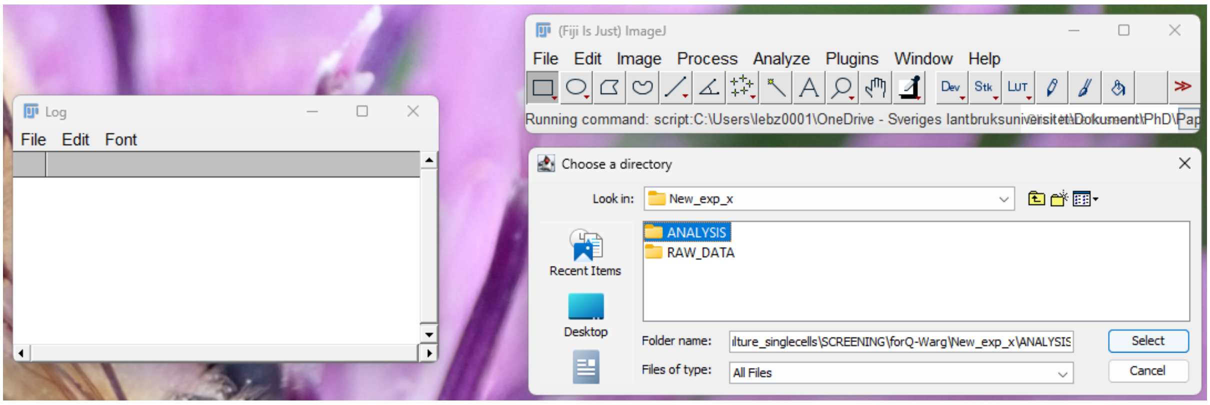


- Let the macro run:
    - Log window opens and allows you to follow the different steps of the process.

  *If you want to run the macro on only one condition, you need to put your sample folder into a working folder that only contains this folder of interest. The macro will report an error if it is running directly in the sample folder.*

  *When running the macro, try not to use the computer, especially the keyboard, because pop-up windows will come on top of all the other windows open and you might terminate the process by mistake.*

    - Check the log window to know when the processing is done

  *The log is also saved into the working folder. It can be checked to verify all processed conditions.*

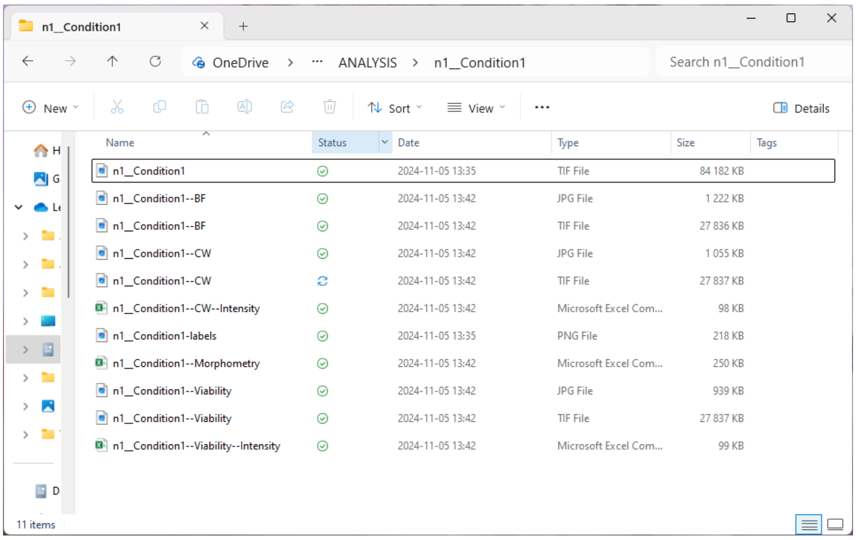


The sample folders now contain many new files:

- The macro splits the 3-channel TIFF image and saves each channel independently (for later potential use).
- Each channel in JPG with the segmentation border to be used to check the images in the Q-Warg shiny app.
- CSV files with morphometry (size, circularity, position), fluorescence intensity for viability and for cell wall.

#### 5- QWARG-notebook.Rmd

##### Analysis and plots

With the R script, the csv files obtained from the macro are combined and reorganized to obtain information about the living cells that have been segmented. Detailed information about the script can be found as comments directly in the script file QWARG-notebook.Rmd. The notebook is composed of several chunks (part of code) and each one is briefly described.

- Open QWARG-notebook.Rmd by double-clicking on it (RStudio will automatically start).
- Run the code: each chunk (a chunk is a block of code that will be executed as a unit, R name for a code block) can be run by clicking on the play sign at the top of it.

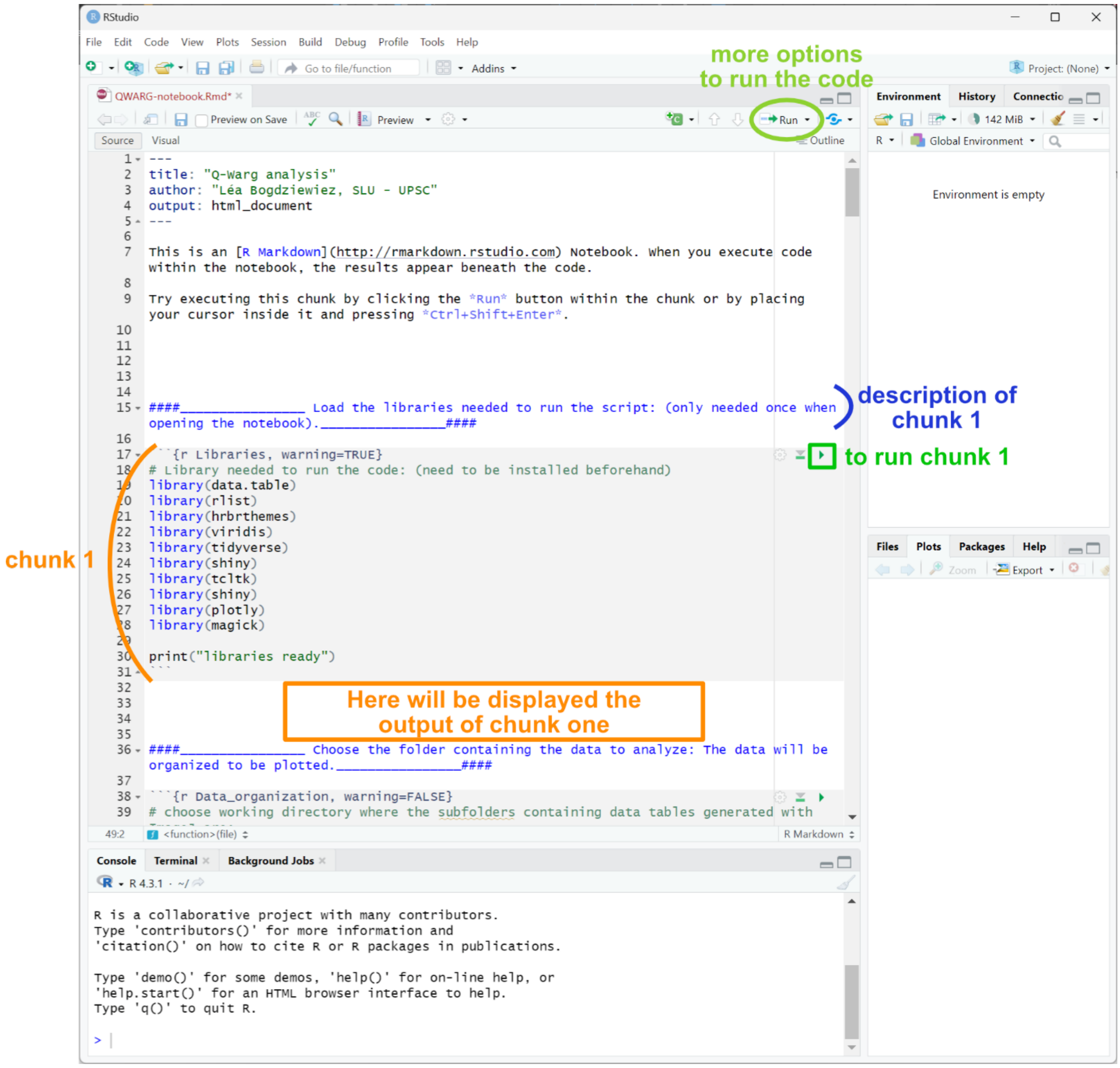

- Chunk 1: load the libraries, always needed to run other parts of the code.

  *If you only need to run the shiny app, or other chunks of the app, you first need to run the first chunk to load the libraries.*

- Chunk 2: data frame
  - Define the working folder (here ANALYSIS) in the pop-up window: choose the Log file created when using the CWR macro.

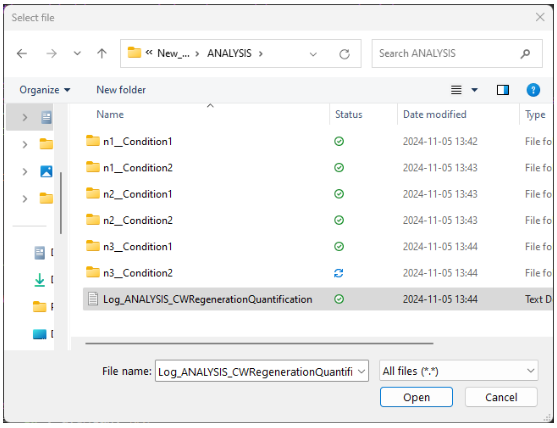


*Possible issue at this step: the window might open in the background, check the taskbar.*

  - Viability threshold is calculated for each condition with both the Huang and the triangle automatic threshold methods.
  - An organized data table is generated (see additional information section for more information).

- Chunk 3: viability threshold
  - An application opens to set the viability threshold for each condition:
  - The viability staining intensity is represented in 2 ways to help estimate which threshold is the most adapted to the data.

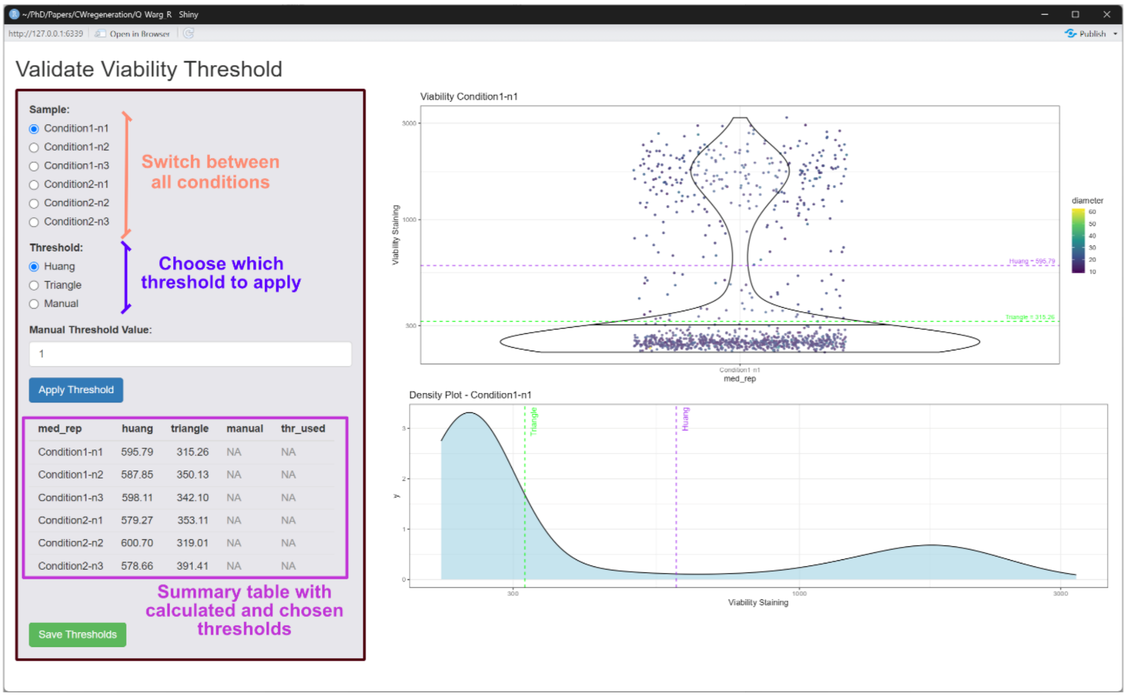


- Choose which threshold to apply for each condition. Once it has been selected (under Threshold) and if relevant, the manual value written, click on Apply Threshold to save the value in the table. A red line will be added on the plots to show the selected value.
- Click on Save Thresholds to save the table once all values have been added to the last column.

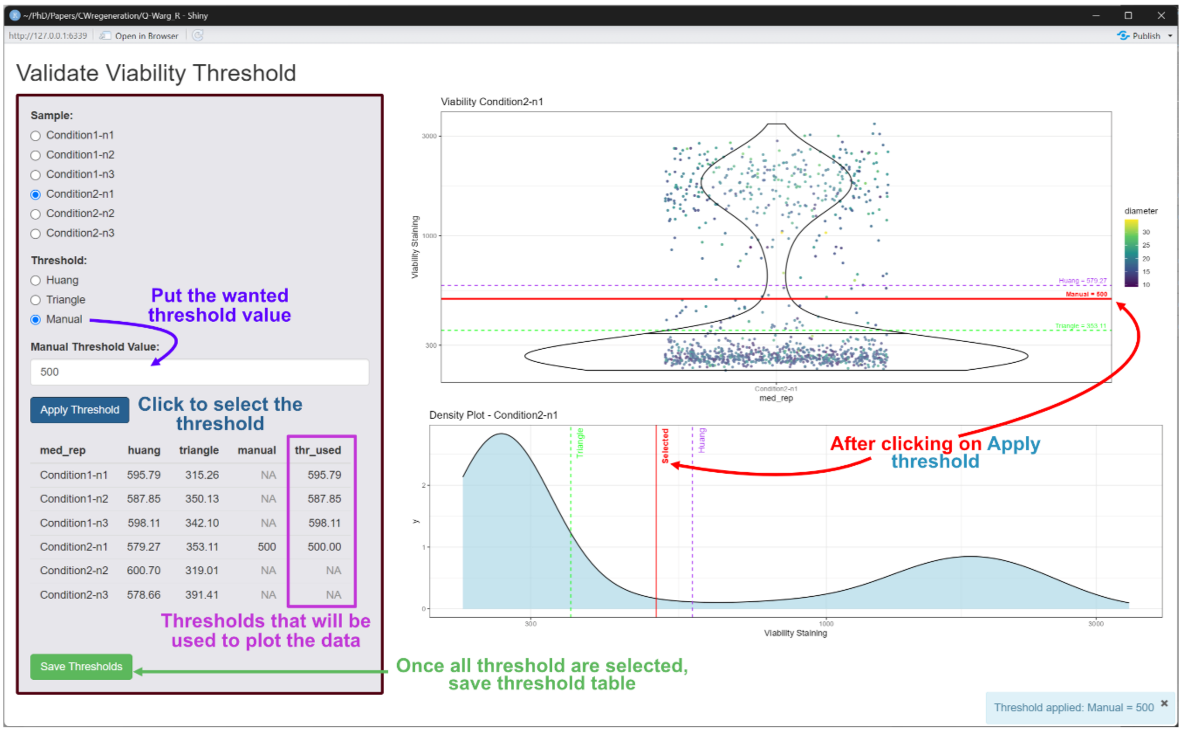


*The threshold table will be saved later in the workflow, with a mention of which type of threshold has been used for each condition.*

To determine if the segmented objects are living cells, the script uses viability staining and the circularity of the cells. A threshold is applied for cell viability staining. Below this value, the cells are considered dead. This threshold is dependent on the viability staining and the imaging setup so it should be adjusted if needed.

2 automatic thresholding methods are available: Huang or Triangle. Only one method should be used for one data set to obtain consistent results.

As automatic thresholding methods do not consider the values but only the shape of the data, it might lead to thresholding even if there is only a dead population. The app counteracts this with the possibility of adding a manual threshold. The manual threshold should be chosen carefully. For instance, it could be the minimal value of the chosen automated threshold on other conditions.

In addition, to ensure that the object is a cell, a high threshold is used for circularity (0,9). Indeed, after protoplasting, the cells lose their original shape and become spherical. In absence of directional force on the cell, they stay spherical in the early phase of regenerating their cell wall. If experiments are conducted with different cell shapes, it might be needed to adjust this parameter. All the code corresponding to the thresholding can be found in the last part of the chunk 2 (indicated as comment in the code).

- Chunk 4: plot cell wall intensity, cell size and cell viability generation
    - The CW staining mean intensity is represented in 2 plots:
        - With dots color corresponding to the replicate
        - with dots color corresponding to cell diameter
    - The CW staining median intensity
    - The viability staining intensity is represented the same way as the CW staining intensity
    - Live and dead objects count and % of each in the population of segmented objects.

  *The black percentage cannot be interpreted as a number of dead cells but non-living objects. While dying, a cell might have burst into several parts that have been segmented.*

    - Plots with automatic viability thresholds: one dot plot and one density curve per condition.

  *All plots corresponding to the data sample can be found on GitHub in the data_sample folder.*

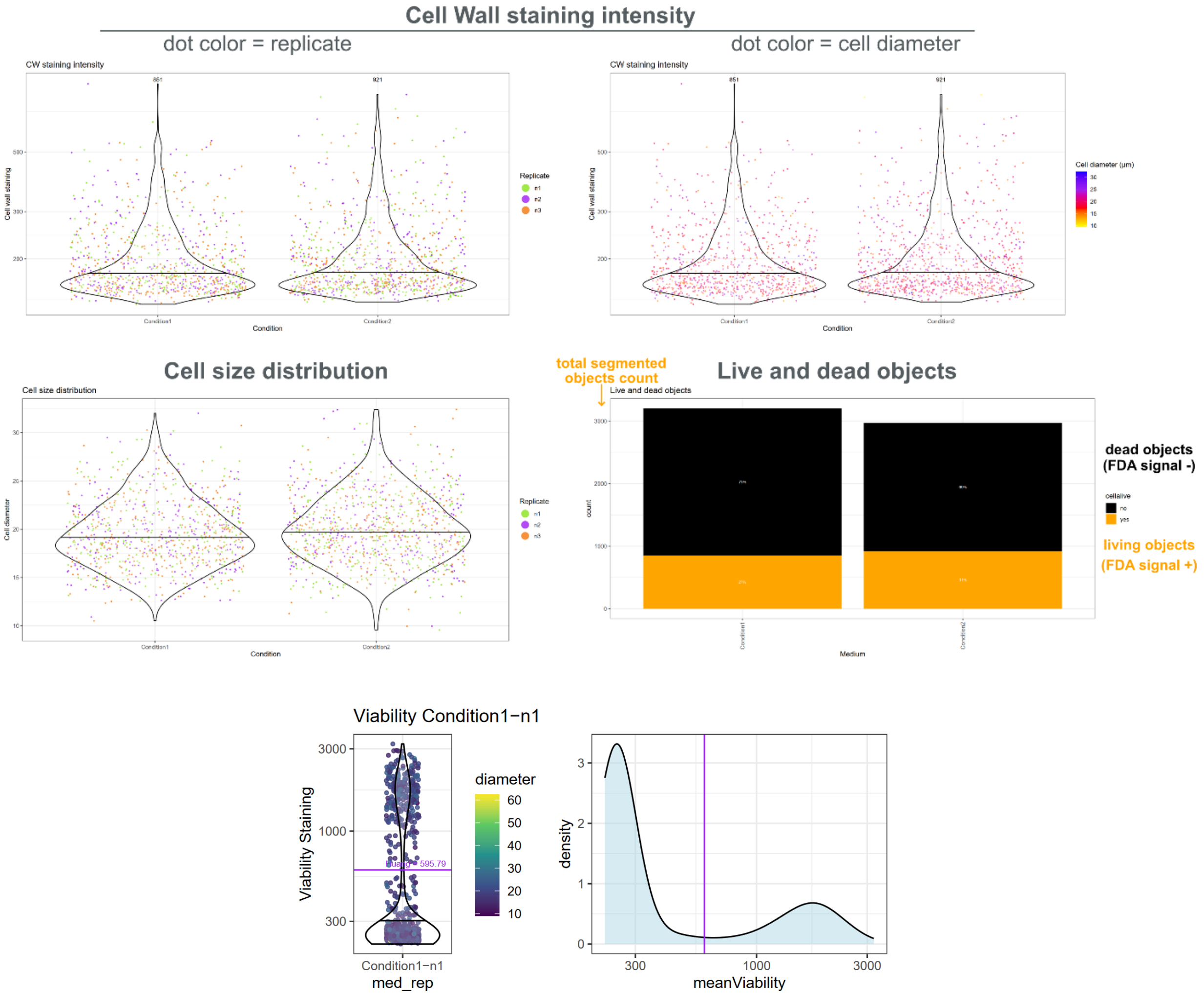


- Statistical analysis is also made in the chunk 4 prior to the plots: We perform statistical analysis withing the R notebook using the xxx library… One-way Analysis of Variance (ANOVA) followed by Tukey's Honestly Significant Difference (HSD) test were performed for multiple comparisons of means for CW staining intensity and cell size plots. Chi-square test of independence (significant if p < 0,05) was performed, followed by pairwise comparisons using the pairwise.prop.test function with Bonferroni correction for multiple testing.

- Chunk 5: save plots and data frame
  - Choose a file in the folder where you want to save the plots and data frame, *e.g.*: Log_Analysis to save in the Analysis folder (same way as for chunk 2)

6 files are saved:

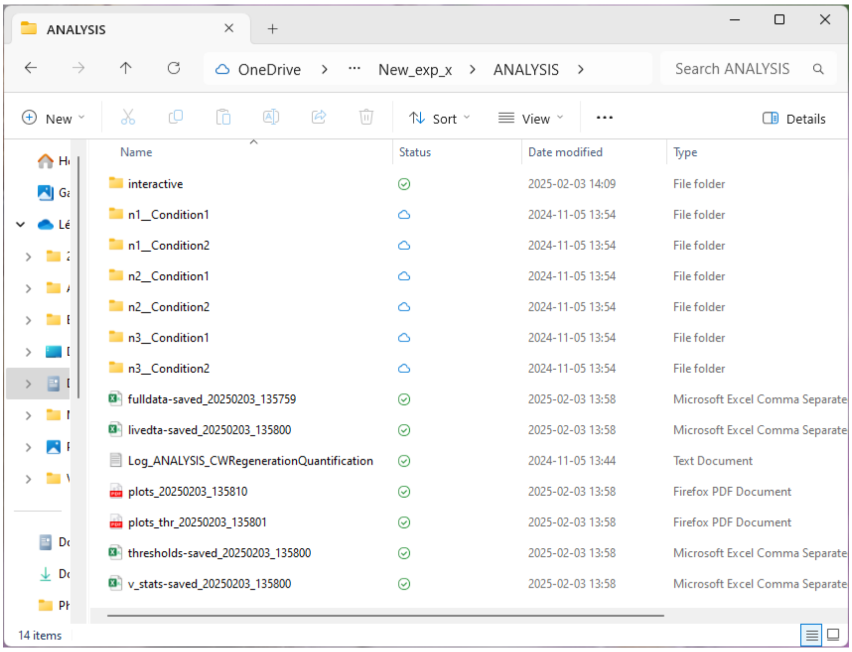


- Plots described in the chunk 3 section:
  - Data plots
  - Viability threshold plots
- threshold_saved.csv: contains the selected threshold values.
- viability_stats.csv: results of the statistical analysis to compare viability of the cells between each condition (chi-square followed by Bonferroni). If the result < 0,05, the difference is significant. This test is to be associated with the viability bar plot.
- full_data_saved.csv: contains all data, for live and dead objects.
- livedta_saved.csv: data for living cells only. It should look like this:

| | cell | medium | diameter | Circularity | meanCW | meanViability | cooX | cooY | cellalive |
|---|---|---|---|---|---|---|---|---|---|
| 1 | 240322_Condition_1-100 | 240322_Condition_1 | 36.69158 | 0.986 | 155.192 | 681.747 | 675.339 | 111.483 | yes |
| 2 | 240322_Condition_1-1000 | 240322_Condition_1 | 27.90336 | 1.001 | 178.394 | 1428.671 | 2164.137 | 1040.369 | yes |
| 3 | 240322_Condition_1-10001 | 240322_Condition_1 | 17.82535 | 0.986 | 263.421 | 1538.608 | 2771.289 | 10304.394 | yes |
| 4 | 240322_Condition_1-10014 | 240322_Condition_1 | 20.87222 | 0.986 | 194.312 | 860.608 | 2891.682 | 10311.533 | yes |

*The names of the files contain date and time. The plots and data table can be made and saved several times without replacing the older ones.*

*A more detailed explanation of the data tables can be found in the additional information section at the end of this user guide.*

#### Q-Warg shiny app

The aim of this app is to check that the analysis pipeline worked as expected: each point of the plot corresponds to a living cell and the viability threshold is correct. It allows you to load the full data file obtained with the previous script and click on each point of an interactive graph to see the corresponding cell in the 3 channels of the original picture.

To access both the images and the data table, the data must be reorganized. All the JPG images and the livedta-saved should be in one folder here called "interactive".

##### Reorganize data in the working folder:

**Option 1:** (automated)

- Run Chunk 6 in the QWARG-notebook.Rmd
- Choose the same working directory as before: choose any file in the Analysis folder. E.g. fulldata_saved.csv

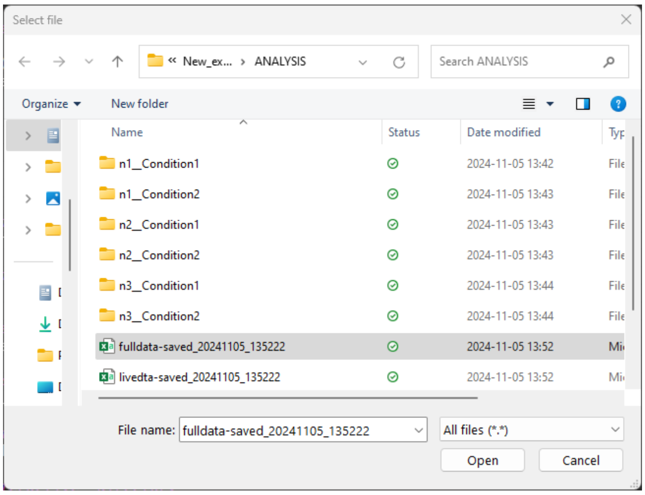


- The interactive folder is created, JPG files are moved inside, and the fulldata_saved csv file is copied.

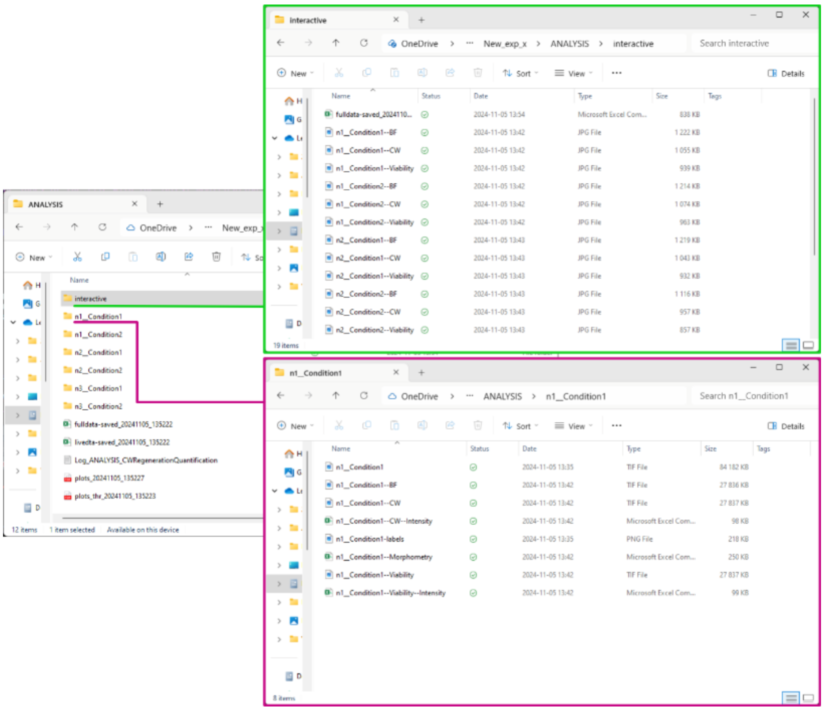

**Option 2:** (manual)

- Create a new folder called interactive.
- Move by hand all JPG images from the different sample folders and the livedta-saved csv file into this folder.

*The name of the folder is not important, if using Option 2 it will be named "**interactive**" by the script. The crucial point is to keep the naming of all files, otherwise the app will not be able to connect the images with the quantifications.*

- Open the application by running Chunk 7
  - Choose any file in the interactive directory in the pop-up window

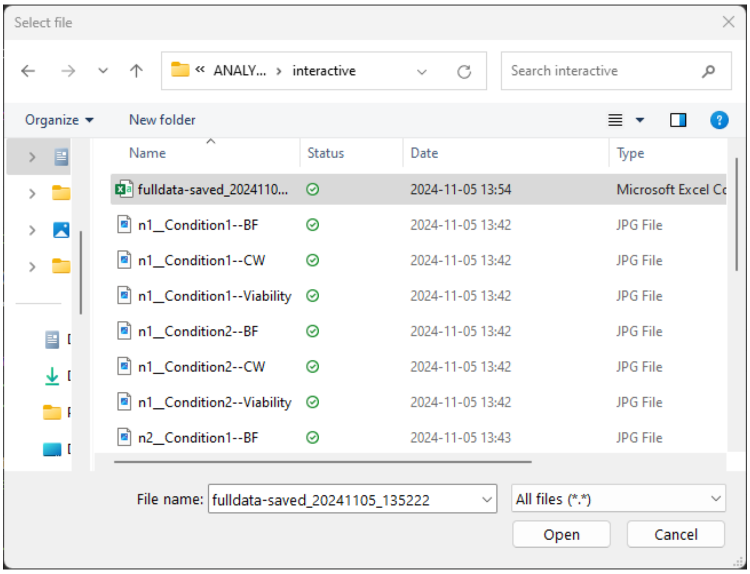


*An issue that can arise at this step: the window might open in background, check the taskbar.*

- The app window is opening.

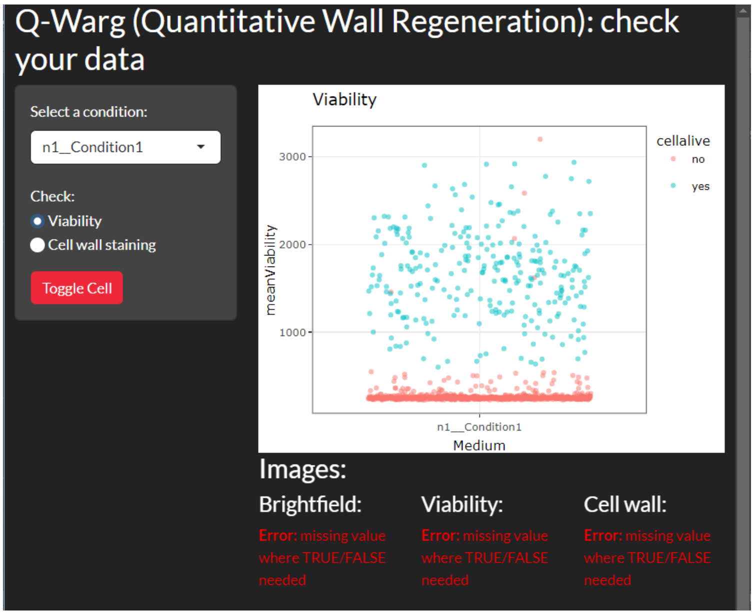


On the left panel, a rolling menu allows you to choose which condition to check. Only one condition at a time can be visible.

Two different plots can be displayed:

- Viability plot with the viability staining intensity in Y axis. All segmented objects are plotted, the color of the dot shows if the point corresponds to a dead or a living object.
- Cell wall staining intensity (Y axis), the color of the dot corresponds to the viability mean intensity. Only living objects are displayed.

The plots are interactive, hovering the mouse over a data point will display information about it:

- The condition it comes from
- CW mean intensity
- Viability mean intensity

By clicking on one data point, other information will be displayed below the plot:

- If the cell is alive yes/no
- Viability mean intensity
- Circularity: to check if the object is considered dead because of its viability staining or the circularity threshold (default=0,9).
- CW staining mean intensity

Below, the 3 channels of the segmented image are displayed with the object corresponding to the clicked point in the middle of the image. We can then check if the segmentation is well made thanks to the contour line that is represented (yellow for BF image, magenta for viability, cyan for CW).

*Before clicking on a data point, the error message (Error: missing value where TRUE or FALSE needed) should be written (see previous image). It just means that a point must be selected to show an image. If another error message is displayed, it probably means there is an issue in the naming of the JPG images in the interactive folder.*

A red button "Toggle cell" makes a cross appear on the images to check which cell is the one designated by the point in the plot.

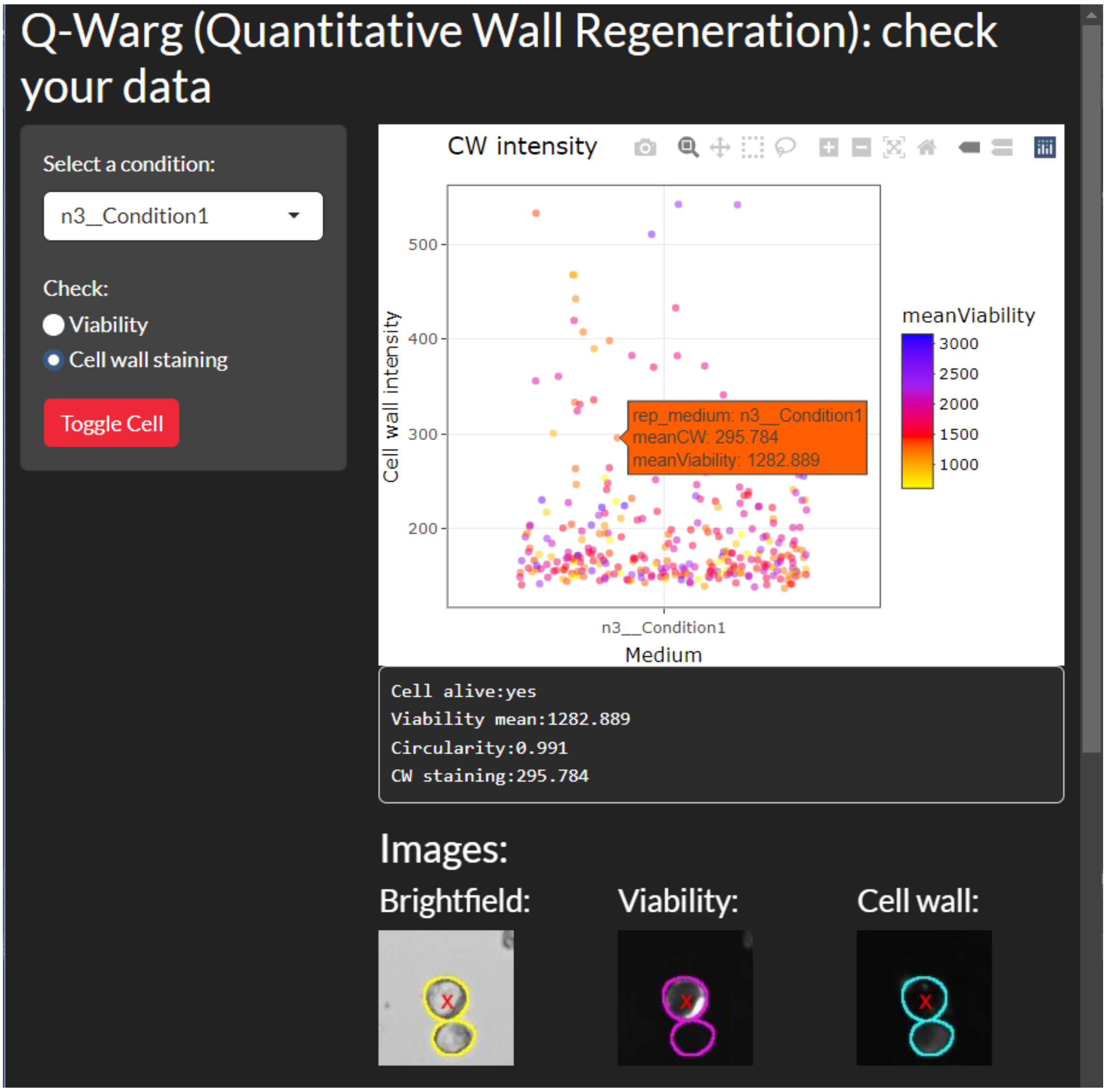


Below the images, text is displayed: "Clicked Point Coordinates" corresponds to the coordinates of the point in the plot. It is based on the Y value that the script will find back the right coordinates in the jpg image corresponding to the chosen condition. "Picture coordinates" correspond to the cooX and cooY columns of the data table.

### Troubleshooting:

#### Issues with the code:

Most errors that could arise are linked to the naming of the files. To reduce this risk, use all automated options of this pipeline and the guidelines for original image naming:

Replicate__condition.tif

No use of **other** double _ nor double -, do not use dots and spaces in the names. It goes also for folder names: if there is a space in the path of the experiment data folder, it might create trouble with the code.

### R:

Several parameters can be modified in the code. The parts of code that can be easily changed are highlighted with comments in the code with the following structure:

```
#______________________________________________________________________________#
##____________________________________________________________________________##
### THRESHOLDS FOR CELL VIABILITY: BASED ON VIABILITY STAINING AND CIRCULARITY ###
##____________________________________________________________________________##
#______________________________________________________________________________#
```

● Check the installation of the libraries

● Interactive Q-warg shiny app: the coordinates of the images need to be converted into pixels for the program to find the cell corresponding to the clicked data point. The value might be different depending on the imaging system. This can be modified in the first part of the code corresponding to the Q-Warg app: chunk 6. First, we need the converting factor from micrometers to pixels:

In ImageJ go to Image > Properties…:

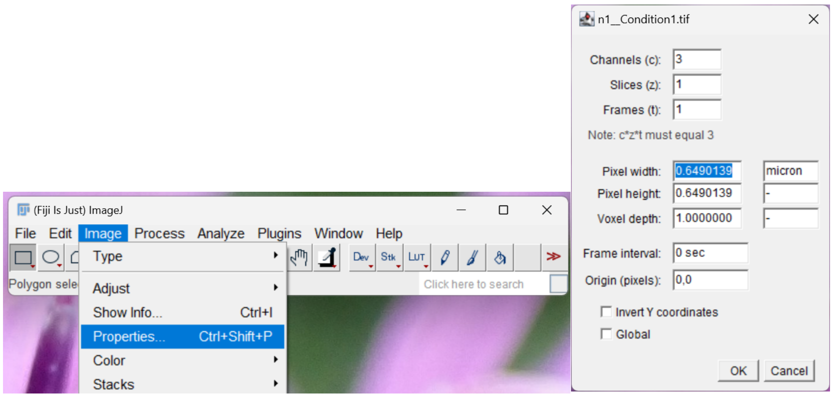


Report the pixel width for cooX and pixel height for cooY in the code:

donnees$cooX_px <- donnees$cooX/0.6490139

donnees$cooY_px <- donnees$cooY/0.6490139

If the cells appear too small or too big in the images displayed in the app, it is possible to adjust it. In chunk 6, server definition: functions output$zoomed_bf, output$zoomed_via and output$zoomed_cw. The functions are highlighted thanks to comments in the code.

##### ImageJ:

- **Check and update the IJPB plugin (see installation section of this user guide)**

#### Issues with the data:

##### Segmentation:

Improper segmentation can lead to incorrect intensity measurements like measuring adjacent cells signal or a smaller part of the cell.

Parameters can be adjusted depending on the microscopy images and the starting biological material (e.g.: size and flow threshold).

If using CellPose, help can be found here: https://cellpose.readthedocs.io/en/latest/faq.html

A model can also be trained to increase segmentation efficiency:
https://cellpose.readthedocs.io/en/latest/train.html

Other segmentation software or plugins can be used as the trainable Weka segmentation tool from ImageJ. https://imagej.net/plugins/tws/

##### Viability threshold:

Depending on the viability marker (FDA, other staining or presence of fluorescence in fluorescent cell line…) and the image acquisition, the way of determining the viability might not be optimal in the original code.

R script can be modified to use another automated thresholding method using the same autothreshold library:

https://www.rdocumentation.org/packages/autothresholdr/versions/1.4.2/topics/auto_thresh

The circularity cut off can also be changed to take more or less "round" objects: change 0.9 for another value in the following code line:

```
wdta$cellalive <- ifelse(wdta$meanViability > wdta$huang & wdta$Circularity > 0.9, "yes", "no")
```

*The code corresponding to the viability is at the end of chunk2.*

### Additional information

● The fulldata-saved file contains:

- med_rep: name of the medium/ condition followed by the replicate
- cell: replicate, medium/condition and cell label
- Medium: name of the medium/condition tested
- Replicate: name of the replicate or date of the experiment (what is before __ in the file naming)
- rep_medium: replicate and condition/medium
- diameter: calculated with the perimeter measured by the macro in the morphometry csv file
- Circularity: measured by the macro in the morphometry csv file
- meanCW: mean intensity of the cell wall measured by the macro in the CW-Intensity csv file
- meanViability: mean intensity of the cell wall measured by the macro in the Viability-Intensity csv file
- cooX: coordinate of the cell in the image on the X axis (used for the Q-Warg-shinyapp)
- cooY: coordinate of the cell in the image on the Y axis (used for the Q-Warg-shinyapp)
- huang: values of automated threshold for each medium/condition and replicate
- cellalive: yes or no (if no, the object might be a debris and not a cell)

● In R scripts, it is asked to select a file in the working directory. Other function(s) exist to directly select the folder but for an unknown reason, it might not work on all computers. As the choose.files() function is more robust, it is the one used here.

● In this user guide, the focus is made on the cell wall regeneration and CW staining intensity. However, the pipeline can be used to check other staining or fluorescence markers. For instance, it could be used to check the efficiency of transformation of fluorescent reporters. The CW channel will be replaced by a fluorescent marker. Be aware that all naming will still contain CW, but it will represent the chosen signal.

● All scripts are commented for easier comprehension and modifiability/flexibility.

### Bibliography


Barrett, T. *et al.* (2024) 'data.table: Extension of "data.frame"'. Available at: https://cran.r-project.org/web/packages/data.table/index.html (Accessed: 19 February 2025).

Chang, W. *et al.* (2024) 'shiny: Web Application Framework for R'. Available at: https://cran.r-project.org/web/packages/shiny/ (Accessed: 19 February 2025).

*Create Interactive Web Graphics via 'plotly.js' [R package plotly version 4.10.4]* (2024). Comprehensive R Archive Network (CRAN). Available at: https://CRAN.R-project.org/package=plotly (Accessed: 19 February 2025).

Garnier, S. *et al.* (2024) 'viridis: Colorblind-Friendly Color Maps for R'. Available at: https://cran.r-project.org/web/packages/viridis/index.html (Accessed: 19 February 2025).

Grosjean, P. (2014) 'tcltk2: Tcl/Tk Additions'. Available at: https://cran.r-project.org/web/packages/tcltk2/index.html (Accessed: 19 February 2025).

Mendiburu, F. de (2023) 'agricolae: Statistical Procedures for Agricultural Research'. Available at: https://cran.r-project.org/web/packages/agricolae/index.html (Accessed: 19 February 2025).

Nolan [aut, R. *et al.* (2023) 'autothresholdr: An R Port of the "ImageJ" Plugin "Auto Threshold"'. Available at: https://cran.r-project.org/web/packages/autothresholdr/index.html (Accessed: 19 February 2025).

Ooms [aut, J. and cre (2024) 'magick: Advanced Graphics and Image-Processing in R'. Available at: https://cran.r-project.org/web/packages/magick/index.html (Accessed: 19 February 2025).

Pedersen, T.L. (2024) 'patchwork: The Composer of Plots'. Available at: https://cran.r-project.org/web/packages/patchwork/index.html (Accessed: 19 February 2025).

Ren, K. (2021) 'rlist: A Toolbox for Non-Tabular Data Manipulation'. Available at: https://cran.r-project.org/web/packages/rlist/index.html (Accessed: 19 February 2025).

Rudis [aut, B. *et al.* (2024) 'hrbrthemes: Additional Themes, Theme Components and Utilities for "ggplot2"'. Available at: https://cran.r-project.org/web/packages/hrbrthemes/index.html (Accessed: 19 February 2025).

Sievert, C. *et al.* (2025) 'bslib: Custom "Bootstrap" "Sass" Themes for "shiny" and "rmarkdown"'. Available at: https://cran.r-project.org/web/packages/bslib/index.html (Accessed: 19 February 2025).

Vries, A. de *et al.* (2023) 'sortable: Drag-and-Drop in "shiny" Apps with "SortableJS"'. Available at: https://cran.r-project.org/web/packages/sortable/index.html (Accessed: 19 February 2025).

Wickham, H. *et al.* (2019) 'Welcome to the Tidyverse', *Journal of Open Source Software*, 4(43), p. 1686. Available at: https://doi.org/10.21105/joss.01686.
