## Supporting method S3 for "The Q-Warg Pipeline: A Robust and Versatile Workflow for Quantitative Analysis of Protoplast Culture Conditions"

///======================MACRO=========================///
macro_name = "CWRegenerationQuantification";
///====================================================///
///File author(s): Stephane Verger======================///

///====================Description=====================///
/* This macro takes as input multichannel .tif images of 
 * brightfield (channel 1),fluorescence for CW (C2) and 
 * Viability (C3), and corresponding cell label images (see
 * input parameters).
 * It outputs several morphometry measurments on the labels
 * for cell shape and size, and fluorescence intensity 
 * measurment for cell wall and viability stain under the 
 * form of .csv files. (see measurment and output parameters).
 * For further details see macro_source.
*/
macro_source = "https://github.com/...PATH_TO_DEFINE";//XXXXXXXXXXXXXXXXXXXXXXXXXXXXXXXXXXXXXXXXXXXXX

///=========Input/output file names parameters=========///
// Input paramaters: input files suffixes
MC_suffix = ".tif"; //Multichannel image containing Brightfield (C1)/CellWall stain (C2)/Viability stain (C3)
Lbl_suffix = "-labels.png"; //Label image from cellpose segmentation

// Measurment parameters (from MorpholibJ):
Int_Measurments = "mean stddev max min median mode skewness kurtosis numberofvoxels volume neighborsmean neighborsstddev neighborsmax neighborsmin neighborsmedian neighborsmode neighborsskewness neighborskurtosis";
Morpho_Measurments = "pixel_count area perimeter circularity euler_number bounding_box centroid equivalent_ellipse ellipse_elong. convexity max._feret oriented_box oriented_box_elong. geodesic tortuosity max._inscribed_disc average_thickness geodesic_elong.";

// Output paramaters: output file suffixes
BF_suffix = "--BF.tif"; //Brightfield image
CW_suffix = "--CW.tif"; //Cell wall stain image (e.g. Calcofluor)
Via_suffix = "--Viability.tif"; //Viability stain image (e.g FDA)
CWInt_suffix = "--CW--Intensity.csv"; //MorpholibJ fluorescence intensity measurment on the CW channel
ViaInt_suffix = "--Viability--Intensity.csv"; //MorpholibJ fluorescence intensity measurment on the viability channel
Morpho_suffix = "--Morphometry.csv"; //MorpholibJ "analyze regions" measurment on labels

// Create a dialog box with a dropdown menu for the user to choose the position of each channel
				Dialog.create("Channel Selection");
				Dialog.addChoice("C1 - ", newArray("BF", "CW", "Via"));
				Dialog.addChoice("C2 - ", newArray("BF", "CW", "Via"));
				Dialog.addChoice("C3 - ", newArray("BF", "CW", "Via"));
				Dialog.show();

				// Get the user's selections
				c1_selection = Dialog.getChoice();
				c2_selection = Dialog.getChoice();
				c3_selection = Dialog.getChoice();

///====================================================///
///====================================================///
///====================================================///

print("\\Clear");

//Select directory
dir = getDirectory("Choose a directory");
dir_name = File.getName(dir);
dir_list = getFileList(dir);

setBatchMode("hide");

//Generate log file for record
log_file_name = "Log_" + dir_name + "_" + macro_name + ".txt";
fLog = File.open(dir + File.separator + log_file_name);
print(fLog, "Files processed with the macro " + macro_name + ".ijm\n(" + macro_source + ")\n\n");
print(fLog, "Directory: " + dir + "\n\nFiles processed:");
File.close(fLog);

s = 0;
//Loop on all the folder in the directory
for (i=0; i<dir_list.length; i++){
	
	//Process folders only
	if (File.isDirectory(dir + dir_list[i])) {
		Folder_name = File.getName(dir + dir_list[i]);
		Folder_list = getFileList(dir + dir_list[i]);

		//Loop on all the images in the folder
		for (j=0; j<Folder_list.length; j++){
		
			//Select image series to process
			if(endsWith (Folder_list[j], Lbl_suffix)){
				print("file_path", dir + Folder_name + File.separator + Folder_list[j]);
		
				//count samples analyzed
				s++;
		
				//Extract generic name, path of the image serie
				File_name = substring(Folder_list[j], 0, indexOf(Folder_list[j], Lbl_suffix));
				Path = dir + File.separator + Folder_name + File.separator;
				
				//Define input and output filenames
				Lbl_Image = File_name + Lbl_suffix;
				MC_Image = File_name + MC_suffix;
				BF_Image = File_name + BF_suffix;
				CW_Image = File_name + CW_suffix;
				Via_Image = File_name + Via_suffix;
				Morpho_Results = File_name + Morpho_suffix;
				CW_Results = File_name + CWInt_suffix;
				Via_Results = File_name + ViaInt_suffix;
		
				//Write to log txt file
				File.append("- Sample number: " + s + "\n" + File_name, dir + File.separator + log_file_name);
				print("Sample: " + File_name);
				print("--> Opening input images");
				
				//Open multichannel image
				open(Path + MC_Image);
						    
			    // Split multichannel image and save/rename channels
				run("Split Channels");
				
				//Open label image
			    open(Path + Lbl_Image);
		    
		    	//Add segmentation contours from the label image
				selectImage(Lbl_Image);
				run("Duplicate...", "title=edge-Image.png duplicate");
				run("Find Edges");
				setMinAndMax(0, 1476);
				setAutoThreshold("Default dark no-reset");
				//run("Threshold...");
				setThreshold(1, 65535, "raw");
				run("Convert to Mask");
			
				
				// Save the channels based on the user's selections
				if (c1_selection == "BF") {
					selectImage("edge-Image.png");
					run("Yellow");
   					selectImage("C1-" + MC_Image);
   					saveAs("Tiff", Path + BF_Image);
   					run("Duplicate...", "title=C1-Image.tif duplicate");
   					run("Grays");
					run("Add Image...", "image=edge-Image.png x=0 y=0 opacity=100 zero");
    				saveAs("Jpeg", Path + BF_Image);
    				close();
				} else if (c1_selection == "CW") {
    				selectImage("edge-Image.png");
					run("Cyan");
    				selectImage("C1-" + MC_Image);
    				saveAs("Tiff", Path + CW_Image);
    				run("Duplicate...", "title=C1-Image.tif duplicate");
   					run("Grays");
					run("Add Image...", "image=edge-Image.png x=0 y=0 opacity=100 zero");
    				saveAs("Jpeg", Path + CW_Image);
    				close();
				} else if (c1_selection == "Via") {
					selectImage("edge-Image.png");
					run("Magenta");
    				selectImage("C1-" + MC_Image);
    				saveAs("Tiff", Path + Via_Image);
    				run("Duplicate...", "title=C1-Image.tif duplicate");
   					run("Grays");
					run("Add Image...", "image=edge-Image.png x=0 y=0 opacity=100 zero");
    				saveAs("Jpeg", Path + Via_Image);
    				close();
				}


				if (c2_selection == "BF") {
					selectImage("edge-Image.png");
					run("Yellow");
   					selectImage("C2-" + MC_Image);
   					saveAs("Tiff", Path + BF_Image);
   					run("Duplicate...", "title=C2-Image.tif duplicate");
   					run("Grays");
					run("Add Image...", "image=edge-Image.png x=0 y=0 opacity=100 zero");
    				saveAs("Jpeg", Path + BF_Image);
    				close();
				} else if (c2_selection == "CW") {
    				selectImage("edge-Image.png");
					run("Cyan");
    				selectImage("C2-" + MC_Image);
    				saveAs("Tiff", Path + CW_Image);
    				run("Duplicate...", "title=C2-Image.tif duplicate");
   					run("Grays");
					run("Add Image...", "image=edge-Image.png x=0 y=0 opacity=100 zero");
    				saveAs("Jpeg", Path + CW_Image);
    				close();    				
				} else if (c2_selection == "Via") {
					selectImage("edge-Image.png");
					run("Magenta");
    				selectImage("C2-" + MC_Image);
    				saveAs("Tiff", Path + Via_Image);
    				run("Duplicate...", "title=C2-Image.tif duplicate");
   					run("Grays");
					run("Add Image...", "image=edge-Image.png x=0 y=0 opacity=100 zero");
    				saveAs("Jpeg", Path + Via_Image);
    				close();
				}

				if (c3_selection == "BF") {
					selectImage("edge-Image.png");
					run("Yellow");
   					selectImage("C3-" + MC_Image);
   					saveAs("Tiff", Path + BF_Image);
   					run("Duplicate...", "title=C3-Image.tif duplicate");
   					run("Grays");
					run("Add Image...", "image=edge-Image.png x=0 y=0 opacity=100 zero");
    				saveAs("Jpeg", Path + BF_Image);
    				close();
				} else if (c3_selection == "CW") {
    				selectImage("edge-Image.png");
					run("Cyan");
    				selectImage("C3-" + MC_Image);
    				saveAs("Tiff", Path + CW_Image);
    				run("Duplicate...", "title=C3-Image.tif duplicate");
   					run("Grays");
					run("Add Image...", "image=edge-Image.png x=0 y=0 opacity=100 zero");
    				saveAs("Jpeg", Path + CW_Image);
    				close();
				} else if (c3_selection == "Via") {
					selectImage("edge-Image.png");
					run("Magenta");
    				selectImage("C3-" + MC_Image);
    				saveAs("Tiff", Path + Via_Image);
    				run("Duplicate...", "title=C3-Image.tif duplicate");
   					run("Grays");
					run("Add Image...", "image=edge-Image.png x=0 y=0 opacity=100 zero");
    				saveAs("Jpeg", Path + Via_Image);
    				close();
				}

			    selectImage("edge-Image.png");
			    close();
			    
			    //Set pixel size for label image based on BF image
				selectImage(BF_Image);
				getPixelSize(unit, pixelWidth, pixelHeight);
				selectImage(Lbl_Image);
				setVoxelSize(pixelWidth, pixelWidth, "1", unit);
				
			    //Quantifications
			    print("--> Measurments:");
		
				// Measure, save and close cell moprhometrics
				print("    --> Morphometry");
				selectImage(Lbl_Image);
				run("Analyze Regions", Morpho_Measurments);
				saveAs("Results", Path + Morpho_Results);
				selectWindow(Morpho_Results);
				run("Close");
		
				//Measure, save and close CW intensity
				print("    --> CW intensity");
				run("Intensity Measurements 2D/3D", "input=" + CW_Image + " labels=" + Lbl_Image + " " + Int_Measurments);
				saveAs("Results", Path + CW_Results);
				selectWindow(CW_Results);
				run("Close");
		
				//Measure, save and close cell viability
				print("    --> Viability\n");
				run("Intensity Measurements 2D/3D", "input=" + Via_Image + " labels=" + Lbl_Image + " " + Int_Measurments);
				saveAs("Results", Path + Via_Results);
				selectWindow(Via_Results);
				run("Close");
				
	
				//Close images
				close("*");
		
				//Write to log, input files used, output file generated, time and date
				File.append("\tInput :\n\t=> " + MC_Image + "\n\t=> " + Lbl_Image + "\n\tOutput :\n\t<= " + Morpho_Results + "\n\t<= " + CW_Results + "\n\t<= " + Via_Results, dir + File.separator + log_file_name);
				getDateAndTime(year, month, dayOfWeek, dayOfMonth, hour, minute, second, msec);
				File.append("\t" + hour + ":" + minute + ":" + second + " " + dayOfMonth + "/" + month + "/" + year + "\n\n", dir + File.separator + log_file_name);
			}
		}
	}
}

setBatchMode("exit and display");

//End of the macro message
print("\n\n===>End of the " + macro_name + " macro");
print("Check output files in:\n" + dir);
print("- " + log_file_name + "\n- *" + Morpho_suffix + "\n- *" + CWInt_suffix + "\n- *" + ViaInt_suffix + "\n(*) For each image analyzed");
